# Adaptive Circuit Dynamics Across Human Cortex During Evidence Accumulation in Changing Environments

**DOI:** 10.1101/2020.01.29.924795

**Authors:** Peter R Murphy, Niklas Wilming, Diana C Hernandez-Bocanegra, Genis Prat Ortega, Tobias H Donner

## Abstract

Many decisions under uncertainty entail the temporal accumulation of evidence that informs about the state of the environment. When environments are subject to hidden changes in their state, maximizing accuracy and reward requires non-linear accumulation of the evidence. How this adaptive, non-linear computation is realized in the brain is unknown. We analyzed human behavior and cortical population activity (measured with magnetoencephalography) recorded during visual evidence accumulation in a changing environment. Behavior and decision-related activity in cortical regions involved in action planning exhibited hallmarks of adaptive evidence accumulation, which could also be implemented by a recurrent cortical microcircuit. Decision dynamics in action-encoding parietal and frontal regions were mirrored in a frequency-specific modulation of the state of visual cortex that depended on pupil-linked arousal and the expected probability of change. These findings link normative decision computations to recurrent cortical circuit dynamics and highlight the adaptive nature of decision-related feedback to sensory cortex.

## Introduction

Many decisions under uncertainty entail the temporal accumulation of noisy information about the state of the world. When the source of the evidence remains constant during decision formation, maximizing decision accuracy requires perfect, linear integration (i.e., equal weight given to each sample of evidence)^1,2^. Progress in computational theory, behavioral modeling, and neurophysiological analysis has translated these ideas into an influential framework wherein decision-relevant evidence is encoded by neurons in sensory cortical regions, and then accumulated over time along sensory-motor cortical pathways^2–4^. This computation can account for the build-up of choice-predictive activity that has been observed in several brain regions during decision formation^5–14^.

Despite its theoretical appeal and popularity, this framework has two fundamental limitations. First, its development has been informed by studying a single, artificial behavioral context: environments in which the source (i.e. stimulus category) generating evidence remains constant during decision formation. In this special case, the agent’s uncertainty originates only from noise corrupting the evidence. However, natural environments can undergo hidden changes in their state, constituting an additional source of uncertainty^15^. Perfect and linear evidence accumulation is suboptimal for changing environments^16,17^. Instead, optimal decision-making in such contexts requires a non-linear evidence accumulation process that *adapts* to the statistics of the environment and, in doing so, appropriately balances stability of decision formation with sensitivity to change^16^. Second, the linear, feedforward integration of evidence in sensory-motor cortical pathways entailed in the above framework contrasts with the recurrent organization and resulting non-linear dynamics of cortical circuits^18–20^.

Here, we set out to address these two limitations which, we reasoned, may be closely related. The goal of the study was twofold. First, we aimed to identify neural signatures of an (approximately) normative decision variable during evidence accumulation in a changing environment, across the cortical pathway transforming sensory input into behavioral choice. Second, we aimed to uncover candidate circuit mechanisms for generating these neural signals.

While human participants accumulated samples of visual evidence in a changing environment, signatures diagnostic of the adaptive, non-linear computation were evident in their behavior and in choice-predictive population dynamics (measured with magnetoencephalography, MEG) of cortical regions involved in action planning. The same behavioral and neural signatures were produced by recurrent synaptic interactions in a biophysically detailed model of a cortical decision circuit. Going further, we also uncovered aspects of cortical dynamics that were not explained by interactions within cortical decision circuits alone: A frequency-specific (alpha-band) component of visual cortical activity was selectively shaped by (i) the decision variable evolving downstream and (ii) phasic arousal signals elicited by evidence samples indicative of change-points. The feedback component was most strongly expressed in early visual cortex (V1) and adapted to the level of environmental volatility, in line with a context-dependent, stabilizing signal.

## Results

To interrogate the mechanisms of adaptive perceptual evidence accumulation, we developed a task with hidden changes in the environmental state (i.e. evidence source; Figure 1a). The evidence samples were small checkerboard patches presented in a rapid sequence at different locations along a semicircle in the lower visual hemifield (see *Methods* for details). Sample positions were generated from one of two noisy sources: the probability distributions shown in the upper row of Figure 1a. The ‘active’ source was chosen at random at the beginning of each trial and could change at any time during the trial, with low probability (hazard rate *H),* which was set to 0.08 in the main experiment. The participant’s task was to report which source was active *at the end* of each sequence via left- or right-handed button presses.

**Figure 1.**
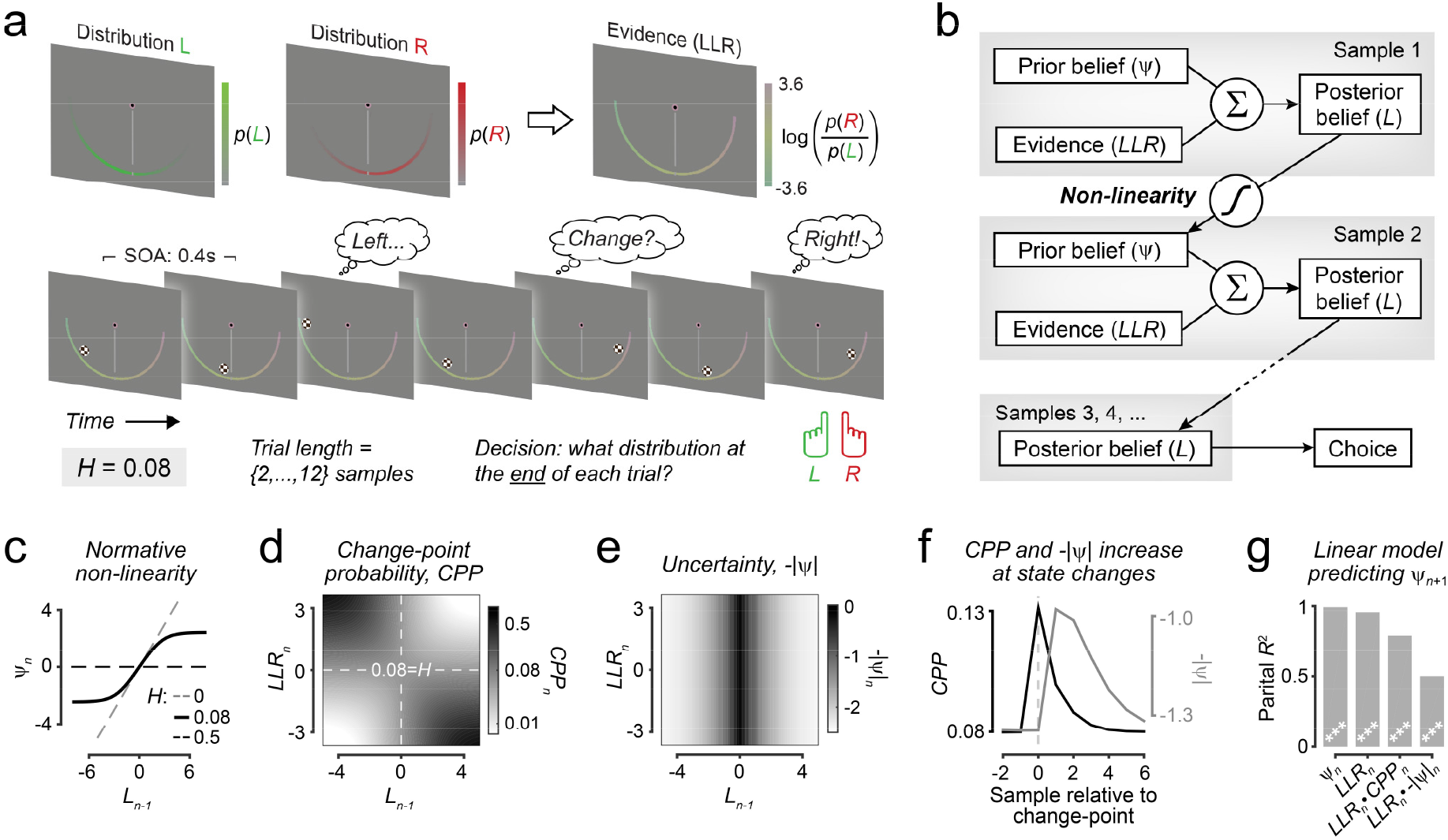
Behavioral task and normative model with diagnostic signatures of non-linear evidence accumulation. (**a**) Schematic of two-alternative perceptual choice task with hidden state changes, including an example sample sequence (bottom; fixation point, vertical reference line and semicircle reflecting evidence strength all visible to participants). (**b**) Schematic of normative evidence accumulation process. (**c**) Non-linearity in normative model for hazard rate used in choice task (H=0.08), and extremes of perfectly stable (H=0; identity line) and unpredictable (H=0.5, flat line) environments. (**d,e**) Change-point probability (CPP; d) and uncertainty (-IψI; e) as function of posterior belief after previous sample (Ln-1) and evidence provided by current sample (LLRn). (**f**) CPP and –IψI dynamics centered on change-points in generative task state. (**g**) Coefficients of partial determination reflecting contribution of key computational variables to belief updating. ***p<10^-6^, two-tailed t-test. Panels d-g based on generative statistics of behavioral task (see main text).

### Signatures of adaptive evidence accumulation in human behavior

The normative model maximizing accuracy on this task (Figure 1b) entails the accumulation of evidence samples that carry information (in the form of log-likelihood ratios, *LLR)* about the two possible environmental states^16^ – a process also known as ‘belief updating’. This model serves as a benchmark against which the behavior and neural activity of human participants can be compared. The key difference between the normative model and previous accumulator models^1^ is the dynamic shaping of the evidence accumulation (the combination of prior belief *ψ* with new evidence *LLR* to yield posterior belief *L*) by the estimated rate of environmental state change (*H*)^16^. Specifically, the prior for the next sample (*ψ*_*n*+1_) is determined by passing the updated belief (*L_n_*) through a non-linear function that, depending on *H*, can saturate (slope≈0) for strong *L_n_* and entail more moderate information loss (0<<slope<1) for weak *L_n_* (Figure 1c; Supplementary Figure 1). By this process, the normative model strikes an optimal balance between formation of strong beliefs in stable environments versus fast change detection in volatile environments.

In our setting, this non-linearity could be cast in terms of sensitivities of evidence accumulation to two quantities: (i) the uncertainty about the environmental state before encountering a new sample (denoted as –*IψI;* Figure 1e) and (ii) the change-point probability *(CPP;* Figure 1d) defined as posterior probability that a state change just occurred, given an existing belief and new evidence sample. *CPP* scaled with the inconsistency between the new sample and existing belief (Figure 1d). Neither variable was explicitly used in the normative computation, but both helped pinpoint diagnostic signatures of adaptive evidence accumulation in behavior and neural activity, as shown below. Across levels of *H* and environmental noise, *CPP* increased transiently when a state change occurred followed by a period of heightened *-IψI* (Figure 1f; Supplementary Figure 1). A linear model in which both quantities modulated belief updating reliably predicted the extent to which an ideal observer integrated a new sample of evidence to form the prior belief for the next sample (**Model 1**, *Methods;* Figure 1g; Supplementary Figure 1; median *R*^2^ across generative settings = 99.7%, range = [89.2% 99.98%]).

Participants performed the task better than expected from two simple strategies (Figure 2a, empty bars; Supplementary Figure 2): deciding based on the sum of all samples (73.9 % ± 1.2 %; *p*<0.0001, two-tailed permutation test; Figure 2a, magenta) or on the final sample only (71.8 % ± 0.6 %; *p*<0.0001, two-tailed permutation test); Figure 2a, gray). Participants’ performance (77.5 % ± 1.9 %; mean and s.e.m.) came close to the benchmark performance of the ideal observer with perfect knowledge of *H* and the generative distributions, and without noise or biases (81.8 % ± 0.9 % (mean and s.e.m. across participant-specific stimulus sequences; Fig. 2a, navy bar).

**Figure 2.**
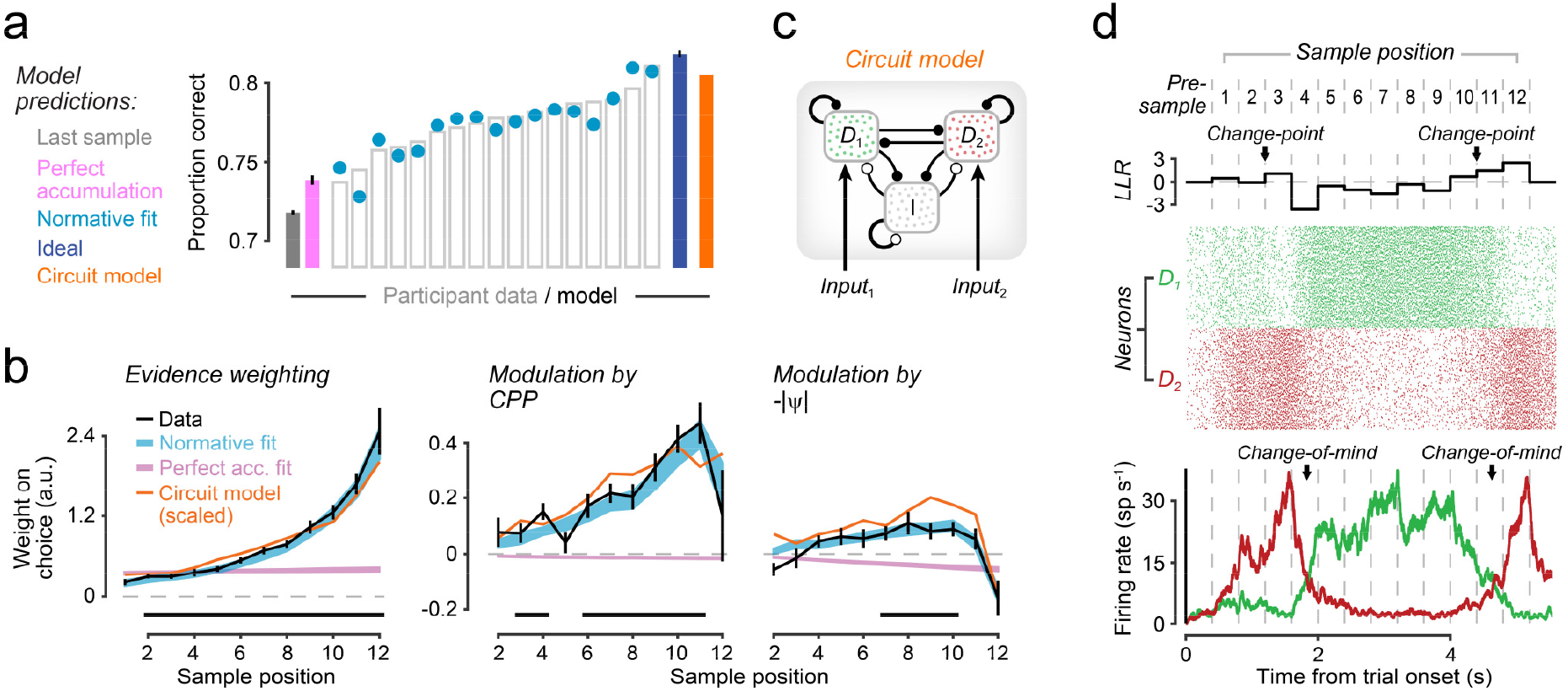
Signatures of adaptive evidence accumulation in human behavior and cortical circuit model. (**a**) Performance of human participants (unfilled bars; N=17), ideal observer (navy), cortical circuit model (orange), and perfect accumulation (magenta) or deciding based on last sample only (gray). Cyan dots, accuracies of normative model fits. (**b**) Time-resolved weight of evidence on choice (left), and its modulation by CPP (middle) and -IψI (right). Black, participants’ data; significance bars, time points where weights differ from zero (p<0.05, two-tailed cluster-based permutation test). Orange, kernels produced by circuit model, scaled to align with kernel magnitudes from participants. Cyan and magenta shadings, fits of normative and perfect accumulator models, respectively. (**c**) Schematic of circuit model. D_1_ (left choice, green) and D_2_ (right choice, red), choice-selective populations of excitatory neurons; I, shared population of inhibitory interneurons. (**d**) Response of choice-selective populations to an example trial. Top, LLRs; middle, raster plot of spiking activity from D_1_ and D_2_ neurons; bottom, population firing rates. Arrows in top and bottom highlight changes in environmental state and changes-of-mind, respectively.

Critically, our participants exhibited the same modulation of evidence accumulation by *CPP* and *-IψI* as the normative process depicted in Figure 1. We quantified this modulation through logistic regression (**Model 2**, *Methods).* The first set of regression weights, which quantified the time course of the leverage of *LLR* on choice (also called psychophysical kernel^12,21^) increased over time (*p*<0.0001, two-tailed permutation test on weights for first 6 samples *vs*. last 6 samples; Figure 2b, *left).* Thus, like the normative model for our task (Supplementary Figure 1d), participants tended to discount early information in their final choices. The second and third sets of regression weights captured the modulation of evidence weighting by *CPP* and *-IψI.* Each revealed strong positive modulations *(CPP: p*<0.0001, Figure 2b, *middle; -IψI: p*=0.0002, Figure 2b, *right;* two-tailed cluster-based permutation tests). The modulatory effect of *CPP* was also larger than that of *-IψI* (*p*<0.0001, two-tailed permutation test averaging over sample position). In other words, both *CPP* and *-IψI* ‘up-weighted’ the impact of the associated evidence on choice. The same evidence weighting signatures were produced by the normative accumulation scheme for a range of task statistics including the present one (Supplementary Figure 1d). The signatures also replicated in a separate dataset at faster and slower sample presentation rates (Supplementary Figure 3).

These behavioral signatures ruled out a range of alternative evidence accumulation schemes. Perfect linear accumulation (without bounds) produces flat psychophysical kernels (Figure 2b, magenta) while perfect accumulation toward an absorbing bound produces stronger weights for early samples^22^. Leaky accumulation with exponential decay of accumulated evidence (refs. ^23^) produces recency but not the *CPP* or -*IψI* modulations of evidence weighting (Supplementary Figure 4). In sum, the signatures in Figure 2b suggested that participants’ behavior used a computation that approximated the normative evidence accumulation process shown in Figure 1b.

This conclusion was supported by fitting different models to participants’ individual choices (Figure 2 and Supplementary Figures 4, 5). We fitted several variants of the normative model, with between 3 and 11 free parameters (see *Methods* for details). Here, we use the term ‘normative model’ as shorthand for the adaptive accumulation scheme from Figure 1b, without implying a noise-free or unbiased computation. All variants included a free subjective hazard rate parameter (*H*), a decision noise parameter and at least one parameter for assigning subjective weight (*LLR*) to the range of possible sample locations^16^. Some variants also included up to seven additional parameters for a non-linear scaling of subjective *LLR*, and/or a gain parameter controlling the evidence weighting depending on its consistency with current belief^16,24–27^. Inclusion of these additional parameters was supported by model comparison (Supplementary Figure 5; *Methods).* The choices of the best-fitting model variant (‘normative fit’ in Fig. 2, cyan) were highly consistent with participants’ choices (88.8 ± 0.8%). The normative fit outperformed all considered versions of perfect and leaky accumulator models (mean Δ Bayes Information Criteria (BIC) = 71.1 ± 7.6; higher BIC for all 17 participants; Supplementary Figure 5).

One alternative to the normative model, perfect accumulation toward non-absorbing bounds, did produce the above evidence weighting signatures (Supplementary Figure 4) and only marginally worse fits to the data than the best-fitting version of the normative model (ΔBIC = 11.1 ± 2.9; higher BIC for 16 of 17 participants; Figure 2c). Indeed, the non-linearity imposed by the non-absorbing bounds can approximate the normative accumulation scheme for our task setting (low *H* and high SNR)^16^. This indicates that saturation for strong belief states (and not leak for weak belief states) was critical in accounting for participants’ behavior.

Taken together, our analyses indicate that, while limited in their performance by internal noise and biased internal representation of task variables, participants approximated the normative evidence accumulation process for the current setting: adaptive, non-linear evidence accumulation, characterized by sensitivity during periods of uncertainty and plateaus in the evolving decision variable during periods of relative stability.

### Recurrent cortical circuit dynamics can implement adaptive evidence accumulation

A prominent cortical circuit model for decision-making^20^ is made up of recurrently connected spiking neurons (Figure 2c). The circuit model has previously been used to simulate single-neuron activity in regions implicated in decision-making during standard perceptual choice tasks^20^. This model precludes perfect accumulation, which limits its performance in standard perceptual choice tasks without change-points. Instead, it exhibits sensitivity to input early in a trial and stable, saturating firing rates (so-called attractor states) during periods of stability (Figure 2d). We therefore reasoned that this model might also exhibit the above-described features of the normative accumulation scheme, its approximation in our generative setting (accumulation toward non-absorbing bounds), and human behavior.

Indeed, when simulated for our task, the circuit model consistently ‘changed its mind’ (i.e. flipped the sign of activity dominance between choice-selective populations) in response to changes in the evidence source (Figure 2d). Even without quantitative fitting, the circuit model’s behavior approximated the performance of the normative model and human participants (Figure 2a), and its choices were highly consistent with both the normative model fits (91.8 ± 0.5%) and the humans participants (86.2 ± 0.7%). Likewise, the single-trial trajectories of the decision variables from normative and circuit models were strongly correlated (median *r*=0.90 ± 0.009, *p*<0.0001, two-tailed permutation test).

Most importantly, the model naturally generated all behavioral signatures of the normative computation – specifically, the modulation of evidence weighting by *CPP* and *–IψI* (Figure 2b, orange; see Supplementary Figure 6 for an assessment of the boundary conditions for this behavior). These findings suggest that the non-linear dynamics of the recurrent circuit are, in fact, adaptive for decision-making in changing environments.

### Motor preparatory cortical activity tracks adaptive evidence accumulation

The above behavioral analyses indicate that participants (and the circuit model) implemented an adaptive evidence accumulation scheme in line with normative principles. We next sought to identify signatures of the resulting decision variable in cortical population activity. Previous work on choice tasks without change-points has identified signatures of evidence accumulation in neural signals reflecting the preparation of an action plan^2,4,6–9,12,13,28,29^. To isolate this motor preparatory activity, we applied a linear (spectral and spatial) filter to the data from the main task *(Methods;* Figure 3a, right panel). The filter was constructed using data from a separate delayed motor response task in which participants prepared left- or righthanded button presses, and which yielded sustained lateralization of alpha (8-14 Hz) and beta (16-30 Hz) band power over motor cortices contralateral to the prepared response (Figure 3a, left).

**Figure 3.**
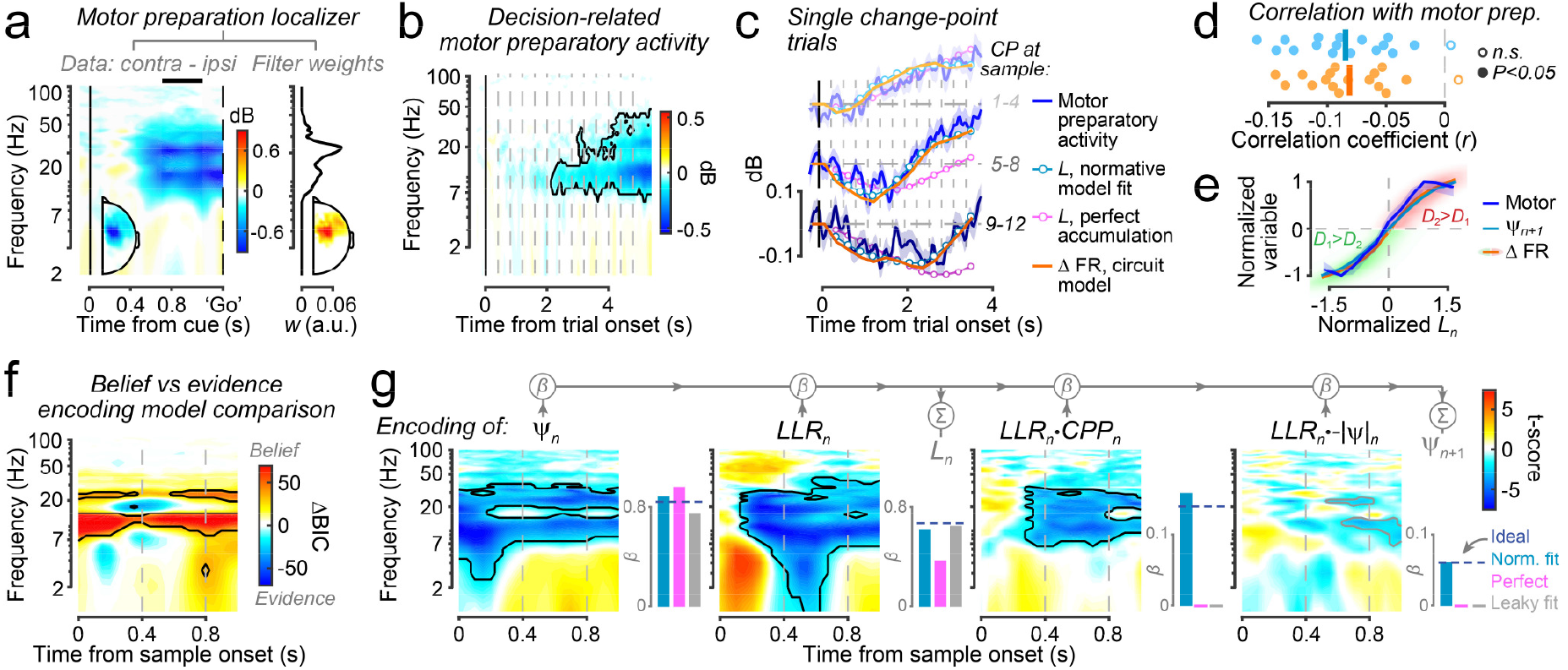
Motor preparatory activity tracks normative and circuit model decision variables. (**a**) Left: Timefrequency maps of MEG power lateralization contra vs. ipsilateral to response during localizer task. Black bar, interval for derivation of filter weights shown on right. See Methods for details. (**b**) Motor preparatory activity during decision-making task. Dashed vertical lines, sample onsets; contour, p<0.05 (two-tailed cluster-based permutation test). (**c**) Motor preparatory activity (lines and shaded areas, mean±s.e.m.) for groups of trials with single change points at different latencies. Cyan, orange and magenta: decision variables from normative fits, circuit model, and perfect accumulator, respectively. Signs of all variables are conditioned on generative state at trial end, and decision variables are vertically scaled and time-shifted by same factors across trial groups. (**d**) Correlation coefficients of single-trial relationship between motor preparatory activity and decision variable trajectories from normative (cyan) and circuit (orange) models. Points, individual participants; lines, group means. (**e**) Relationships of normative Ln with motor preparatory activity (blue; shaded area, bootstrap 95% CI) and circuit model decision variable (orange). Red and green shading, Gaussian kernel density estimation of distribution. Cyan, non-linearly transformed decision variable (ψ_n+1_) from normative fits, using same binning. (**f**) Comparison of pure evidence (LLRn) versus non-linearly transformed decision variable (ψ_n+1_) encoding model fit to motor preparatory activity. Contour, ΔBIC=±35. (**g**) Single-trial encoding of model variables in motor preparatory activity. Colors, group-level t-scores. Black contours, p<0.05 (two-tailed cluster-based permutation test); gray contours, largest sub-threshold clusters (p<0.05, two-tailed t-test, uncorrected). Insets, regression coefficients from linear model predicting belief updating for different accumulation schemes. Only the normative model yields non-zero coefficients for all terms.

The selective motor preparatory activity built up gradually throughout the prolonged period of evidence accumulation (Figure 3b). All analyses pooled across correct and error trials because, unlike in tasks without change-points, errors do not necessarily imply a failure of inference: Even the ideal observer performing our task made errors on ~18% of trials, despite the absence of noise and biases, and the inference process being exactly the same as on correct trials (Figure 2a, navy).

The build-up of motor preparatory activity closely mirrored the trajectories of the decision variables from the normative and circuit models, but *not* from the perfect accumulator, which failed to rapidly respond to change points (Figure 3c,d). The single-trial activity was significantly correlated with the decision variable trajectories of both the normative fit and the circuit model across the group (Figure 3d; normative: mean *r*=-0.085, *p*<0.0001, two-tailed permutation test; circuit: mean r=-0.081, *p*<0.0001, two-tailed permutation test), and individually within 16 of 17 participants (Fig. 3d, filled circles). Also, motor preparatory activity after each updating step (i.e., after computation of *ψ_n+1_,* the prior for next sample) showed a sigmoidal dependence on *L_n_* just as predicted by both the normative and circuit models (Figure 3e).

To quantify the adaptive nature of the dynamics of motor preparatory activity following each sample, we regressed it on a linear model made up of prior belief (*ψ_n_*), new evidence (*LLR_n_*), and the interactions of new evidence with *CPP_n_* and -*IψI_n_* (*Methods*: **Model 3**). While the two interaction terms would be zero for linear (perfect or leaky) accumulation schemes (Figure 3g, pink/gray bars), a signal reflecting the normative decision variable should have non-zero regression coefficients for all terms, with a stronger interaction term for *CPP_n_* than -*IψI_n_* (Figure 3g, cyan).

This is what we found for the motor preparatory activity (Figure 3g). The first three terms were encoded in motor preparatory activity in both beta- and alpha-bands (*ψ*: *p*=0.0006; *LLR*: *p*=0.0002; *LLR_n_*CPP_n_*: *p*=0.0009; two-tailed cluster-based permutation tests) with sustained encoding of the prior, and delayed encoding of the evidence sample that was in turn modulated by *CPP.* The *-IψI_n_* interaction was weaker than the *CPP_n_* interaction as expected (cyan bars for normative fits), showing only a trend in the expected direction (*p*=0.08, cluster-corrected; Figure 3g). Formal comparison between ‘versions of the above regression model containing only evidence encoding or only belief encoding *(Methods:* **Models 4-5**) showed that the motor preparatory alpha-/beta-band activity was better explained by encoding of the non-linearly updated decision variable *(ψ_n+1_*) than encoding of momentary evidence *(LLR_n_*; Figure 3f, group-level BIC scores; BIC_dv_<BIC_evidence_ for 15 of 17 participants averaged over highlighted clusters). We also note that *LLR* was weakly positively encoded in transient low-frequency delta-/theta-band activity (1-7 Hz; figure 3g, middle left). However, this effect did not survive correction for multiple comparisons (*p*=0.11), possibly because the cluster was small in spatiotemporal extent.

In sum, the dynamics of population activity in the human motor cortical system approximated the dynamics of the normative decision variable for our change-point task. These adaptive cortical dynamics (Figure 3) and the associated behavioral signatures (Figure 2) naturally emerged from the cortical circuit model. This alignment between circuit model, normative model and motor preparatory activity was remarkable, because we did not quantitatively fit the circuit model to any feature of the data. Yet, the transformation of sensory input into behavioral choice unfolds in a pathway made up of many, recurrently connected cortical regions spanning sensory to motor cortex, each of which have different properties (e.g. strength of self-recurrence)^30,31^. In perceptual choice tasks without change-points signatures of the decision variable (action plan) have been observed in all these areas, including even early sensory cortex^14,21,32^. Furthermore, brainstem arousal systems exert powerful effects on cortical circuit dynamics during such tasks^33^. These considerations motivated a set of further analyses described in the following sections. These aimed to (i) elucidate the evolution of the decision variable across the sensory-motor pathway for our change-point task, from primary visual cortex (V1) to primary motor cortex (M1), and (ii) quantify the arousal modulation of the decision computation across this pathway.

### Visual cortical encoding of evidence and decision variable in distinct frequency bands

We reconstructed the dynamics of selective activity lateralization in set of regions of interest (ROIs) along the sensory-motor pathway *(Methods):* multiple visual field maps in occipital, parietal, and temporal cortex as well as parietal and frontal regions encoding left-vs. righthand movements^32^. In all ‘action-related’ cortical regions except aIPS, the lateralization of alpha- and beta-band activity exhibited sustained encoding of *LLR* (Figure 4a) and prior belief (*ψ*), as well as modulation of *LLR* encoding by *CPP* (Figure 4b). This is consistent with the sensor-level results from Figure 3 and further supports the representation of the decision variable in the format of the associated action plan.

**Figure 4.**
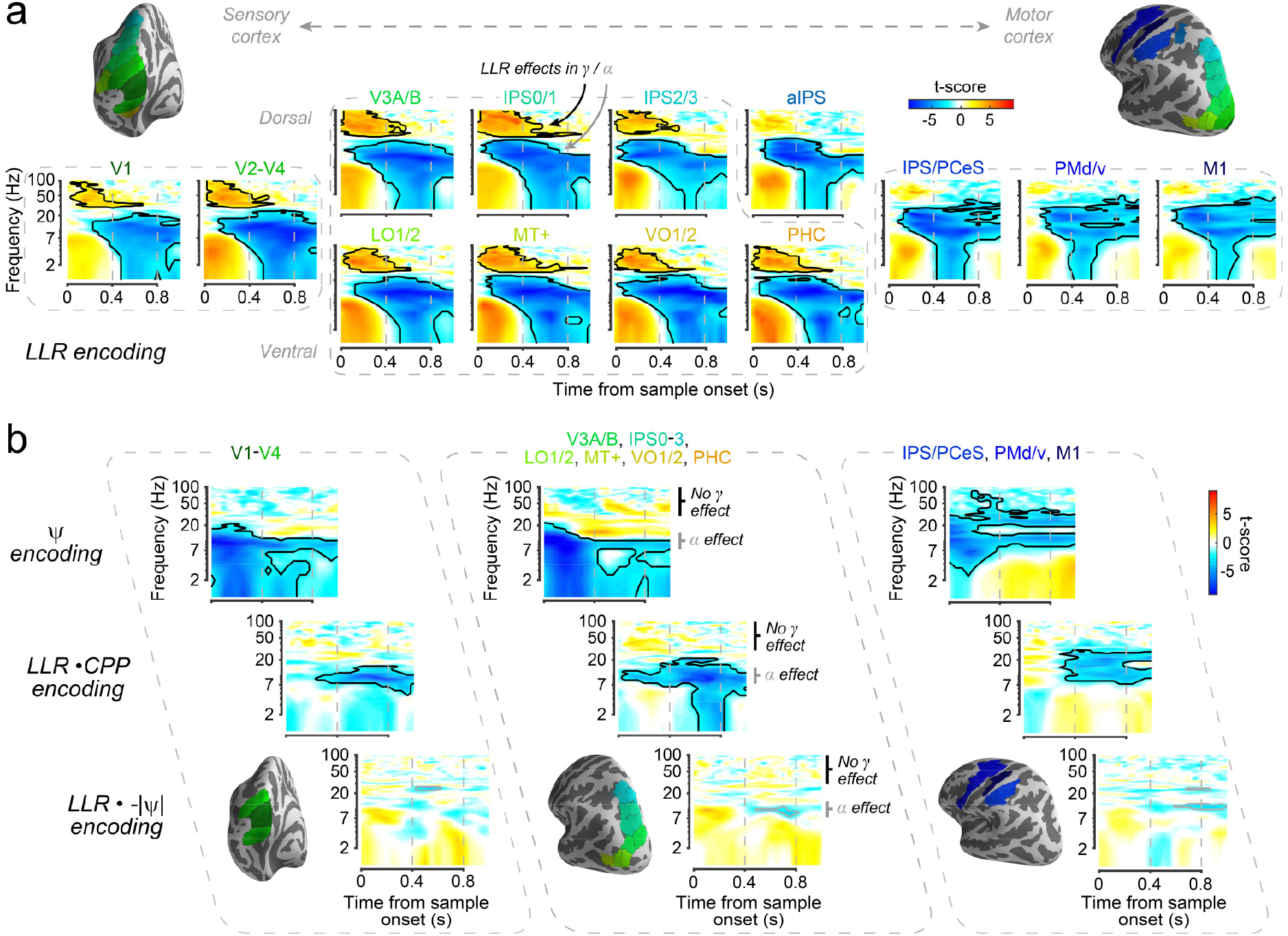
Frequency-specific encoding of evidence and decision variable along visuo-motor cortical pathway. (**a**) Time-frequency maps of evidence (LLR) encoding across anatomical ROIs in visual, temporal, posterior parietal and motor/premotor cortices (insets). Colors, group-level t-scores. Note positive LLR encoding in gamma-band (40-65 Hz; black arrow for IPS0/1) in all visual field maps but absent in aIPS, IPS/PCeS, PMd/v, M1, and negative LLR encoding in alpha-band (8-14 Hz; gray arrow for IPS0/1) in all ROIs. aIPS lacks both transient gamma-band response of posterior ROIs and sustained beta-band (16-30 Hz) response of anterior ROIs, and was omitted from further clustering. (**b**) Time-frequency maps reflecting encoding of other constituent components of adaptive evidence accumulation (same as Figure 3g), averaged for compactness across three sets of ROIs (insets) encompassing occipital visual field maps (V1-V4), temporal and parietal visual field maps (LO1/2, MT+, VO1/2, PHC, V3A/B, IPS0-3), and anterior intraparietal / motor cortices (IPS/PCeS, PMd/v, M1).. Black contours, p<0.05 (two-tailed cluster-based permutation test); gray contours, largest sub-threshold clusters of LLR --\ψ\ effect, (p<0.05, two-tailed t-test).

In the visual field maps, the encoding of computational variables exhibited different features. First, the lateralization of short-latency gamma-band (40-65 Hz) responses transiently encoded *LLR* (Figure 4a), but none of the other variables (Figure 4b) – a profile consistent with pure evidence encoding. By contrast, alpha-band lateralization in visual cortex mirrored the profile of alpha/beta lateralization in downstream action-related regions, in particular the sustained encoding of prior belief (*ψ*) and an interaction between *LLR* and *CPP* (Figure 4). Thus, alpha-band activity even in early visual cortex (V1-V4) was also consistent with encoding of the normative decision variable.

Overall, alpha-band activity in visual cortex was reminiscent of feedback of the evolving decision variable identified in earlier work on standard perceptual choice tasks (i.e., without change-points)^19,21,32^, as well as the idea that visual cortical alpha-band activity reflects cortical feedback signaling^32,34,35^. We further explored this idea in a series of analyses focusing on a hierarchically ordered set of dorsal visual field maps^36^ (V1, V2-V4, V3A/B, IPS0/1, IPS2/3) as described in the next section.

### Signatures of decision feedback in early visual cortex adapt to environmental volatility

We first delineated the feedforward processing of sensory evidence by means of multi-variate decoding of the spatial patterns of evoked responses^27,37^, separately for each ROI (Figure 5a-d). The *LLR* decoding latency of action-related regions (termed ‘motor’ in Figure 4) lagged that of V1 (*p*=0.005, two-tailed weighted permutation tests, *Methods).* The timescale of *LLR* decoding increased from V1 to extrastriate (*p*=0.048) and motor areas (*p*=0.007; extrastriate vs. motor: *p*=0.021, two-tailed weighted permutation tests, Figure 5d). Within the visual cortical hierarchy, timescales increased monotonically (*p*=0.017, two-tailed weighted permutation test on slope of regression line across areas; Figure 5c). The increase of evidence decoding timescales across the visual cortical hierarchy was mirrored by progressively slow intrinsic activity fluctuations during the pre-trial interval (Supplementary Figure 7), in line with monkey work^30^. Taken together, these results indicate feedforward processing of evidence from V1 to higher-tier visual to motor regions, in line with previous work^27,37^.

**Figure 5.**
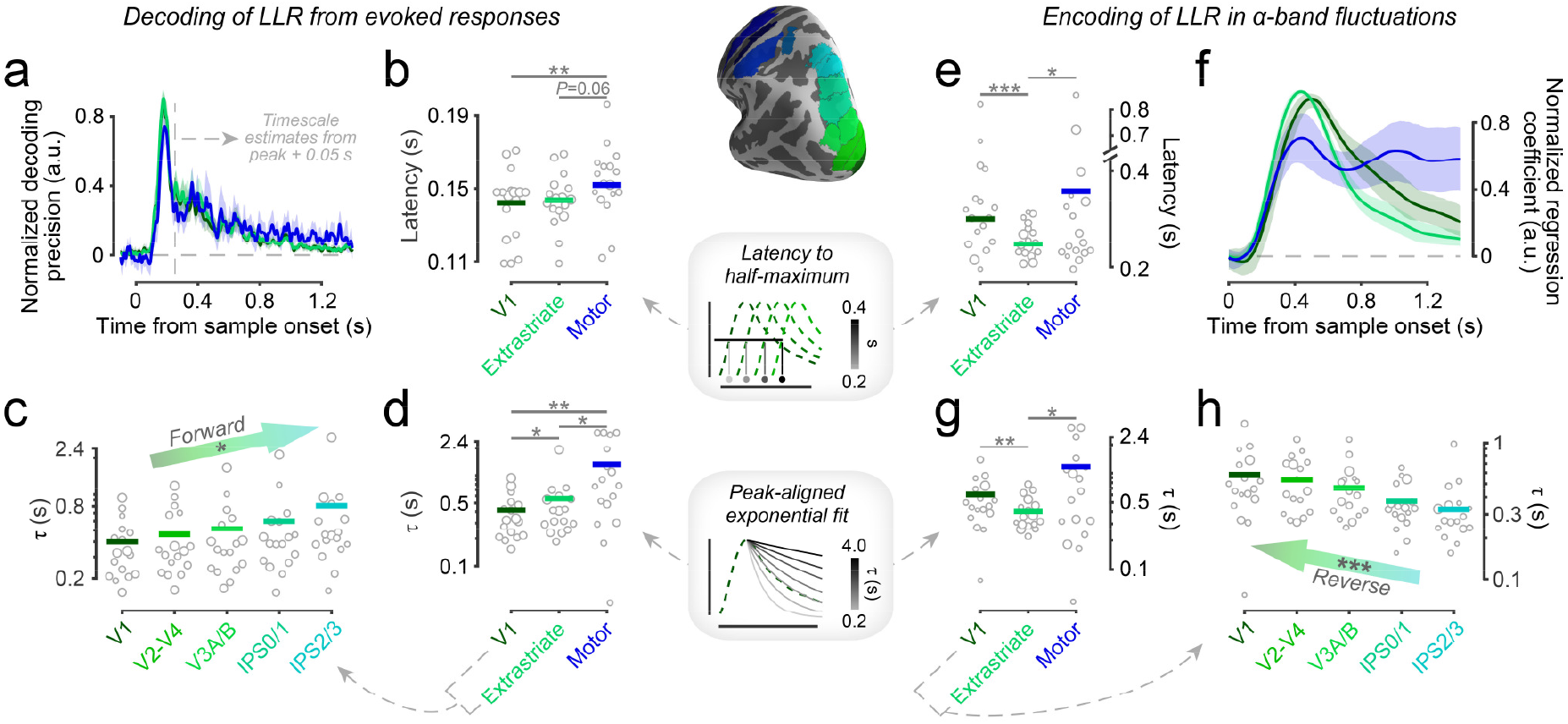
Properties of decision signals across visual cortical hierarchy. (**a**) Time-courses of vertex-based decoding of single-sample LLRs from evoked MEG responses, averaged within 3 ROI groups: V1, extrastriate visual (V2-IPS3 pooled), and motor (IPS/PCeS, PMd/v, M1 pooled). Decoding precision (cross-validated Pearson correlations) normalized per subject to peak at 1 (Methods). Lines, weighted means; shaded areas, bootstrap 95% confidence intervals. (**b**) Latencies to half maximum of LLR decoding traces for same ROI clusters. Lines, weighted means; dots, individual participants with dot size proportional to weight determined by participant-specific LLR decoding precision. (**c**) Timescales of decaying exponentials fit to LLR decoding traces for a dorsal set of visual cortical areas. Same format as in b. (**d**) Timescales averaged within the same ROI clusters as in b. (**e-h**) Same format as a-d, but for LLR encoding selectively in the 8-14 Hz lateralization signal. *p<0.05, **p<0.01, ***p<0.001, two-tailed weighted permutation test; ‘forward’ and ‘reverse’ labels indicate direction of hierarchy inferred from slopes of parameters across ROIs.

The feedforward scheme suggested by Figure 5a-d and Supplementary Figure 7 stood in sharp contrast to the signatures of *LLR* encoding in alpha-band lateralization (Figure 5e-h). Alpha-band *LLR* encoding latency of V1 lagged that of extrastriate (*p*<0.0001, two-tailed weighted permutation test) but not of motor regions (*p*=0.11; Figure 5e), and the *LLR* encoding timescale was slower in V1 than in extrastriate (*p*=0.002) but not motor regions (*p*=0.28; Figure 5g). Alpha-band *LLR* encoding timescales monotonically *decreased* across the visual cortical hierarchy (*p*=0.0004, two-tailed weighted permutation test; Figure 5h, compare with c). Taken together, the dynamics (latencies and timescales) of *LLR* encoding in alpha-band lateralization were also consistent with a feedback signal.

Previous monkey^21^ and human^32^ physiology of perceptual choice tasks without change-points have inferred decision-related feedback from the co-variation between intrinsic (stimulus-independent) fluctuations of neural activity and behavioral choice. Here, we computed these fluctuations as the residuals, over and above the activity explained by the components of the normative decision variable depicted in Figure 3g (**Model 6**, *Methods).* Fluctuations in the alpha-/beta-band lateralization of IPS/PCeS, PMd/v, and M1 were robustly predictive of choice toward the end (Figure 6a, right), but not the beginning of the trial (Figure 6b, right). Choice-predictive fluctuation is expected for these downstream regions, because they encode the action plan that ultimately controls the behavioral choice, even when that choice deviates from the choice prescribed by the normative decision variable. Further, this choice-predictive, action-related activity is expected specifically for such late trial intervals because, due to possible changes of mind, decision states earlier in the trial could differ substantially from the choice-determining state toward the end of the trial (Figure 3c).

**Figure 6.**
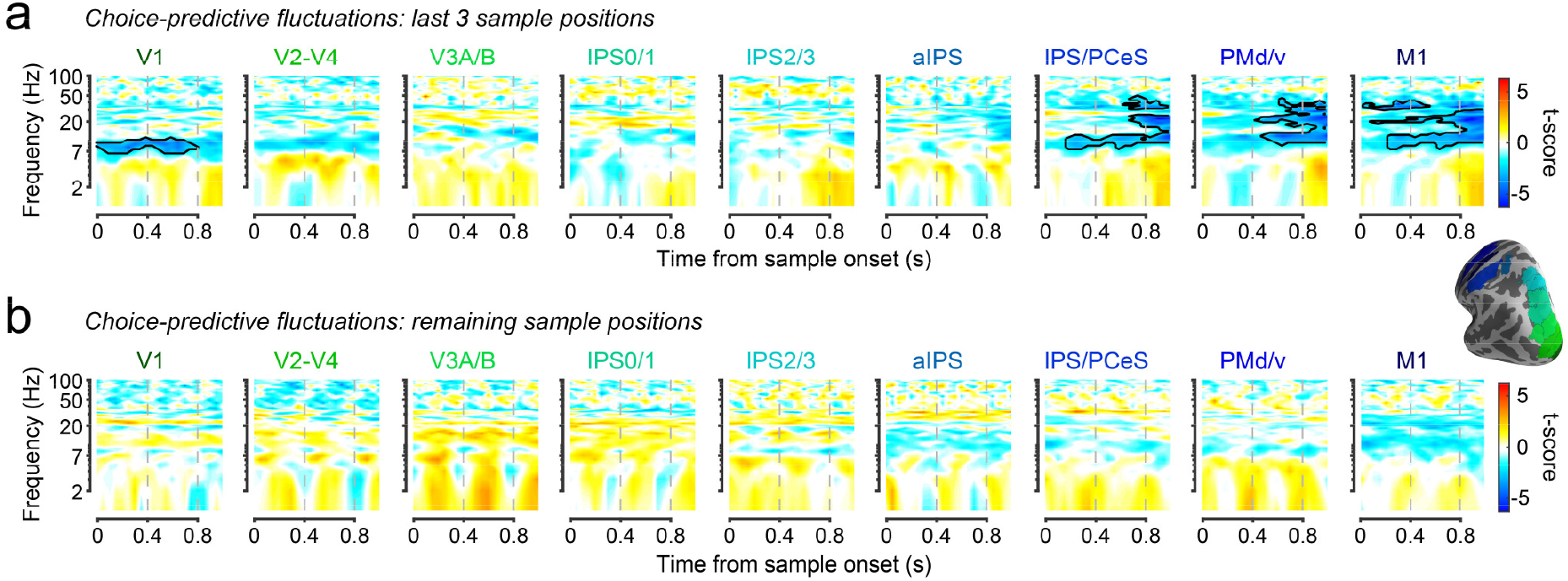
Choice-predictive signals across the visuo-motor pathway. (**a**) Weight of residual fluctuations in the MEG lateralization signal on final choice. Effects shown are averaged over final 3 evidence samples (positions 10-12) in the trial. Colors, group-level t-scores. Contours, p<0.05, two-tailed cluster-based permutation test. (**b**) Same as a, but averaged over all earlier samples positions in the trial; no significant effects in any ROI.

Remarkably, early visual cortex (V1 and weakly in V2-4), but not higher-tier visual field maps, exhibited similarly late, choice-predictive fluctuations, specifically expressed in the lateralization of alpha-band activity (Figure 6a). Again there was no effect earlier in the trial (Figure 6b), ruling out the possibility that this visual cortical alpha-band signal might reflect a biased baseline state before the start of evidence accumulation, due for example to slow attentional drift or choice history effects across trials^21,38^.

Taken together, we found similar encoding of the evolving decision state in the alpha-band in from motor cortex and V1, and more prominent decision signatures when progressing backwards along the visual cortical hierarchy, from extrastriate cortex to V1. Because MEG source reconstruction is limited by signal leakage (*Methods*), a possible concern is that the alpha-band effects observed in visual cortex reflect signal leakage from the downstream motor regions. However, the absence, or even sign reversal (top-middle panel in Figure 4b and Figure 6a), of beta-band effects in visual cortex render this scenario unlikely. Most importantly, the increase of the latency, timescale and (in the case of choice-predictive fluctuations) strength of alpha-band effects with increasing anatomical distance from motor regions shown in Figures 5 and 6 cannot be explained by signal leakage. Thus, the overall pattern of results provides strong support for the notion of feedback of the evolving decision variable to early visual cortex during decision formation.

It has been speculated that decision-related feedback to sensory cortex may stabilize the evolving decision state^19,21^, akin to confirmation bias^39^. Such an active consolidation of the evolving decision through feedback may be beneficial in relatively stationary environments with protracted periods of stability, as was the case in our task (*H*<<0.5). But it would be disadvantageous in highly volatile environments that are likely to change. In another experiment, we assessed if the feedback signatures in early visual cortex *adapt* to changing levels of environmental volatility. We used the same task as in Figure 1, but now manipulated the volatility to be either similarly low as before (*H*=0.1) or high (*H*=0.9) in different blocks (Figure 7a and Methods).

**Figure 7.**
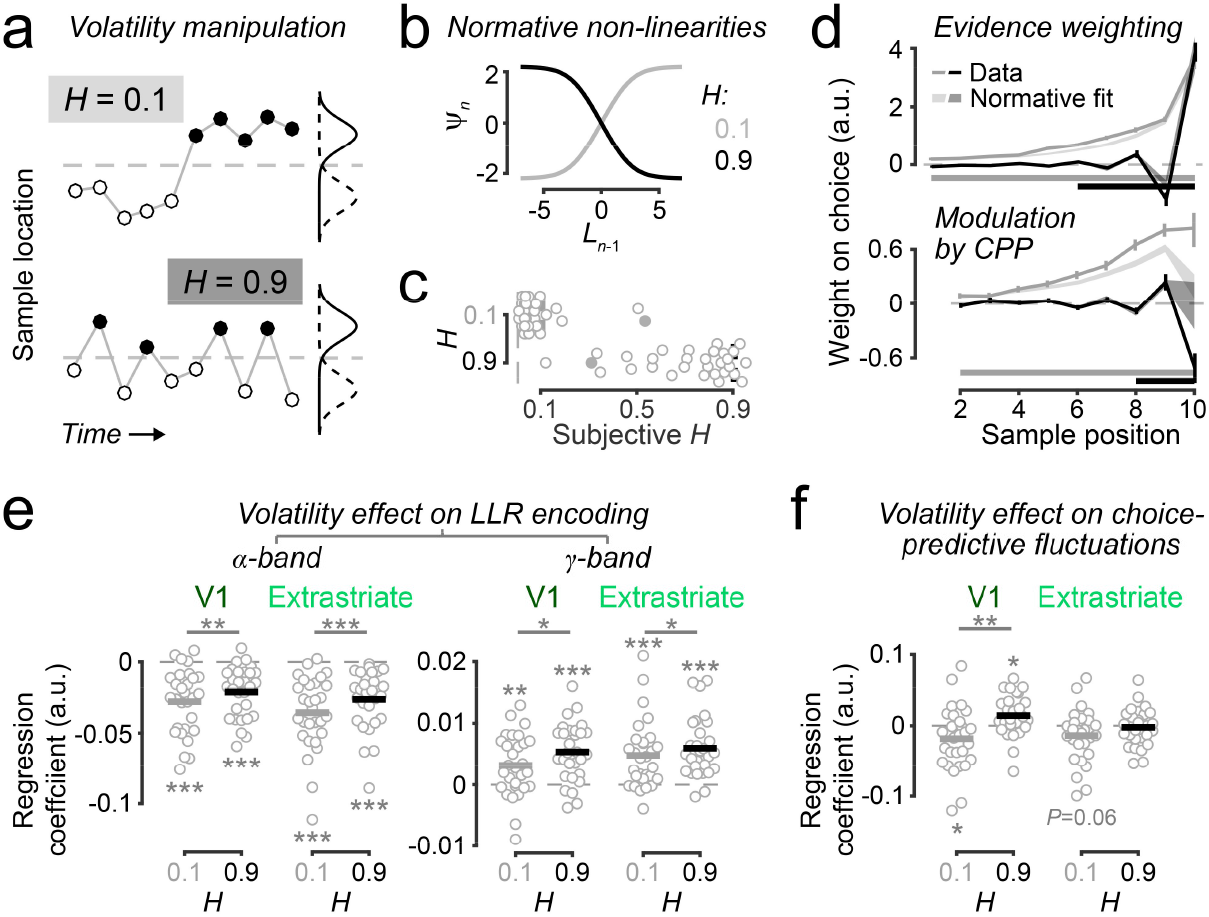
Decision-related feedback in early visual cortex adapts to environmental volatility. (**a**) Schematic of manipulation of environmental volatility. H denotes hazard rate of state changes. For illustration, SNR of generative distributions is higher than that used in experiment. (**b**) Non-linearity in normative model for the two H levels used. (**c**) Subjective H estimated from normative model fits (N=30). Dashed lines, true H. For one participant, H estimates (filled dots) approached 0.5 in both conditions and regression model fits did not converge. Thus, this participant could not be included in the analyses shown in subsequent panels. (**d**) Psychophysical kernels reflecting weight of evidence on choice (left), and its modulation by change-point probability (right). Lines, participants’ data; significance bars, positions where weights differ from zero (p<0.05, two-tailed cluster-based permutation test). Shaded areas, kernels produced by normative model fits. (**e**) Effect of volatility on LLR encoding in the MEG lateralization signal, separately for alpha (8-14 Hz) and gamma (40-65 Hz) frequency bands in V1 and a cluster of extrastriate visual field maps (V2-IPS3). (**f**) Weight of residual fluctuations in a narrow band of the MEG lateralization signal (12-17 Hz, corresponding to the main locus of the choice effect in downstream M1; Supplementary Figure 8c) on final choice. Effects shown are averaged over final 3 evidence sample positions (as in Figure 6a). Panels d,e: *p<0.05, **p<0.01, ***p<0.001; all two-tailed permutation tests.

All but one participant adapted their behavior to these different settings (Figure 7c) and again performed in line with the normative evidence accumulation scheme (Figure 7d, Supplementary Figure 8b; the remaining participant based choices on only the last evidence sample). Please note that the contribution of *-IψI* to normative accumulation was negligible for the generative settings of this experiment (Supplementary Figure 8a). For the low-volatility condition, we again replicated the consistently positive evidence weighting on choice, stronger weighting of late evidence, and up-weighting of evidence associated with high *CPP* (Figure 7d, gray; compare with Figure 2b and Supplementary Figure 3). By contrast, in the high-volatility condition, evidence-weighting time courses now switched sign from sample to sample, again as predicted by the normative evidence accumulation scheme (Figure 7d, black).

For low volatility, we also replicated the pattern of *LLR* encoding (positive for gamma: *p*=0.0028 for V1, *p*<0.0001 for extrastriate regions V2-IPS3; negative for alpha: *p*<0.0001 for V1 and V2-IPS3; Figure 7e) and of choice-predictive fluctuations in the lateralization of early visual cortical activity (*p*=0.036, all two-tailed permutation tests; Figure 7f). Critically, both signatures adapted to the statistics of the environment (Figure 7e,f). The strength of the (negative) alpha-band *LLR* encoding was significantly reduced under high volatility, both in V1 (*p*=0.003, two-tailed permutation test) and extrastriate visual cortex (V2-IPS3 pooled, *p*=0.0002; Figure 7e). The opposite was true for the (positive) *LLR* encoding in the gamma-band, which was enhanced under high volatility (V1: *p*=0.022; V2-IPS3, *p*=0.040). Likewise, the pattern of choice-predictive fluctuations in V1 alpha-band activity also changed from low to high volatility (difference: *p*=0.004) with even a switch from negative to positive sign under high volatility (*p*=0.011 two-tailed permutation test; Figure 7f), possibly indicating an active *destabilization* of the evolving decision state in this environmental setting.

In sum, high environmental volatility seemed to boost the bottom-up sensory evidence encoding (gamma-band) and reduce the feedback signaling in early visual cortical alpha-band activity, in a fashion that was flexibly adapted to the stability of the environment.

### Phasic arousal modulates evidence accumulation and the state of visual cortex

Cortical circuit dynamics are not only shaped by intra-cortical feedback signals but also by input from brainstem arousal systems^33,40–42^. Transient (phasic) arousal signals, specifically from the noradrenaline system, might dynamically adjust the impact of new evidence on belief updating, particularly when the evidence is surprising^43–45^. In a final set of analyses of data from the main experiment, we used rapid, sample-evoked dilations of the pupil to assess the involvement of brainstem arousal^46–48^. We quantified rapid pupil responses as the first derivative (i.e. rate of change) of the raw response in order to increase temporal precision (Figure 8a) and specificity for noradrenaline release^49^.

**Figure 8:**
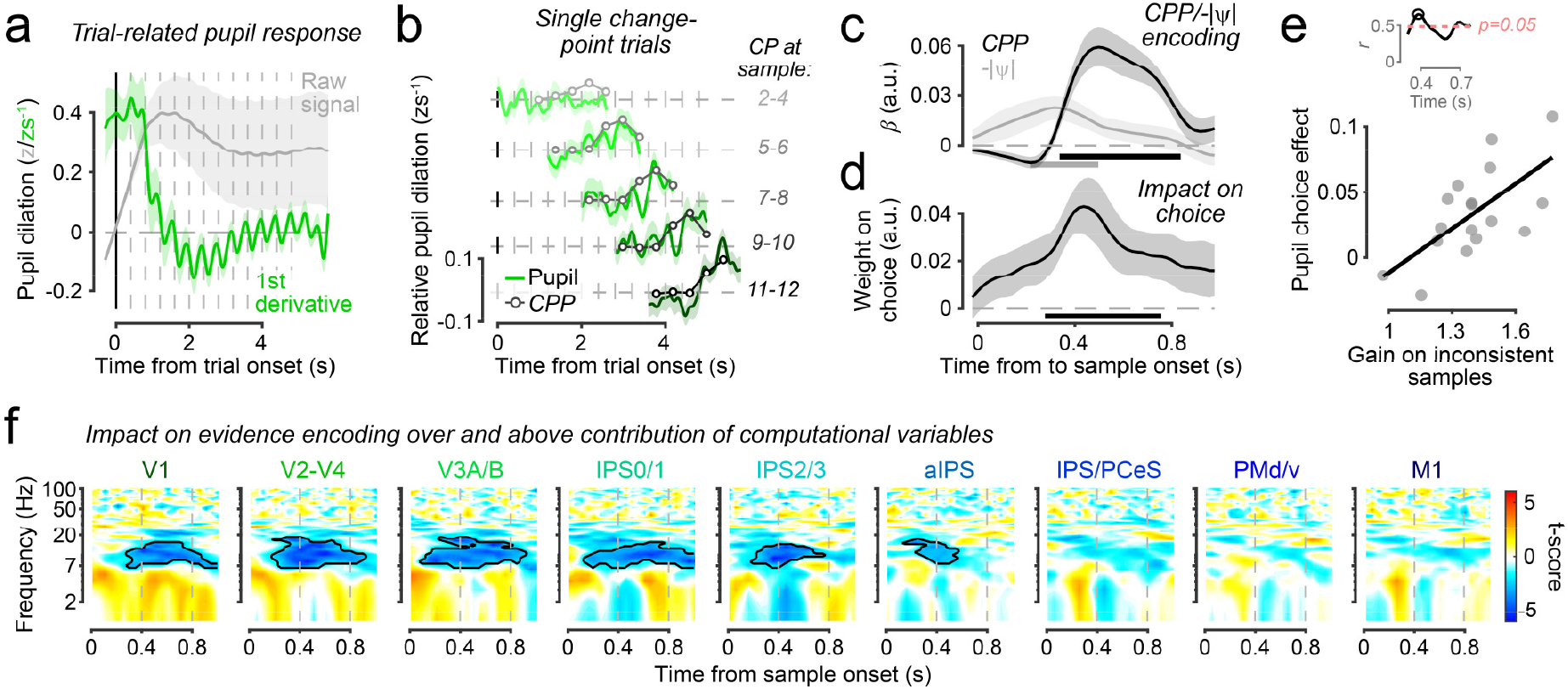
Arousal responses to perceived state changes up-weight the associated evidence in brain and behavior. (**a**) Average trial-related pupil response (grey) and its first derivative (green). Dashed grey vertical lines, sample onsets. (**b**) Pupil derivative for subsets of trials where only single change-points (CP) occurred at different latencies, relative to pupil derivative on trials without any state change. Greyscale traces, relative change-point probability for same trials from normative model fits (scaled and time-shifted by same factors across trial groups). Traces truncated from 3 samples before to 1 sample after average time of state change. (**c**) Encoding of changepoint probability and uncertainty in pupil responses to individual samples. (**d**) Multiplicative effect of intrinsic, singletrial pupil derivative fluctuations on weighting of evidence samples in choice. (**e**) Across-participant correlation of pupil choice effect (panel d) with fitted model parameter reflecting additional weight ascribed to new evidence that was inconsistent with belief. Inset, time-resolved Pearson correlation coefficients spanning significant temporal cluster from panel d; highlighted time-point used to generate scatterplot. (**f**) Multiplicative effect of pupil response to individual samples on encoding of new evidence in the lateralized, regionally-specific MEG signal. Colors reflect group-level t-scores. Contours, p<0.05 (two-tailed cluster-based permutation test). Shaded areas in panels a-d indicate s.e.m. Significance bars in c and d, p<0.05 (two-tailed cluster-based permutation test).

Samples strongly indicative of a change in environmental state evoked strong pupil responses (Figure 8b-c). We quantified this effect by fitting a linear model consisting of model-derived variables including the *CPP* and *-IψI* associated with each sample to the corresponding pupil response (**Model 7**, *Methods).* We found a robust positive contribution of *CPP* (*p*<0.0001, twotailed cluster-based permutation test) in addition to a weaker positive contribution of -I*ψ*I (*p*=0.023). Thus, in our task, phasic arousal was recruited by the two computational quantities – *CPP* and *-IψI* – that modulate evidence weighting in the normative model.

The sample-to-sample fluctuations in the evoked arousal response predicted an ‘up-weighting’ of the impact of the associated evidence sample on choice (Figure 8d-f). Variations in pupil responses beyond those explained by the above computational variables (**Model 8**, *Methods)* exhibited a positive, multiplicative interaction with *LLR* in its impact on choice (Figure 8d; *p*=0.003, two-tailed cluster-based permutation test). The magnitude of this effect also correlated with a participant-specific gain parameter, estimated in our normative model fits, which quantified the weighting of belief-inconsistent evidence beyond the upweighting entailed in the normative model (Figure 8e; peak *r*=0.66, uncorrected *p*=0.004, two-tailed).

The pupil-predicted up-weighting of the behavioral impact of evidence (Figure 8d) was accompanied by a corresponding modulation of *LLR* encoding across the visual cortical hierarchy (Figure 8f). Here, we expanded our decomposition of cortical dynamics (Figure 4) by a term that reflected the modulation of evidence encoding by the associated pupil response (**Model 9**, *Methods).* This showed a transient modulatory effect of pupil responses on selective evidence encoding in the power lateralization of visual cortical areas (Figure 8f). The selective pupil-linked effect was superimposed onto a more sustained, global (across hemispheres) effect on low-frequency power across the visuo-motor pathway (Supplementary Figure 9). The selective effect was specifically expressed in alpha-band power lateralization (Figure 8f) and not evident in downstream, action-related regions (all *p*>0.3; Figure 8f). Thus, phasic arousal modulated evidence encoding in the visual cortical system in the specific frequency-band that our previous analyses of the same neural data had linked to decision feedback.

## Discussion

Computational analyses have developed normative models of evidence accumulation for solving decision-making tasks like the one used here^16^. Other work has dissected the synaptic interactions implementing cognitive computations, leading to an influential circuit model for evidence accumulation^18–20^. We aimed to develop an integrated understanding of evidence accumulation across these different levels of analysis^50^. We reasoned that the possibility of hidden change, a fundamental feature of natural environments^51^, may be critical in this pursuit. Some studies have started to explore how perceptual decisions are made in changing environments^16,17,23^, but focused on algorithmic-level modeling of behavior. Other work has characterized the dependence of neural activity in sensory cortex on evidence volatility (roughly analogous to the variance of evidence samples in our task) but under a fixed environmental state^52^.

Here, we have identified the cortical representation of an approximately normative decision variable for changing environments. We found that this decision variable is encoded in the buildup activity of cortical regions involved in action planning and that its adaptive, non-linear computation naturally emerges from recurrent synaptic interactions in cortical microcircuits. In addition, we show that the state of visual cortex, expressed in activity fluctuations in the alpha frequency band, is shaped by both an adaptive, stabilizing feedback of the evolving decision variable and pupil-linked, phasic arousal. We conceptualize ‘cortical state’ as modulations of cortical activity distinct from feedforward sensory responses. Our data are in line with the notion that such state modulations emerge from the interplay of selective cortical feedback and neuromodulatory signals^53^. This interplay culminates in selective changes in the lateralization of alpha power with respect to the incoming evidence and evolving decision. The first part of our results (behavior and the activity of action-planning regions) forge a tight link between existing circuit models and normative evidence accumulation, while the second part (state dynamics in visual cortex) call for an extension of circuit models for evidence accumulation by (i) multiple processing stages with feedback and (ii) neuromodulatory input. We propose that these features enable (biological or artificial) circuits to effectively adapt evidence accumulation and the resulting behavior to a range of environmental statistics.

Our account starts from the realization that adaptive evidence accumulation in changing environments typically entails the non-linear modulation of evidence weighting by changepoint probability and uncertainty. The impact of these modulations varies across environments. They are negligible in contexts that preclude the formation of strong belief states: high (external or internal) noise and/or a rate of change that makes the environment generally unpredictable (*H*≈0.5). Correspondingly, a leaky (linear) accumulator fits the behavior of rats with strong internal noise well in an auditory task similar to ours^23^. Even so, these modulations make a significant contribution to normative accumulation in a wide range of environmental contexts (Supplementary Figure 1). We reiterate that change-point probability and uncertainty are not an explicit component of the normative model used here. Rather, the variables served as a vehicle for identifying diagnostic behavioral and neural signatures of the adaptive evidence accumulation scheme. This contrasts with other models of decision-making in changing environments, which make explicit use of measures of surprise and uncertainty^45,54,55^ (see Supplementary Figure 10 for the relationship between change-point probability and commonly-used measures of surprise). In line with other work^44,45^, we found that both change-point probability and uncertainty were robustly encoded in participants’ pupil-linked arousal responses to evidence samples (Figure 8c). In fact, pupil responses encoded change-point probability more strongly than alternative measures of surprise (Supplementary Figure 10e). These observations further validate the use of change-point probability and uncertainty in our dissection of the adaptive accumulation process.

Previous work on statistical learning has shown that humans adaptively tune their learning rate as a function of change-point probability (or other forms of surprise) and uncertainty^54,56,57^. Indeed, some of these effects have been linked to pupil dynamics^45^. Learning tasks commonly operate on timescales at least an order of magnitude slower than our perceptual evidence accumulation task. Yet, there is a direct analogy between the dynamic adjustment of learning rate by surprise/uncertainty and pupil-linked arousal and the dynamic modulations of evidence weighting we identified here. An interesting question for future research is whether this analogy originates from shared circuit mechanisms – and, specifically, to which extent the circuit operations identified here can be stretched from perceptual evidence accumulation over seconds to learning over minutes or longer. At the neural level, the previous work on dynamic adjustments of learning has focused on identifying correlates of the variables that control the regulation (i.e., surprise or uncertainty) across the brain^54,56,57^. By contrast, our current study illuminated how such computational variables modulate the selective cortical signals that encode the output of the adaptive computation, the evolving belief state itself.

The computation of the decision variable entails intra-cortical, recurrent network interactions that produce timing jitter^20,58^. Consequently, neural signatures of the decision variable may not be precisely phase-locked to the onset of evidence samples, but rather manifest in slower variations of the amplitude of local field potential (MEG signal) fluctuations^58^. Further, computational variables are commonly encoded in specific frequency bands of the signal fluctuations, sometimes with opposite sign between bands^58,59^. We therefore reasoned that our frequency-resolved encoding analysis should be well-suited for detecting correlates of the decision variable (see also *Methods, Spectral analysis).* Our results corroborate this reasoning, and highlight the potential of the approach for tracking cognitive computation across time and cortical space.

Our results also support the notion that attractor dynamics in frontal and parietal cortical microcircuits implement the formation of categorical decision states and their maintenance in working memory^18,20,60,61^. A circuit model with such attractor dynamics (Figures 2, 3, Supplementary Figure 6) accurately reproduced the detailed characteristics of both behavior and build-up activity in parietal and frontal cortical regions involved in action planning. One signature of such cortical attractor states during decision-making is the sigmoidal relationships of cortical activity and model decision variables found here (Figure 3e) and elsewhere^9,37^. Critically, we identified an adaptive function of these attractor dynamics: balancing stable accumulation of the evidence and flexible response to change, which results from a combination of properties of the attractor dynamics (Supplementary Figure 6). Even so, these circuit dynamics will only be beneficial in a limited range of environments, and changes in environmental statistics likely require a tuning (e.g. through neuromodulatory input) or reconfiguration (e.g. through adaptive recruitment of long-range cortical feedback connections) of the local or large-scale circuit properties.

Feedback signaling to sensory cortex from downstream cortical regions during perceptual evidence accumulation could convey predictions that are compared with incoming sensory data in an iterative fashion^62^. The feedback may also mediate the impact of expectations inherited from previous trials, through mechanisms akin to attention^21,38^. Finally, when instigated early during decision formation and originating from downstream accumulator circuits, feedback may consolidate the evolving network state^19,32,39^. Our current results show that decision feedback emerges during (not before) evidence accumulation. Critically, they also establish the dependence of this feedback on environmental volatility in line with an actively recruited, adaptive mechanism. These findings support the notion of an active stabilization of evolving belief states through feedback. Relatedly, our results also provide a new perspective on the idea that feedforward vs. feedback signaling across the visual cortical hierarchy is multiplexed in gamma-vs. alpha-frequency channels^34,35^: We establish the sensitivity of these channels to higher-level environmental statistics (volatility). The particularly pronounced and adaptive signatures of decision feedback in the alpha-band activity of V1, which were stronger than in any other visual cortical area, point to a special role for early visual cortex in perceptual evidence accumulation.

An influential account holds that surprise-related phasic responses of the brainstem noradrenaline system cause a shift toward more ‘bottom-up’ relative to ‘top-down’ signaling across the cortical hierarchy, and thus a greater impact of new evidence on the evolving belief^43,45^. Our observation that pupil responses to evidence samples indicative of environmental state change predicted an up-weighting of the impact of that sample on choice is broadly consistent with this idea. However, the associated modulations of cortical activity were inconsistent with a simple strengthening of the bottom-up processing of the evidence. Such an effect should have been evident in visual cortical gamma-band responses and produced corresponding changes in action-related regions (i.e., stronger evidence encoding in motor alpha-/beta-activity). Instead, pupil-linked arousal modulated evidence encoding only in visual cortex, not in downstream regions, and only in the alpha-band, not the gamma-band, consistent with an enhancement of feedback signals to visual cortex. The effect of the arousal signal also occurs relatively long after the arrival of the new evidence sample. It is tempting to speculate that, in the current settings, feedback enhancement through phasic arousal effectively stabilizes the updated belief state induced by surprising evidence.

To conclude, taking explicit account of the volatility of natural environments, our approach uncovers a direct link between the non-linear dynamics of cortical circuits and adaptive decision computation. Our results also call for the extension of current circuit models for decision-making by cortical feedback and modulation through ascending arousal systems.

## Methods

This paper reports data from three experiments, all using different variants of the same behavioral task (Figure 1): two MEG experiments and one behavioral control experiment. Experiment 1 was the main experiment of this study (Figures 1–6, 8; Supplementary Figures 1–2, 4–7, 9–10) and required participants to perform the task at a fixed level of environmental volatility during MEG recordings. Experiment 2 was a behavioral control experiment, in which participants performed the task under two different sample onset asynchronies (Supplementary Figure 3). Experiment 3 was a follow-up MEG experiment to Experiment 1, which required participants to perform the task under two levels of volatility (Figure 7; Supplementary Figure 8).

### Participants

The study was approved by the ethics committee of the Hamburg Medical Association. All participants provided written informed consent before the start of testing. All had normal or corrected-to-normal vision and no history of psychiatric or neurological diagnosis.

*Experiment 1.* Seventeen human participants (mean ± s.d. age of 28.22 ± 3.89 years, range 23-36; 11 females), including the first author of this manuscript, took part in the experiment. Each participant completed one training session (120 min), three (16 participants) or four (1 participant, the first author) sessions of the main experiment (about 150 min each), and one session to obtain a structural MRI (30 min). Participants received remuneration for their participation in the form of an hourly rate, a completion bonus and an additional performance-dependent bonus.

*Experiment 2.* Four participants (ages of 25, 26, 27, and 28 years; 2 males) took part in the experiment. Each participant completed one training session (120 min), followed by three (1 participant) or four (3 participants) sessions (about 120 min each) that provided behavioral data used for the analysis. Participants received remuneration according to the same scheme used for Experiment 1.

*Experiment 3.* Thirty participants (age=26.07 ± 3.49 years, range 20-35; 18 females), including the first author of this manuscript, took part. Each participant completed one training session (120 min), three sessions of the main experiment (about 480 min each), and one session to obtain a structural MRI (30 min). Participants received remuneration according to the same scheme used for Experiments 1 and 2. This experiment also included a within-subject pharmacological intervention (40 mg atomoxetine, 5 mg donepizil, placebo; one per session, randomized and double-blind; results reported here from pooling data over drug conditions) and was subject to additional participant exclusion criteria: smokers, individuals with an average consumption of >15 units of alcohol per week, illegal drug users, pregnant females, individuals taking medication for chronic disease (e.g. asthma, hyperthyroidism), and individuals with narrow-angle glaucoma, pheochromocytoma, cardiovascular disease or known hypersensitivity to atomoxetine or donepezil were all excluded from participation. A comparison of the drug effects is beyond the scope of this study and will be reported in a separate paper.

In Experiment 3, choice regression model fits for one participant did not converge. Fits of the normative model showed that this participant had subjective hazard rates approaching 0.5 in both volatility conditions (see Figure 7b, filled dots), consistent with use of a non-adaptive heuristic of basing decisions only on the sign of the last evidence sample in each trial. This participant thus had to be excluded from all analyses of the behavioral and MEG data using regression models.

### Main behavioral task

#### Task design

The main task was a two-alternative forced choice discrimination task, in which the generative task state *S={left, right}* change unpredictably over time (Figure 1a). On each trial of the task in Experiment 1, participants viewed a sequence of evidence samples consisting of small checkerboard patches (see below: *Stimuli*). Samples were presented for 300 ms each, with sample-onset asynchrony (SOA) of 400 ms. The 100 ms blank between samples interval ensured that each patch was differentiable from a temporally adjacent patch even in rare cases when they overlapped in space. Samples were centered on spatial locations (specifically, polar angles) *x*_1_,...,*x_n_* drawn from one of two probability distributions *p*(*x*|*S*). These distributions *p*(*x*|*S*) were truncated Gaussians, with matched variance (*σ_left_*=*σ_right_*=29°), means symmetric with respect to the vertical midline (*μ_left_*=-17°, *μ_right_*=+17°), and truncated at −90° (+90°). In instances where a drawn *x_i_* was <−90° (>+90°), it was replaced with −90° (+90°). The generative state at the start of each trial (i.e., the distribution generating the first sample), was chosen at random. After the presentation of each sample, *S* could change with a fixed probability, or hazard rate: *H=p*(*S_n_=right|S_n-1_*=left)=*p*(*S_n_=left|S_n-1_*=*right*)=0.08. Participants were asked to maintain fixation at a centrally presented mark (see below: *Stimuli*) throughout the sample sequence, monitor all samples, and report *S* at the *end* of the sequence. That is, participants needed to infer which of the two probability distributions generated the position of the final sample.

The majority (75%) of sequences in each block of trials contained 12 samples. The lengths of the remaining sequences (25%, randomly distributed throughout each block) were uniformly distributed between 2 and 11 samples. Thus, the sequence durations ranged between 0.8 s (2 samples) and 4.8 s (12 samples). The shorter sequences were introduced in order to discourage participants from ignoring the early samples in the sequence and encourage them to accumulate evidence over time. For 12-sample sequences the hazard rate of *H*=0.08 yielded 39.9% of trials with no state change, 38.0% with one state change, and 22.1% with >1 state change.

Before the onset of each trial, there was a preparatory period of variable duration (uniform between 0.5 and 2.0 s) during which participants were instructed to maintain fixation and a stationary checkerboard patch was presented at a central location (0° polar angle). Trial onset was signaled when this checkerboard began to flicker, and 400 ms later the first evidence sample was presented (see *Stimuli).* Immediately after the sample sequence, the stationary patch was again presented at 0° and remained there until the start of the following trial. After a variable interval following sequence completion (uniform between 1.0 and 1.5 s) a ‘Go’ cue instructed participants to report their choice via button press with the left or right thumb (indicating state *left* or *right,* respectively). Auditory feedback was provided 0.1 s post-response informing the participant about the accuracy of their choice relative to the true generative state at the end of the sequence (see *Stimuli).* A rest period of 2 s immediately followed feedback, during which participants were instructed to blink if necessary, and this was followed by the preparatory period for the next trial.

Experiments 2 and 3 used the same task, with the following exceptions. In Experiment 2, samples could be presented either for 100 ms with a 200 ms SOA, or for 500 ms with a 600 ms SOA. Presentation duration and SOA remained fix within a task block (see below), but alternated between task blocks. In Study 3, *H* was set to either 0.1 or 0.9. The *H* level at the start of each session was chosen pseudo-randomly, under the constraint that the starting *H* be chosen equally often across the group for each of the three sessions, and every participant would start with each *H* level at least once. It then remained fixed for 8 task blocks (see below), participants were given a 45-to 60-minute break, and *H* then switched to the other level for another 8 task blocks. The sample generative distributions in Experiment 3 were parameterized as follows: *μ_left_*=-28°, *μ_right_*=+28°, *σ_left_*=*σ_right_*=28°. Also in Experiment 3, the majority (65%) of sample sequences in each task block contained 10 samples; the lengths of the remaining sequences (35%, randomly distributed throughout each block) were uniformly distributed between 2 and 9 samples; the duration of the pre-trial preparatory period was 0.5-1.5 s; the duration of the variable interval following sequence completion and preceding ‘Go’ cue was 0.7-1.2 s; and the duration of the post-trial rest period was 1.2 s.

#### Stimuli

All visual stimuli described below were presented against a grey background. Two placeholder stimuli were on screen throughout each block of trials: a light-grey vertical line extending downward from fixation to 7.4 degrees of visual angle (d.v.a.) eccentricity; and a colored half-ring in the lower visual hemifield (polar angle: from −90 to +90°; eccentricity: 8.8 d.v.a.), which depicted the log-likelihood ratio associated with each possible sample location (see below). The colors comprising this half-ring, along with those of the fixation point, were selected from the Teufel colors^63^.

Each evidence sample consisted of a black and white, flickering checkerboard patch (temporal frequency: 10 Hz; spatial frequency: 2 d.v.a.) within a circular aperture (diameter = 0.8 d.v.a.). The patches varied in position from sample to sample, whereby their eccentricity was held constant (8.1 d.v.a.) and polar angle varied across the lower visual hemifield (i.e., from −90 to +90°.). The colored half-ring described above was presented at a larger eccentricity than the checkerboard patches to avoid overlap.

The fixation mark, which was presented in the center of the screen, was a black disc of 0.18 d.v.a. diameter superimposed onto a second disk of 0.36 d.v.a. diameter and with varying color. The fixation mark was present throughout each task block, with the color of the second disk informing participants about task-relevant events. During the preparatory period, sample sequence presentation, and subsequent delay, the color of the second disk was light red; the ‘Go’ cue took the form of the second circle becoming light green; and, the inter-trial rest period was indicated by the second circle becoming light blue.

Auditory feedback consisted of a 0.25 s ascending 350→950Hz tone if the choice was correct and a descending 950→350Hz tone if incorrect.

### Task training protocol

Each participant completed a task training protocol in a separate session that took place in a behavioral psychophysics room one week or less prior to the first main experimental session. The protocol started with a visual illustration of each generative distribution, explained through analogy with a deck of cards. The participant then completed 12 trials of a static version of the task (*H*=0), after which they received feedback on their mean choice accuracy and were given the option to complete another 12 trials. The experimenter enforced repetition in cases where accuracy was less than 90%.

Next the participant was informed that, during the real task, the deck that was being used to draw the dot locations could change at any time within a sequence of dots. In Experiments 1 and 2, this was explained through analogy with a biased coin flip between each dot presentation, where a ‘heads’ would be the outcome on “about 11 out of 12 flips” and would not change the deck, while a ‘tails’ would be the outcome on “about 1 out of 12 flips” and would change the deck. The participant then performed 20 trials of the full task (*H*=0.08; at the longer of the two possible SOAs in Experiment 2) in which, whenever a change in generative state occurred, the text string ‘CHANGE!’ (22-point white Helvetica font) appeared 1.45 d.v.a. above fixation for the duration of the following dot. Participants were informed that they could use this change-point signaling to help them make decisions during the training exercise but that it would not appear during the real task. They again received feedback on their mean choice accuracy and were given the option to complete another 20 trials, with the experimenter enforcing repetition in cases where accuracy was less than 80%. Lastly, participants completed a median of 5 (range = 2-6) blocks of the full task, without change-point signaling, at fixed SOA in Experiment 1 or alternating SOAs (200 or 600 ms) in Experiment 2. Each block consisted of 76 trials, after which participants received feedback on their mean choice accuracy in that block and took a short, self-timed break.

In Experiment 3, the procedure was identical with the following exceptions. The coin flip analogy was also used during participant instruction but no specific numbers were mentioned about the frequency of deck changes in order to avoid biasing subjective *H* estimates across conditions. Participants received the following information: “For some task blocks, switches will not be too frequent and on some trials there will be no switches at all. For other blocks, switches will be extremely frequent and on most trials, multiple switches will occur in a row.” They then performed 20 trials of the full task with *H*=0.1, including the same overt changepoint signaling as above and the option to repeat if desired by either participant or experimenter. Next they performed 20 trials with *H*=0.9, again with the option to repeat. Lastly, they completed two blocks of the full task with *H*=0.1 followed by two blocks with *H*=0.9 without change-point signaling. Each block consisted of 86 trials and was followed by feedback on mean accuracy and a short break.

### Localizer tasks

Within each session of the main task used in the two MEG experiments (Experiments 1 and 3), participants also completed a single block of each of two ‘localizer’ tasks for (i) decoding the position (polar angle) of the checkerboard patterns from visual cortical responses and (ii) measuring motor preparatory activity for the hand movement used to report the choice, without participants performing the decision-making task described above. Data from the polar angle localizer task were not used for the current study.

We used a delayed hand movement task to measure motor preparation in the absence of evidence accumulation and decision-making. The task was analogous to the delayed saccade task used to identify neurons encoding saccade plans in occulomotor structures of the monkey brain in previous studies of visual decision-making (e.g. ref ^64^). Participants fixated the central fixation mark while a sequence of lexical cues and subsequent ‘Go’ cues for responding were presented. A trial began with the presentation of one of two lexical cues (‘LEFT’ or ‘RIGHT’; 15-point white Trebuchet font; 0.3s duration) that appeared 1.25° above the central fixation mark (of same construction as above; color of second disk light red). The participants’ task was to prepare the associated motor response (left thumb button press for ‘LEFT’; right thumb for ‘RIGHT’) during a following 1 s blank delay period within which only the fixation mark was presented, and to execute that response as quickly as possible when the delay was over (marked by the second disk turning light green). The second fixation disk became light blue for 2 s post-response, indicating a rest period during which participants were instructed to blink if necessary. The fixation mark color returned to light red after the rest period and, after a variable interval (uniform distribution with bounds of 0.75-1.5 s), the next trial began. A block of the motor preparation task was comprised of 60 trials (30 left and 30 right responses, randomly distributed), and was administered after the sixth block of the decision-making task in each session.

### Procedure and Apparatus

For Experiments 1 and 2, the experimental sessions for each participant took place on consecutive days. They comprised between 7 and 9 blocks depending on time constraints, and included 2-minute breaks between blocks and one longer break of approximately 10 minutes in the middle of the session. Like during training for these Experiments, each block was comprised of 76 trials and followed by feedback about choice accuracy, now including the mean for that block and a running mean for that session. In Experiment 1, participants completed a median of 26 blocks (range=22–33), corresponding to a median of 1976 trials (range=1628–2508). In Experiment 2, the four participants completed 33, 37, 45 and 47 blocks of mostly 76 trials each (a small subset curtailed for time purposes), corresponding to 2100, 2581, 2970 and 3174 trials. For Experiment 3, experimental sessions for each participant usually comprised 16 blocks, including 2-minute breaks between blocks, approximately 5-to 10-minute breaks between blocks 4/5 and 12/13, and a longer 45-to 60-minute. Each block was comprised of 86 trials (again, a small subset curtailed for time purposes) and followed by feedback about choice accuracy in the same manner as Studies 1 and 2. Study 3 participants completed a median of48 blocks in total (range=42–51), corresponding to a median of 4087 trials (range=3612-4386).

All stimuli were generated using Psychtoolbox 3 for Matlab^65,66^. Visual stimuli were back-projected on a transparent screen using a Sanyo PCL-XP51 projector with a resolution of 1920×1080 at 60 Hz (presented on a VIEWPixx monitor during training and all of Study 2 with the same resolution and refresh rate). Subjects were seated 61 cm from the screen in the MEG room, or with their head in a chinrest 60 cm from the monitor (training and all of Study 2) in an otherwise unlit room.

### Normative model and derivation of change-point probability and uncertainty

The normative model for solving the inference problem in the above decision-making task prescribes that incoming evidence about the generative task state is accumulated over time in a manner that balances stable belief formation with fast change detection^16^:

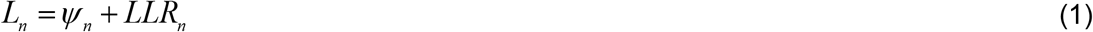

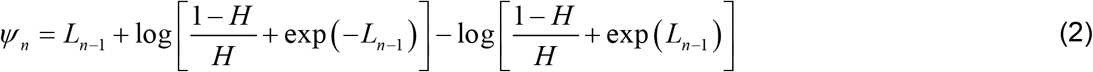

Here, *L_n_* was the belief of the observer after having observed the *n^th^* sample of evidence, and expressed in log-posterior odds of the alternative task states; *LLR_n_* was the relative evidence for each alternative carried by the *n^th^* sample, expressed as a log-likelihood ratio (*LLR_n_* = *log*(*p*(*x_n_*|*right*)|*p*(*x_n_*|*left*))); and *ψ_n_* was the prior expectation of the observer before encountering the *n^th^* sample. The key feature that sets this model apart from traditional evidence accumulation models^1^ is the transformation of *L_n-1_* into *ψ_n_* (Equation 2) and how this takes into account the hazard rate (i.e., probability of state change), *H*. In the special case that *H*=0 (no state changes), the two rightmost terms in Equation 2 cancel and Equation 1 reduces to perfect evidence accumulation, as in drift diffusion^1^. When *H*=0.5 (no state stability), *ψ_n_* cancels and so no evidence is accumulated (posterior belief depends only on current evidence). For intermediate values of *H*, the model strikes a balance between these extremes, and thus between stability and sensitivity to change (i.e. flexibility; Supplementary Figure 1).

We used this previously established model to derive two computational quantities: *CPP* and – |*ψ*|. This exercise was motivated by similar treatments for a different form of change-point task (continuous belief updating^45^). The goal was to recast the non-linearity in eq. 2 in a way that would directly illuminate the underlying neural computations and hence, decision-related MEG and pupil signals.

#### Change-point probability

We derived a formal expression for the probability that a change in generative task state has just occurred given the expected *H*, the evidence for each state carried by a newly encountered sample *x_n_*, and the observer’s belief before encountering that sample *L_n-1_*. We refer to this quantity as *CPP*. In what follows we denote all encountered samples up to and including the new sample as *X*_j∈N_, refer to *right* and *left* states as *S*_1_ and *S*_2_ for convenience, and note that the log-posterior odds *L* can be re-expressed as the probability of each generative state: *p*(*S*_1_)=1-*p*(*S*_2_)=1/(*e^L^*+1).

The task structure enabled two state transitions that define a change-point: *S*_1_→*S*_2_ and *S*_2_→*S*_1_; and two transitions that are not a change-point: *S*_1_→*S*_1_ and *S*_2_→*S*_2_. Thus, we could write the probability of a change-point transition as follows:

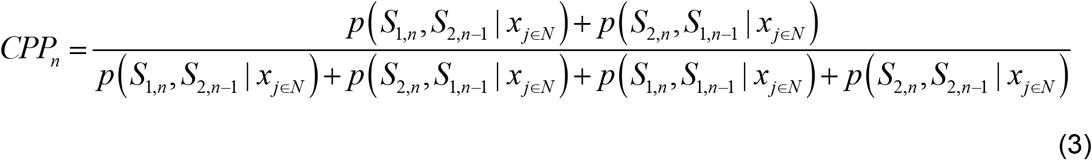

where the denominator accounted for all possible state transitions between consecutive samples *n*-1 and *n*. Using the definition of conditional probabilities and separating the set of all observed samples into the new sample *x_n_* and all previous samples *x*_1...*n*-1_, we decomposed each of the four possible state transition probabilities in the following way:

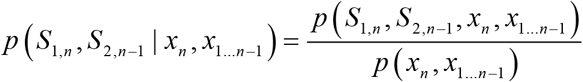

This expression could be expanded via the product rule:

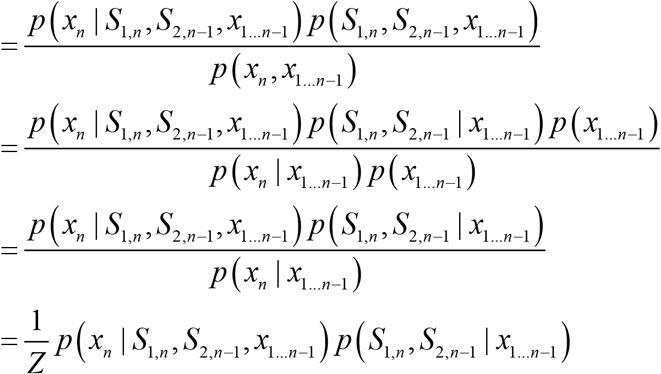

where *Z* = *p*(*X_n_*|*X*_1...*n*-1_). Because previous observations and beliefs are irrelevant for determining the probability of a new sample or state transition when the current generative state is known, we could simplify the above expression and expand via the product rule:

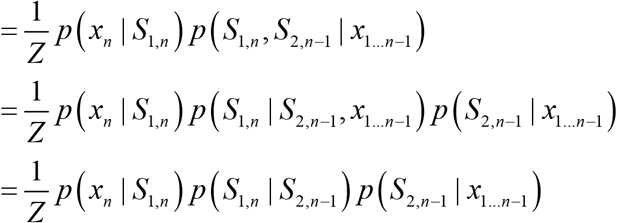

As established previously, *p*(*S*_1,*n*_|*S*_2,*n*-1_)=*p*(*S*_2,*n*_|*S*_1,*n*-1_)=*H*’ thus reducing the above to:

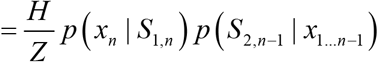

Finally, in our case the generative stimulus distribution *p*(*x|S*_1_) is Gaussian such that:

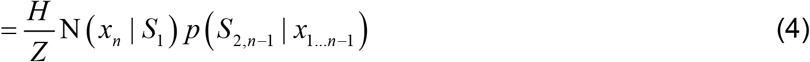

where N(*X*|*S*) denoted the probability of sample *X* given a normal distribution with mean *μ_s_* and s.d. *σ_S_*. As described above, the final term in eq. 4, which we abbreviated to *p*(*S*_2,*n*-1_) below, could be computed directly from *L*_*n*-1_.

The remaining transition probabilities in eq. 3 could be derived analogously to eq. 4. Replacing each term in Equation 3 with the expressions derived by doing so, the *Z* terms cancelled to yield the following expression for *p*(*CP*) that could be readily computed:

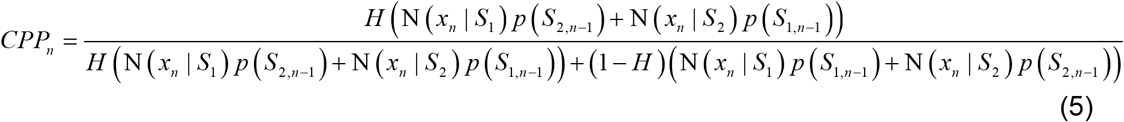

This quantity has intuitive characteristics (Figure 1d). First, the numerator computes a weighted sum of the likelihood of the new sample under both generative states assuming that a change has occurred, with each weight determined by the strength of the observer’s existing belief in the *opposing* state. This means that a new sample of evidence that is inconsistent with the observer’s belief (i.e. sign(*LLR*_n_)≠sign(*L*_*n*-1_)) will yield a larger *CPP* than a sample that is consistent. Second, if the new sample carries no information about the current generative state (i.e. *LLR*_*n*=0_), eq. 5 evaluates to *H*. In other words, when a new sample is ambiguous, the observer must rely more on their base expected rate of state change as an estimate for *CPP.* Similarly, if the observer is completely agnostic as to the task state (i.e. *L*_*n*-1_=0), eq. 5 again evaluates to *H*. That is, a belief about a change-point having occurred over and above the base expected rate of change can only form if the observer has some level of belief in the task state before encountering the new sample.

#### Uncertainty

We also defined belief uncertainty prior to observing a new evidence sample *X_n_* as –|*ψ_n_*|. Thus, the closer the observer’s prior belief to the category boundary of zero, the higher their uncertainty (Figure 1e). This measure thus identifies instances when the observer is at the steepest part of the non-linearity of the normative model, where belief updating is generally strong.

#### Influence of change-point probability and uncertainty on evidence accumulation

We used simulations to understand the influence of *CPP* and –*|ψ|* on evidence accumulation in the normative model. We evaluated this influence as a function *H* and signal-to-noise ratio (SNR) of the generative distributions (defined as the difference between distribution means divided by their matched s.d.). For each point on a 5×5 grid (*H* ={0.01, 0.03, *0.08*, 0.20, 0.40}, SNR={0.4, 0.7, *1.2*, 2.0, 5.0}; italicized values match the generative statistics of our task in Experiments 1 and 2), we simulated a sequence of 10,000,000 observations, passed these through the normative accumulation rule described by eqs. 1 and 2, and calculated the persample computational variables described above. We then assessed the influence of different variables on belief updating by fitting the following linear model (**Model 1**):

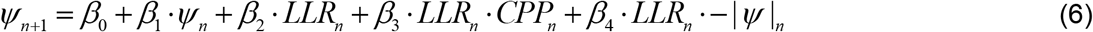

where *CPP_n_* was log-transformed to reduce skew, and both log(*CPP_n_*) and – *|ψ|_n_* were z-scored to reduce co-linearity between the interaction terms and *LLR_n_*. In this regression model, *CPP* and – *|ψ|* modulated the gain with which new evidence influenced the observer’s existing belief. Note that *β*_1_ and *β*_2_ fully captured the simple summation part of the normative accumulation rule (eq. 1). *β*_3_ and *β*_4_ thus approximated the non-linear component of the accumulation dynamics introduced by eq. 2. We assessed the contribution of each of the four terms to sample-wise belief updating by calculating their coefficients of partial determination (Figure 1g; Supplementary Figure 1):

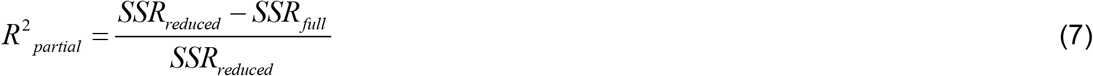

where *SSR_full_* was the sum of squared residuals of the full model in eq. 6 and *SSR_reduced_* was the sum of squared residuals of an otherwise identical model that excluded the term of interest.

We repeated the above analysis using the two levels of *H* used in Experiment 3 (0.1 and 0.9; Supplementary Figure 8a). Here we submitted *CPP* to the logit-transform rather than logtransform, since the latter was found to accentuate, rather than reduce, distribution skew in the high *H* condition. We also repeated the analysis with two alternative measures of surprise: ‘unconditional’ Shannon surprise calculated using only knowledge of the generative distributions (Spearman’s *ρ* with log(*CPP*)=0.00); and ‘conditional’ Shannon surprise calculated using both knowledge of the generative distributions and the observer’s existing belief state (Spearman’s *ρ* with log(*CPP*)=0.36). Definitions of these metrics and their modulatory effects on normative belief updating are reported in Supplementary Figure 10.

### Modelling human choice behavior

The similarity between our human participants’ choices from Experiment 1 and those of various candidate decision processes was evaluated in three ways. First, we computed the accuracy of the humans’ choices with respect to the true generative state at the end of each trial, and compared this to the accuracy yielded by three idealized decision processes presented with the same stimulus sequences as the humans (Figure 2a): the normative accumulation process for our task described above (with *H*=0.08), perfect accumulation of all *LLR*s (which is normative only for the special case of *H*=0), and basing one’s decision on the sign of the final evidence sample (normative for the special case of *H*=0.5). For each strategy and trial, choice *r* (*left*=-1, *right*=+1) was determined by the sign of the log-posterior odds after observing all samples: *r_trl_* = *sign* (*L_n,trl_*) for the normative rule, 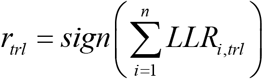 for perfect accumulation, and *r_trl_* = *sign* (*LLR_n,trl_*) for last sample only, where *n* indicated the number of samples presented on trial *trl*.

Second, for each strategy and human participant, we computed choice accuracy as a function of the duration of the final environmental state on each trial (i.e. the number of samples presented to the participant after the final change-point occurred; ranging from 1 on trials where a change-point occurred immediately before the final sample, to 12 on full-length trials where no change-points occurred; Supplementary Figure 2).

Third, we assessed the consistency between the humans’ choices and the choices generated by each idealized strategy by computing the slope of a psychometric function relating the strategy-specific log-posterior odds to human choice (Supplementary Figure 2). For each strategy and participant, we normalized log-posterior odds across trials and described the probability 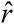 of a making a *right* choice on trial *trl* as:

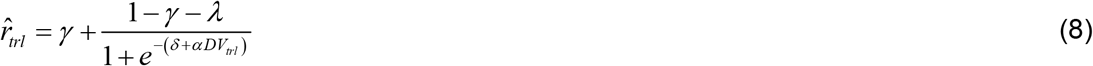

where *γ* and *λ* were lapse parameters, *δ* was a bias term, *DVr* was the z-scored log-posterior odds on trialwas computed as *DV_trl_*, and *α* was the slope parameter, which reflected the consistency between the choices produced by a given strategy and those of a human participant. We estimated *γ*, *λ*, *δ* and *α* by minimizing the negative log-likelihood of the data using the Nelder-Mead simplex search routine, under the constraint that the lapse rate was not dependent on choice (i.e. *γ* and *λ* were equal). Differences in *α* between candidate strategies were tested for via paired *t*-test.

#### Normative model fits

We also fitted variants of the normative model to the participants’ behavior. We assumed that choices were based on the subjective log-posterior odds *L_n,trl_* for the observed stimulus sequence on each trial *trl*, given eqs. 1–2. This per-trial variable was also corrupted by a noise term *v*, such that choice probability 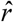 was computed as:

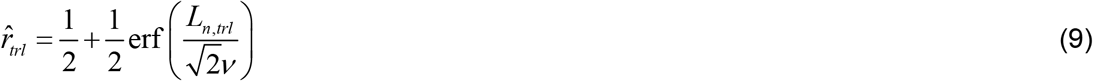

In addition to the noise term, we allowed for the possible presence of three further deviations of the participants’ away from the ideal observer: misestimation of the hazard rate, *H*; a bias in the mapping of stimulus location to *LLR*; and, a bias in the weighting of evidence samples that were (in)consistent with an existing belief state (see following section for motivation and details).

We fit the model to each participant’s data by minimizing the cross-entropy between human and model choices:

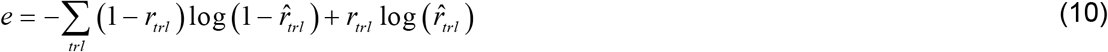

where *r_trl_* indicates the human participant’s choice on trial *trl* as above, and 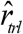 is the model’s choice probability calculated as per eq. 9. The sum of the cross-entropy *e* with any regularization penalty terms (see below) was minimized via particle swarm optimization^67^, setting wide bounds on all parameters and running 300 pseudorandomly-initialized particles for 1500 search iterations.

The relative goodness of fit of different model variants was assessed by calculating the Bayes Information Criterion (BIC):

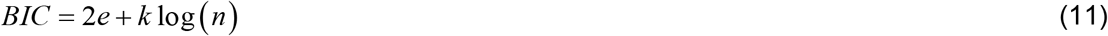

where *k* was the number of free parameters (see below), *n* was the number of trials and *e* was the cross-entropy as per eq. 10 (equivalent here to the negative log-likelihood of the data given the fitted model).

#### Motivation of constraints for normative model fits

The best-fitting version of the normative model to the data from Experiment 1 allowed for four deviations from the purely ideal observer. We motivate the inclusion of each of these here:

1. *Choice selection noise:* As shown in eq. 9, we applied a noise term only to the final log-posterior on a given trial (i.e. to the translation of final belief into a choice, after accumulation of all presented samples had taken place). We acknowledge that the accumulation process itself is likely susceptible noise, as reported for other tasks^68^. Accumulation noise is inherent in the cortical circuit model and an assumption underlying our analyses linking residual MEG and pupil fluctuations to choice (Figures 6, 7f, 8d). Nonetheless, we here used choice selection noise for simplicity, consistency with previous implementations of the normative model^16^, and because this yielded decent model fits. The exact locus and nature of the internal noise is beyond the scope of the present study.
2. *Misestimation of H:* In line with previous work on change-point tasks^16,69^, we allowed for participant-specific subjectivity in the hazard rate, *H*. Indeed, the model fits indicated that Experiment 1 participants had a tendency to underestimate *H* (subjective *H*=0.039 ± 0.005 (s.e.m.); *t*_16_=-7.7, *p*<10^-6^, two-tailed one-sample *t*-test of subjective *H* against true *H* of 0.08; Supplementary Figure 5b). This systematic bias could reflect prior expectations toward relative environmental stability at the stimulus presentation rate used here.
3. *Non-linear stimulus-to-LLR mapping:* In line with observations from other tasks^26^, we allowed for a non-linearity in the mapping of the decision-relevant stimulus dimension (polar angle) onto *LLR*. This was motivated by an analysis of our data in which we estimated the weight ascribed by participants to objective stimulus positions using an approach described elsewhere^2^. This analysis estimated the subjective weight of evidence associated with samples falling into evenly spaced bins (bin spacing=0.6 in true *LLR* space) using logistic regression:

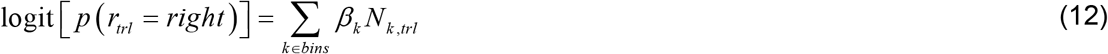

where *N_k,trl_* was the number of samples appearing on trial *trl* with a true *LLR* falling in bin *k*. Fitting this regression model to choices produced by both our human participants and the normative accumulation rule with matched noise revealed that human participants tended to give particularly strong weight to extreme samples (Supplementary Figure 5g). To account for this effect in our model fits without making assumptions about the shape of the participants’ weighting functions, we estimated the subject-specific mappings of stimulus polar angle to subjective *LLR* as a non-parametric function that was fit to the observers’ choices alongside the other free parameters described here. We expressed the subjective *LLR* as an interpolated function of stimulus polar angle *x* whereby values were estimated at *x*={12.5, 25, 37.5, 50, 62.5, 75, 82.5, 90}, we assumed symmetry around *x*=0 and *LLR*(*x*=0)=0, and interpolation was performed using cubic splines. For fitting, we used Tikhonov regularization of the first derivative of the function (thus promoting smoothness) by adding a penalty term to the objective function (see above): 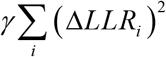, where *i* indexes the value of *x* for which a *i* subjective *LLR* was estimated and *γ* = 1/20 was determined through ad hoc methods^16^. These fits revealed that the participants tended to over-weight evidence samples in the extrema of the stimulus space, an effect that has been shown previously to countermand the decrease in accuracy incurred by noise.
4. *Bias in weighting of (in)consistent evidence:* Lastly, we allowed for the possibility that humans might give greater weight, relative to the normative accumulation process, to samples of evidence that were (in)consistent with their existing belief state^24,25^. To do so, we applied a multiplicative gain factor *g* selectively to *LLRs* associated with inconsistent samples, such that the effective evidence strength 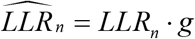 for any sample *n* where *sign* (*LLR_n_*)≠ *sign (ψ_n_*). Thus, *g* > 1 corresponds to a relative upweighting of inconsistent samples, while 1 > *g* ≥ 0 corresponds to a relative upweighting of consistent samples. We found that participants assigned higher weight than the normative model to samples that were inconsistent with their existing beliefs (fitted weight=1.40 ± 0.04 (s.e.m.); *t*_16_=8.2, *p*<10^-6^, twotailed one-sample *t*-test of fitted weights against normative weight of 1; Supplementary Figure 5d), perhaps reflecting constraints on the neural circuit implementation of non-linear evidence accumulation. In total, the full fits of the normative model plus bias terms to the data from Experiment 1 consisted of 11 free parameters per observer (1 noise term, 1 subjective *H*, 8 stimulus-to-*LLR* mapping parameters, and 1 gain factor on inconsistent samples). We also fit more constrained variants of the normative model that lacked various combinations of the deviations from the ideal observer described above (Supplementary Figure 4a). To facilitate fair comparison across model variants, for this set of analyses we re-parameterized the stimulus-to-*LLR* mapping function for each sample *n* as a scaled exponential:

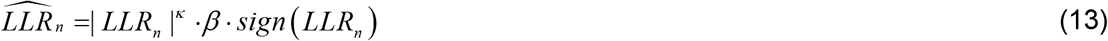

which was more constrained than the interpolated function in previous fits but can produce convex *(κ* > 1), concave (*κ* < 1) or linear (*κ* = 1) mapping functions using only two free parameters (exponent *κ* and scale parameter *β).*

We also fit several variants of the normative model to data from Experiment 3 (Supplementary Figure 8b), with the following differences from the above approach. First, we always let the subjective *H* parameter vary across the two objective *H* conditions. Second, because the nonlinear stimulus-to-*LLR* mapping yielded only a minor improvement in goodness of fit to the Experiment 1 data (Supplementary Figure 5a) and this mapping is expected to be invariant with respect to subjective *H*, we allowed for only a linear multiplicative scaling factor *β* to be applied to the true *LLRs* from Study 3 (i.e. 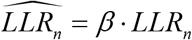); and we constrained this parameter to be fixed across *H* conditions. Third, because the normative non-linearity for the high (i.e. *H*=0.9) but not the low (i.e. *H*=0.1) *H* condition of Study 3 prescribes a sign flip in the transformation of *L*_*n*-1_ into *ψ_n_* (Figure 7b), this provides an opportunity to assess whether the consistency weighting parameter *g* is best defined relative the sign of *ψ_n_* or the sign of *L*_*n*-1_. We did so by comparing the goodness of fit of model variants assuming either definition. Fourth, we fit model variants which allowed us to assess whether choice selection noise or *g* might vary across *H* conditions (Supplementary Figure 8b).

#### Fitting the L to ψ mapping

To assess how well the critical non-linearity in the normative model (eq. 2) captured the belief updating dynamics of the human participants from Experiment 1, we also fit a model in which we estimated this function directly from the observers’ choice data without constraining its shape^16^ (Supplementary Figure 4c). As with the subjective stimulus-to-*LLR* mapping function above, we here estimated the mapping of *L*_*n*-1_ to *ψ_n_* as an interpolated non-parametric function. We assumed symmetry of the function around *L*_*n*-1_=0 and *ψ*(*L*_*n*-1_=0)=0, and *ψ_n_* was estimated for values of *L*_*n*-1_ that were spread evenly between 1 and 10 in steps of one. We applied Tikhonov regularization to the first derivative as described above, here with *γ* = 1/2. This model had a total of 20 free parameters, with the new mapping function replacing the subjective *H* parameter from the previous fits.

#### Alternative accumulator model fits

We additionally fit three alternative, sub-optimal accumulator models to the data from Experiment 1 that each lacked a key characteristic(s) of the normative accumulation process. The first of these was a perfect accumulator that linearly integrates all evidence samples without loss. This model is common in the literature but lacks both the non-linearity and leak components that are characteristic of the normative accumulation process^16^.

The second sub-optimal model was a linear approximation of the normative model that employed leaky accumulation but lacks the stabilizing, non-absorbing bounds in the normative *L* to *ψ* mapping. This linear model has been shown to be capable of approximating some operating regimes of the normative model^16,23^, and substitutes eq. 2 for the following *L* to *ψ* mapping:

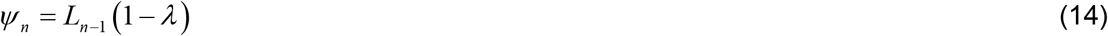

where *λ* is a free parameter reflecting the amount of leak.

The third sub-optimal model employed perfect, lossless evidence accumulation toward non-absorbing bounds. This model could capture the critical non-linearity in the normative *L* to *ψ* mapping, but lacks the leaky accumulation that is an important prescription of the normative model when beliefs are uncertain and *H* is high^16^. This model substitutes eq. 2 for the following:

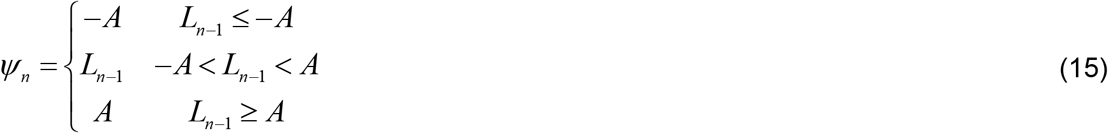

where *A* is a free parameter reflecting the height of the non-absorbing bounds.

Our fits of the three alternative models variously incorporated noise, non-linear *LLR* mapping and an inconsistency bias in the same ways as described above (Supplementary Figure 5). Thus, with *λ* or *A* replacing subjective *H*, the latter two models had the same number of free parameters as our fits of the normative model plus bias terms, while the unbounded perfect accumulator had one fewer.

#### Psychophysical kernels

We estimated ‘psychophysical kernels’, which quantify the impact (regression weight) of evidence on choice as a function of time. To do so, we fit variants of the following logistic regression model to choices on full-length trials (**Model 2**):

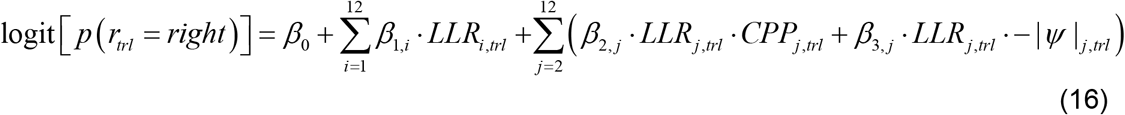

where *i* and *j* indexed sample position within the stimulus sequence on trial *trl* (12 sample positions for Experiments 1 and 2, 10 sample positions for Experiment 3), and *LLR* was the true *LLRs* given the generative task statistics (i.e. not subjected to the non-linearity or inconsistency bias in the model fits). For Experiment 3, we excluded the – |*ψ*| modulatory terms because they did not contribute to normative belief updating in this generative setting (Supplementary Figure 8a). For the humans the dependent variable was the empirically observed choice *(left=0, right=1);* for the models it was the choice probability calculated as per eq. 9. The set of coefficients *β*1 estimated the time-dependent leverage of evidence on choice. The additional set of interaction terms *β*_2_ and *β*_3_ estimated the modulation of evidence weighting by change-point probability and uncertainty, respectively. As in **Model 1** (eq. 6), *CPP* was log-transformed (or logit in the case of Experiment 3, see above), and both transformed *CPP* and –*|ψ|* were z-scored before multiplication with *LLR* to reduce collinearity. Additionally, all final regressors were z-scored across the trial dimension to yield fitted coefficients on the same scale. Cluster-based permutation testing^70^ was used to identify sample positions for each of the three sets of terms at which fitted weights differed significantly from zero (two-tailed one-sample *t*-test; 10,000 permutations; cluster-forming threshold of *p*<0.05). In the high *H* condition of Experiment 3, kernels were expected to exhibit a damped oscillation (Figure 7d); for the statistical testing only, we therefore flipped the sign of the regression coefficients at every odd sample position to ensure a fair comparison. Differences between the *CPP* and –*|ψ|* weights were tested via paired *t*-test after averaging weights over sample position.

#### Circuit modelling

We simulated the choice behavior of an established cortical circuit model for decision-making^20^ on the variant of our task used in Experiment 1. The model consisted of 1600 pyramidal cells and 400 inhibitory interneurons, all of which were spiking neurons with multiple different conductances (see below). The pyramidal cells were organized into three distinct populations: 240 neurons selective for the ‘left’ choice (population *D*_1_), 240 neurons selective for the ‘right’ choice (*D*_2_), and the remaining neurons non-selective. Neurons within the choice-selective populations sent recurrent connections to neurons in the same population as well as to a common pool of inhibitory interneurons (*I*), which fed back onto both *D*_1_ and *D*_2_. Pyramidal neurons projected to AMPA and NMDA receptors (with fast and slow time constants, respectively) on target cells, and interneurons projected to GABA_A_ receptors. The parameterization and update equations of the circuit model were taken from their original description in ref. ^20^, except for three changes described in the following.

First, in the original implementation of the model, stimulus-driven inputs to the choice-selective populations varied linearly with stimulus strength, were symmetric around 40 Hz, and always summed to 80 Hz. Thus, both choice-selective populations received average input of 40 Hz when stimulus strength was zero, while one received 80 Hz and the other 0 Hz at maximal stimulus strength. This stimulus input function produced excessive primacy in evidence weighting, inconsistent with the data. Stimulus inputs (sample-wise *LLR*) needed to be stronger for changes-of-mind to occur in response to inconsistent evidence (see also ref. ^20^). Thus, we used a threshold-linear input function that was also symmetric around 40 Hz for the two choice-selective populations, but imposed no upper bound on the input:

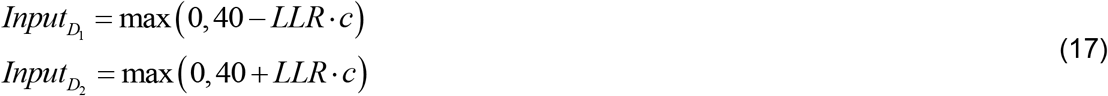

where *Input_x_* was the input to choice-selective population *x*, and *c* was a multiplicative scaling factor applied to the sample-wise *LLR*. We set *c* to 19, yielding an input of ~110 Hz to the favored choice-selective population for the strongest possible stimulus (and 0 Hz to the other population). We verified that strong stimulus input, rather than the threshold-linear function itself, was the key factor for approximating normative decision-making in our task: An input function that was again symmetric around 40 Hz, but had different slopes for *LLR*<0 and *LLR*>0 and produced an input of 0 Hz to the non-favored population only for the most extreme evidence strength, yielded very similar behavior as what we report here (data not shown; see ref.^20^ for further examination of this form of input function).

Second, the recurrent connectivity between pyramidal cells in the model was structured to enforce stronger coupling between cells within the *same* choice-selective population than between cells in *different* choice-selective populations. Specifically, within a choice-selective population, *w_j_*=*w*_+_, where *w*_+_>1 determined the strength of ‘potentiated’ synapses relative to the baseline level of potentiation. Between two different choice-selective populations, and from the non-selective population to the selective ones, *w_j_*=*w_-_*, where *w*_-_ <1 determined the relative strength of synaptic depression. *w*_-_ was directly determined by *w*_+_ such that the overall recurrent excitatory synaptic drive without external stimulus input remained constant as a function of *w*_+_ is varied^20^. In the original implementation of the model *w*_+_=1.7, which produced relatively strong and stable attractor states that, as with weakly scaled stimulus input, prohibited changes-of-mind in response to inconsistent evidence for all but the strongest evidence strengths. We thus set *w*_+_=1.68, with the resulting mildly weakened attractor dynamics allowing the model to better approximate normative decision-making on our task (stronger recency, and increased sensitivity to *CPP*).

Third, in the original model implementation simulations were run with an integration time step *dt*=0.02 ms. Because we needed to simulate ~25,000 trials of 5.6 s each and the model is slow to simulate, we set *dt*=0.2ms. We verified through simulations at the original *dt* that this did not significantly change the behavior or population rate trajectories of the model.

We ran simulations of the model for all full-length (12-sample) trials presented to our human participants. The network was initialized at 0.4 s before the onset of the first evidence sample and the simulation ended 0.4 s after offset of the final evidence sample. Each evidence sample was assumed to provide external input to the choice-selective populations according to the stimulus input function described above, from the time of its onset to 0.4 s thereafter; during the 0.4 s periods before and after the sample sequence, external input was set to zero. The instantaneous mean population firing rates of the choice-selective populations, *r*_*D*_1__ and *r*_*D*_2__, were calculated by summing all spikes across a population within a 50 ms window centered on the time-point of interest, and dividing by the number of neurons in the population and the time window. The evolving model decision variable *X* was defined as the difference between the instantaneous firing rates of the two populations: 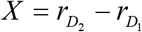. The model’s choice was determined by *sign*(*X*) at the end of each simulated trial; thus, like the human participants, the model needed to maintain a memory of its decision state during a post-evidence sequence delay period without external input. The model’s updated decision variable given a new evidence sample was taken to be *X* at 0.4 s after onset of that sample.

We estimated the model’s psychophysical kernels as per eq. 16, using *CPP* and –*|ψ|* metrics from the ideal observer variant of the normative model. Because our goal was not to fit the circuit model to the data (which cannot be achieved in a principled fashion, due to the large number of parameters), but rather to test whether the circuit model can reproduce the key *qualitative* features of our behavioural and neural data, we did not manipulate the internal noise level of the circuit model. As a consequence, internal noise in our implementation was low relative to the strength of the sensory input, and the circuit model kernels were thus of larger magnitude relative to those of the human participants and normative model fits. In order to facilitate comparison of the qualitative features of model and behavioral data, we applied a single multiplicative scaling factor (0.375) to all three kernels produced by the circuit model, which approximately matched kernel magnitudes with the human participants while preserving relative differences between the kernel types. This scaling factor was chosen manually.

We also simulated the choice behavior of a reduction of the above biophysical circuit model that was described by the diffusion of a decision variable *X* in the double-well potential *φ*:

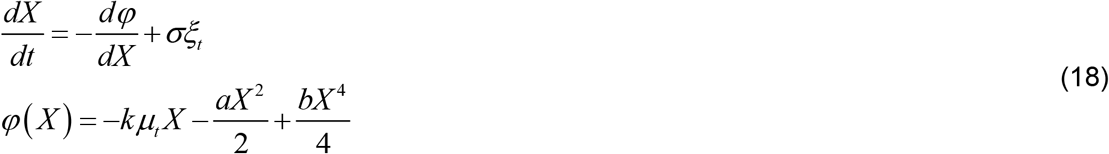

where *μ_t_* was the differential stimulus input to the choice-selective populations at time point *t* relative to trial onset (in our case the per-sample *LLR,* which changed every 0.4 s) that was linearly scaled by parameter *k; ξ_t_* was a zero-mean, unit-variance Gaussian noise term that was linearly scaled by parameter *σ,* and *a* and *b* shape the potential^71^. The model was initialized on each trial at the onset of the first sample with *X* set to 0, simulated with time-step *dt* set to 25 ms, and its choice was determined by *sign(X)* at 0.4 s after the onset of the final sample in each sequence. As with the detailed spiking neuron model, we fed the reduced model the stimulus sequences for all full-length trials presented to our human participants, and calculated psychophysical kernels as per eq. 16.

The reduced model helped us to explore boundary conditions for approximating human behavior. We manually explored the parameter space to find a set of parameters at which the reduced model produced psychophysical kernels that approximately matched those of our human participants (*k*=2.2, *σ*=0.8, *a*=2, *b*=1). The shape of the potential corresponding to this combination of *a* and *b* indicated weak bi-stable attractors (Supplementary Figure 6, top). Keeping *k* and *σ* constant, we then simulated the behavior of the model for three qualitatively different dynamical regimes: a single attractor centered on *X*=0 (*a*=0, *b*=1), which produced extreme recency in evidence weighting; perfect integration (*a*=0, *b*=0), which produced a flat psychophysical kernel and no sensitivity to *CPP* or –*|ψ|*; and strong winner-take-all dynamics (*a*=5, *b*=1), which produced extreme primacy in evidence weighting (Supplementary Figure 6).

### Data acquisition

MEG data were acquired for Experiments 1 and 3 on a whole-head CTF system (275 axial gradiometer sensors, CTF Systems, Inc.) in a magnetically shielded room at a sampling rate of 1,200Hz. The location of the participant’s head was recorded and visualized in real-time using three fiducial coils, one fixed at the outer part of each ear canal and one on the nasal bridge. Participants were instructed to minimize movement during task performance. A template head position was registered at the beginning of each participants’ first session, and the experimenter guided the participant back into that position before initializing each task block.

We used Ag/AgCl electrodes to measure the electrocardiolgram (ECG), vertical electrooculogram (vEOG), and electroencephalogram (EEG) from three scalp locations (Fz, Cz and Pz according to the 10/20 system) with a nasion reference, though these data are not analyzed here. Eye movements and pupil diameter were recorded during task performance at 1,000Hz using an MEG-compatible Eyelink 1000 Long Range Mount system (SR Research).

T1-weighted structural magnetic resonance images (MRIs) were acquired from Experiment 1/3 subjects to generate individual head models for source reconstruction (see below).

### MEG data analysis

#### Preprocessing

MEG data were analyzed in MATLAB (MathWorks) and Python using a combination of the Fieldtrip toolbox^72^, MNE^73^ and custom-made scripts.

Continuous data were first segmented into task blocks, high-pass filtered (zero-phase, forward-pass FIR) at 0.5Hz, and bandstop filtered (two-pass Butterworth) around 50, 100 and 150 Hz to remove line noise. For Experiment 1, data were then resampled to 400 Hz and re-segmented into single trials with the following task-dependent timings: from 1 s before trial onset to the onset of the ‘Go’ cue for the decision-making task; and from 1 s before to 2 s after lexical cue onset for the motor localizer task. Trials containing any of the following artifacts were discarded from further analysis: (i) head motion of any of the three fiducial coils exceeding a translation of 6 mm from the first trial of the recording; (ii) blinks (detected using the standard Eyelink algorithm); (iii) saccades (detected with velocity threshold=30°s^-1^, acceleration threshold=2000°s^-2^) exceeding 1.5° in magnitude; (iv) squid jumps (detected by applying Grubb’s test for outliers to the intercepts of lines fitted to single-trial/-sensor log-power spectra from data without a high-pass filter); (v) sensor(s) with a min-max data range exceeding 7.5 pT, usually caused by cars driving past the MEG laboratory; (vi) muscle artifacts (detected by applying a 110-140 Hz Butterworth filter to all MEG sensors, z-scoring across time, and applying a threshold of *z*=20 to each sensor).

Due to a moderate increase in the number of observed blink artifacts, we adopted a different approach to preprocessing data from Experiment 3 (decision-making task only). Motion, squid jump, min-max (within 2 s windows) and muscle artifacts were identified as above, but now in the continuous time series for single MEG recordings (typically consisting of four task blocks, excluding data recorded during between-block breaks). The remaining data for that recording

were then subjected to temporal independent component analysis (ICA) using the infomax algorithm, and independent components capturing stereotyped artifacts caused by blinks and heart beats were identified by visual inspection and removed. The resulting cleaned data were then segmented into single trials, from 1 s before trial onset to the onset of the ‘Go’ cue.

#### Spectral analysis

In order to isolate induced (non-phase-locked) activity components from the activity of each sensor, we first subtracted each sensor’s trial-averaged (‘phase-locked’) response from its single-trial activity time courses. Thus, results of all spectral analyses reported in this paper reflect modulations of cortical activity that are not phase-locked to external events, but rather are likely generated by recurrent synaptic interactions in cortical circuits^58^ (see also *Discussion).* Repeating the analyses on the ‘raw’ signals (i.e. including both phase-locked and non-phase-locked activity components) did not reveal any additional features (data not shown). This likely reflects the fact that the computation of all variables assessed here entailed recurrent interactions in cortical micro- and macro-circuits^58^. Note that while phase-locked responses to single samples in early visual cortex were present in our data, this does not imply that these responses encode the to-be-accumulated evidence for the decision process; rather, deriving decision-relevant evidence (*LLR*) from sensory responses encoding polar angle required a transformation that incorporates knowledge of the stimulus statistics in our task.

We used sliding-window Fourier transform to compute time-frequency representations (TFRs) of the single-trial activity of each MEG sensor from both the decision-making and motor preparation tasks. Specifically, for low frequencies (1-35 Hz in steps of 1 Hz), we used a sliding-window Fourier transform with one Hanning taper (window length of 0.4s in steps of 0.05s; frequency smoothing of 2.5 Hz). For high frequencies (36-100 Hz in steps of 4 Hz), we used the multi-taper method with a sequence of discrete proloid slepian tapers, a window length of 0.25 s in steps of 0.05 s, and 6 Hz frequency smoothing. We converted the complex-valued time-frequency representations of activity into units of power by taking the absolute values and squaring.

For sensor-level analyses, the axial gradiometer data were decomposed into horizontal and vertical planar gradients prior to time-frequency decomposition and these were combined afterwards to yield readily interpretable topographies.

We converted power estimates into units of modulation relative to pre-trial baseline as follows. Power estimates for each time-point *t*, frequency *f* and sensor *c* were normalized and baseline-corrected via the decibel (dB) transform *dB_t,f,c_*=10*log_10_(*power_t,f,c_/baseline_f,c_*). Here, *baseline_f,c_* refers to the trial-averaged power collapsed across the interval from −0.4 to −0.2 s relative to trial onset. This was done to normalize (via the log-transform entailed in the above equation) the power estimates for linear regression analyses reported below.

#### Construction of linear filters for motor preparatory activity

We used data from the Experiment 1 delayed motor response task (‘motor localizer’) to construct a set of filters for isolating hand movement-specific motor preparatory activity in the data from the decision-making task. Those filters are referred to as ‘motor filters’ in the following.

First, any motor localizer trial on which a manual response was executed before the ‘Go’ cue was excluded from analysis. This was the case on the majority of trials for 5 participants who apparently misunderstood the task instructions, and their data were not used for construction of the filters. Motor localizer MEG datasets from the remaining 12 participants were segregated into sensors covering the left side of the head (labelled ‘ML*’ by the CTF system) and those covering the right side of the head (‘MR*’). For each matching pair of left/right sensors (e.g. left frontal sensor ‘MLF45’ matches with right frontal sensor ‘MRF45’; 131 pairs in total), we calculated a lateralization index *LI* for each time-point, frequency and trial by subtracting the power modulation estimate *dB_t,f,c_* for the left sensor from the power estimate for the right sensor. We then fit the following linear regression model to each participant’s data:

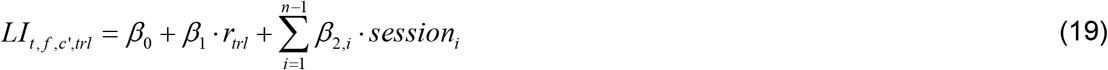

where *t* indicated time-point relative to trial onset, *f* indicated frequency, *c’* indicated sensor pair, *trl* indicated trial, *r_trl_* was the single-trial response executed by the participant *(left=0, right=1),* and *session_i_*, was a group of binary nuisance regressors included to absorb any main effect of experimental session (with *n* denoting the total number of sessions for a given participant).

The quantity of interest was the t-score associated with *β*_1_, which provided a reliability-weighted measure of the strength with which *LI* encodes the motor response. We averaged these t-scores across the interval from 0.7 to 1.1 s post-lexical cue to generate a single sensor*frequency t-map per participant. This interval captured the period of the task during which planning, but not execution, of the motor response takes place. We then used cluster-based permutation testing (10,000 permutations with cluster-forming threshold of *p*<0.01, two-tailed) to identify spatio-spectral clusters that were significantly different from zero at the group level. This procedure yielded a single cluster (*p*<0.001; *p*>0.57 for all other clusters) with an associated sensor*frequency matrix *M* of group-level t-scores where all spatio-spectral points lying outside the cluster bounds were set to zero. The matrix *M* was used to construct the motor filters.

We generated three sets of motor filters: spectral filters, spatial filters, and spatio-spectral filters. To generate weights *w_f_* for a *spectral* filter, we integrated *M* over the spatial dimension such that:

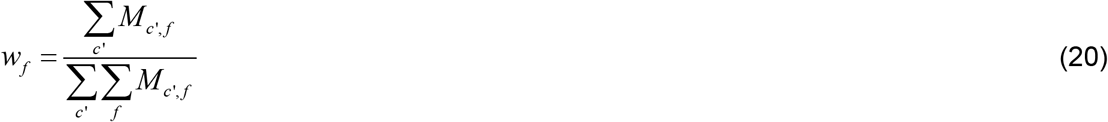

where the denominator normalized the weights to integrate to one. Weights for *spatial* and *spatio-spectral* filters were generated analogously by integrating over the spatial dimension in the numerator or not at all. The group-average spectral and spatial filters are shown in Figure 3a (right).

The resulting filters could then be applied to independent *LI* data by computing the dot product between data and filter along the desired dimension(s), yielding a filtered motor preparatory signal that we refer to as *motor* below. For example, the spectral filter computed through eq. 20 could be applied to yield a spatially-resolved motor preparatory signal as follows:

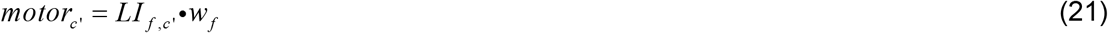

#### Model-based analysis of motor preparatory activity during decision-making task

We applied the movement-selective motor preparation filters to sensor-level *LI* data from fulllength (12-sample) trials of the Experiment 1 decision-making task in the manner described above. Application of the *spatial* filter to these data in a time-resolved fashion generated TFRs of the relative motor preparation for each choice alternative during decision formation (Figure 3b). These were then segmented from 0 to 1s relative to the onset of each of the 12 samples per trial to yield a four-dimensional matrix (time*frequency*sample*trial), through which we assessed the sensitivity of the motor preparatory signal to decision-relevant computational variables.

It became apparent that fits of the normative model that allowed for non-linear stimulus-to-*LLR* mappings invariably yielded computational variables (in particular, *LLR*, *CPP* and *L*) with a small proportion of highly deviant values. This was caused by the tendency of human observers to assign especially strong weight to infrequently-encountered stimuli at the extrema of the stimulus range (Supplementary Figure 5). These outliers were in turn problematic for our analyses relating model-derived variables to neural measurements, the majority of which relied on linear regression. For all such analyses, we therefore derived computational variables from model fits in which the stimulus-to-*LLR* mapping was constrained to be linear (such that 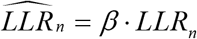, where *β* sets the slope of the mapping function and is a free parameter). Although this model variant yielded marginally worse goodness-of-fit to observers’ choices compared to the full model (Supplementary Figure 5), the sample-wise computational variables generated by each were highly correlated (*ψ*: *ρ* = 0.988 ± 0.014; *L: ρ* = 0.988 ± 0.015; *CPP*: *ρ* = 0.986 ± 0.018; – *|ψ|*: *ρ* = 0.959 ± 0.045).

To determine the sensitivity of movement-selective motor preparation to each of the key components of normative belief updating established previously (Figure 1f; **Model 1**), we fit the following linear model (**Model 3**):

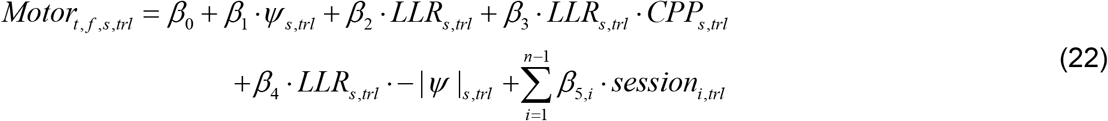

where *motor_t,f,s,trl_* was the motor preparatory activity (see above) at time *t* relative to sample onset, frequency *f*, sample *s* and trial *trl*. We averaged *β*_1-4_ across the sample dimension, thereby obtaining single time-frequency maps per participant reflecting the strength with which each computational quantity is encoded in the varying motor preparation signal (Figure 3g). Clusters of significant encoding across time and frequency in the sample-averaged maps were identified via cluster-based permutation test (two-tailed one-sample *t*-test; 10,000 permutations, cluster-forming threshold of *p*<0.05), as were significant differences between the *CPP* and – *|ψ|* interaction terms (two-tailed paired *t*-test; same threshold and permutation number).

To complement the above analysis, we constructed two regression models, each of which contain complementary parts of the ‘full’ model from eq. 22. The idea was to fit neural signals that encoded either new evidence (*LLRs*), or the non-linearly updated decision variable (*ψ_s+1_*), associated with sample *s*. We then fit these models to *motor_t,f,s,trl_* on full-length trials (**Models 4-5**):

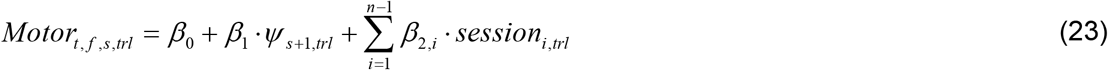

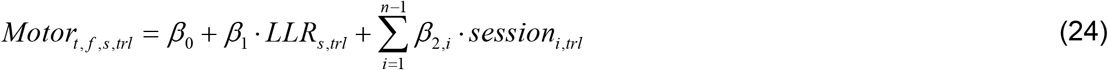

For each model, time-point *t*, frequency *f* and sample *s*, we computed a ‘super-BIC’ score reflecting that model’s goodness-of-fit at the group level. This metric was calculated as per eq. 11, here with the negative-log-likelihood, number of free parameters and number of observations determined by the sum of these values across participants. We then averaged across the sample dimension and subtracted BIC scores for the ‘belief model’ (eq. 23) from those for the ‘evidence model’ (eq. 24). This generated a single time-frequency map of relative goodness-of-fit in which positive values indicate stronger encoding of belief relative to sensory evidence in the motor preparation signal, and *vice versa* for negative values (Figure 3f). We further compared the belief model of eq. 23 to an alternative in which the non-linearly transformed decision variable after accumulating sample *s* (*ψ_s+1_*) was replaced by the untransformed posterior log-odds (*L_s_*). The same model comparison approach described above revealed that *ψ* was the superior predictor of the motor preparation signal (data not shown), as expected given the observed modulation of the motor preparation signal by change-point probability and uncertainty (Figure 3g).

We also characterized the shape of the relationship between *motor_t,f,s,trl_* and model-derived belief state as follows. We applied the *spatio-spectral* motor preparation filter to the decisionmaking task data to yield a single scalar value per time point reflecting the relative motor preparation for each alternative (Figure 3c). This metric was then averaged from 0.4-0.6s after each sample *s* on full-length trials (thus capturing latencies at which the signal is modulated by *CPP* while minimizing contamination by responses to subsequent samples; Figure 3g), and *z*-scored across trials separately for each sample position and session. We also normalized the posterior log-odds *L_s_* in an analogous fashion to remove any differences in the range of *L* across participant-specific model fits. Next, for each sample position and participant we sorted the normalized motor preparation by normalized *L_s_* into 11 equal-sized bins, and calculated the mean of both metrics per bin *b*. We then averaged across sample positions and participants to yield a single *belief encoding function* for the motor preparation signal which, for visualization, we rescaled with sign preservation to have upper and lower bounds of +1 and −1, respectively (Figure 3d).

In order to also estimate corresponding function for the normative model, we repeated the above procedure after replacing the motor preparation signal with *ψ_s+1_* (Figure 3d). The resulting function will generate the *L*-to-*ψ* mapping for the normative model described by eq. 2, after application of the normalization, binning and averaging process described above. We further repeated the procedure using the difference in firing rates between choice-selective populations in the biophysical model, the variable reflecting this model’s decision variable (see above). For this analysis we extracted difference measures at 0.4s after each sample onset.

We also quantified the strength of the correlations between the motor preparatory signal and model-derived decision variables at the single-trial level (Figure 3e). For each full-length trial we extracted scalar values of the relevant variables for each sample position *s* (*motor*: averaged from 0.4-0.6s after sample onset; normative model: the per-sample *L*; circuit model: difference in population rates 0.4s after sample onset) and quantified the correlation over sample positions between the motor signal and each model variable via Pearson correlation. For each participant we tested whether the distribution of single-trial correlation coefficients was different from zero via two-tailed, one-sample permutation test. We tested for a corresponding group-level effect by taking the median of the single-trial correlation coefficients for each participant and testing whether the group distribution of medians was different from zero, again via two-tailed permutation test.

#### Source reconstruction

We used linearly constrained minimum variance (LCMV) beamforming to estimate activity time courses at the level of cortical sources^74^. We first constructed individual three-layer head models from subject-specific MRI scans using Fieldtrip^72^ (functions: ft_volumesegment and ft_prepare_mesh; 4 of 30 Experiment 3 participants lacked MRI scans, in which case we used the Freesurfer average subject). Second, head models were aligned to the MEG data by a transformation matrix that aligned the average fiducial coil position in the MEG data and the corresponding locations in each head model. Transformation matrices were generated using the MNE software package, and we computed one transformation matrix per recording session. Third, we reconstructed cortical surfaces from individual MRIs using FreeSurfer^75^,^76^ and aligned two different atlases to each surface (see *Functional ROIs* below). In a fourth step we used the MNE package to compute LCMV filters for projecting data into source space. LCMV filters combined a forward model based on the head model and a source space constrained to the cortical sheet (4096 vertices per hemisphere, recursively subdivided octahedron) with a data covariance matrix estimated from the cleaned and segmented data. We computed one filter per vertex, based on the covariance matrix computed on the timepoints from trial onset until 6.2 s later (Experiment 1; 5.4 s for Experiment 3) across all trials. We chose the source orientation with maximum output source power at each cortical location. In a final step, we projected the broadband single-trial time series, as well as the complex-valued spectrograms (computed by the same spectral analysis approach described above) into source space. In source space we computed TFR power at each vertex location by first aligning the polarity of time-series at neighboring vertices (because the beamformer output potentially included arbitrary sign flips for different vertices) and then converting the complex Fourier coefficients in each vertex into power (taking absolute value and squaring).

We finally averaged the estimated power values across all vertices within a given ROI (see *Functional ROIs* below). As with the sensor-level analysis, source-power estimates were baseline-corrected using the dB transform (from −0.4 to −0.2 s relative to trial onset). We then computed source-level lateralization indices *LI*_t,f,trl,roi_ for each time-point, frequency, trial and ROI by subtracting the power estimate for the left hemisphere ROI from the power estimate for the right hemisphere ROI.

The code that produced source estimates is available at www.github.com/Donnerlab/pymeg.

#### Regions of interests

We focused on a total of 31 regions of interest (ROIs), which were delineated in previous functional MRI work^47,77,78^ and likely participated in the visuo-motor transformation that mapped patch locations to behavioral choice in our task. These regions were composed of (i) retinotopically organized visual cortical field maps provided by the atlas from Wang et al.^77^ and (ii) three regions exhibiting hand movement-specific lateralization of cortical activity: aIPS, IPS/PCeS and the hand sub-region of M1^47^; and (iii) a dorsal/ventral premotor cortex cluster of regions from a whole-cortex atlas^78^.

Following a scheme proposed by Wandell and colleagues^79^, we grouped visual cortical field maps with a shared foveal representation into clusters (see Table 1), thus increasing the spatial distance between ROI centers and minimizing the risk of signal^74^ (due to limited filter resolution or volume conduction). ROI masks from both atlases^77,78^ as well as ref. ^47^ were coregistered to individual MRIs.

**Table 1:** ROI definition along the sensorimotor pathway.

| Cluster | Functional areas | Source |
| --- | --- | --- |
| V1 | Dorsal and ventral parts of V1 | Wang et al., 2015 |
| V2-V4 | Dorsal and ventral parts of V2, V3, V4 |  |
| V3A/B | V3A, V3B |  |
| **IPS0/1** | IPS0, IPS1 |  |
| **IPS2/3** | IPS2, IPS3 |  |
| **LO1/2** | LO1, LO2 |  |
| **MT+** | TO1, TO2 |  |
| **VO1/2** | V01, V02 |  |
| **PHC** | PHC1, PHC2 |  |
| **aIPS** | aIPS1 | de Gee et al., 2017 |
| **IPS/PCeS** | IPS/Post-central sulcus |  |
| **M1 (hand)** | M1 |  |
| **PMd/v** | 55b, 6d, 6a, FEF, 6v, 6r, PEF | Glasser et al., 2016 |

#### Model-based analysis of different ROIs during decision-making task

At the level of these functional ROIs, we repeated the model-based regression analyses described above for the sensor-level motor preparatory activity (see *Analysis of motor preparatory activity during decision-making task).* Here, we replaced the sensor-level motor preparation signal *motor_t,f,s,tri_* with source-level hemispheric lateralization indices (*LI_t,f,s,trl_*) for each ROI. We fitted the linear regression model comprising each of the key components of normative belief updating (**Model 3**; eq. 22) to the data from Experiment 1 for each ROI. We plotted time-frequency maps of the group-level t-scores reflecting sample-wise encoding of *LLR* (*β*_2_), averaged over sample positions, to initially highlight qualitative differences between individual ROIs (Figure 4a), and averaged within early visual (V1-V4), dorsal/ventral visual (V3A/B, IPS0-3, LOC1/2’ MT+, Ventral occipital, PHC) and ‘decision-encoding’ (IPS/PCeS, PMd/v, M1) sets for the remaining terms (Figure 4b).

The above fits of **Model 3** revealed that signatures of the normative decision variable were widely distributed across cortex in the alpha frequency band (8-14 Hz). We then dissected the nature of this decision-related alpha-band activity in more detail. For a set of early/dorsal visual, intraparietal and motor regions which we assumed were ordered hierarchically^36^ (V1, V2-4, V3A/B, IPS0/1, IPS2/3, IPS/PCeS, PMd/v, M1), we fit the linear model described by eq. 22 to *LI* data for each ROI segmented from 0 to 1.4s around sample onset, extracted fitted *β*_2_ coefficients, and averaged across sample positions and 8-14 Hz to yield a vector reflecting the temporal evolution of alpha-band *LLR* encoding. We flipped the sign of this vector such that the peak encoding was positive, interpolated to millisecond temporal resolution using cubic splines, and normalized such that the maximum of the vector was equal to one (normalized vector 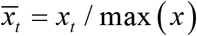; Figure 5f). We then calculated the latency to half-maximum of the vector for each ROI (Figure 5e), as well as the timescale *τ* of the *LLR* encoding by fitting a decaying exponential:

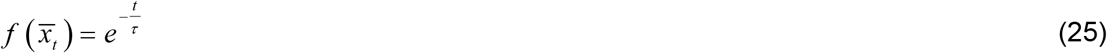

to the right portion of the peak-aligned vector (Figure 5g-h; fit by minimizing the sum of squared residuals between vector and fit via simplex).

We averaged latency and timescale estimates over ROIs within three sets (V1; V2-IPS3; IPS/PCeS, PMd/v and M1), and used two-tailed weighted permutation tests (10,000 permutations) to test for parameter differences between pairs of ROI sets. For each such test, each participant *p* was assigned a weight *W_p_* that was determined by the minimum strength of their time-averaged alpha-band *LLR* encoding across a given ROI pair. Any participant with a negative average *LLR* encoding for either ROI set of the pair was assigned *wp*=0 for that test, and weights were constructed such that 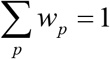. *w_p_* determined both the contribution of *P* each participant to the weighted mean over participants and the probability that they would be drawn, with replacement, in each iteration of the permutation test. This procedure downweighted the contribution of participants with weak/absent *LLR* encoding effects, for whom latency and timescale estimates were much more susceptible to noise in the *LLR* encoding traces.

We also fitted the regression model in **Model 3** (eq. 22) to the source-reconstructed data from Study 3, separately for each *H* condition. This allowed us to assess whether the strength of *LLR* encoding in the alpha and gamma frequency bands depended on the level of environmental volatility. For alpha (gamma), we averaged *β*_2_ over sample positions, 8-14 (40-65) Hz and 0.25-0.65 (0.05-0.45) s, with the specified time window covering the peak *LLR* encoding at that frequency band averaged over *H* conditions (not shown). We then tested whether there was a significant difference in the resulting scalar estimates of *LLR* encoding strength across *H* conditions (Figure 7e) via two-tailed permutation test (10,000 permutations). We also tested whether the direction of this *H* effect differed between the two frequency bands via two-tailed sign test.

Spatial gradients in the estimated parameters across the visual cortical hierarchy (V1, V2-V4, V3A/B, IPS0/1, IPS2/3; Figure 5h) were also tested for via two-tailed weighted permutation test, here fitting a line to the hierarchically-ordered parameters and testing if the slope of the fitted line differed from zero.

#### Recovery of latency differences via simulation

The latency differences in alpha-band *LLR* encoding that we identified between groups of ROIs (Figure 5e) were on the order of approximately 50-100 ms. We performed simulations to ensure that our spectral analysis approach is sensitive to latency differences of this magnitude. We generated a synthetic 10 Hz oscillation (approximating the frequency band of the candidate feedback effects observed in the data) and subjected this signal to a transient amplitude modulation, using the average V1 *LLR* encoding time course (8-14 Hz band) as a template. We assumed the amplitude modulation to be a decrease in power as we see for the data in the 8-14 Hz range (the results generalize to power increase). We then increased the latency of this amplitude modulation across a range from 10 to 200 ms and, for each step, computed a latency effect by comparing the recovered latency (see below) to the ‘baseline’ latency of the original V1 amplitude modulation according the following procedure. We, i) simulated 1000 ‘trials’ of 10 Hz signals, with phases randomized over trials; ii) added pink noise of varying amplitude to the signals; iii) computed 10 Hz power of the single-trial traces; iv) averaged power traces over trials and normalized the average trace to be positive-going and have a peak of 1; and v) computed the latency to half-maximum of this normalized trace. This procedure simulated physiologically plausible signals (pink noise and randomized oscillation phases) and entailed the exact same transformations we applied to the data in the respective analyses. This exercise revealed that our spectral analysis approach is well capable of detecting latency effects, even as small as 10 ms and for a high level of noise (data not shown).

#### Decoding of LLR from evoked responses

We used multivariate pattern classification techniques to decode *LLR* from the estimated evoked response (also known as ‘event-related field’) activity patterns of individual ROIs from the Experiment 1 data. Specifically, we used ridge regression to predict per-sample *LLR* using the evoked responses from all vertices across both cerebral hemispheres, separately for each ROI within the subset that was the focus of the majority of analyses above (V1, V2-4, V3A/B, IPS0/1, IPS2/3, IPS/PCeS, PMd/v, M1). This analysis was carried out on the raw data, without subtracting out the trial-averaged response as for the spectral analyses above. We first low-pass filtered the single-trial source-reconstructed signal to 40 Hz to remove high-frequency noise, sub-sampled the data from 400 to 200 Hz for computational efficiency, segmented from −0.1 to 1.4 s around individual sample onset, and baseline-corrected the signal at each vertex by subtracting the mean signal across the 0.075 s preceding sample onset. For each subject, ROI, sample position and peri-sample time-point, per-vertex signal values were z-scored across trials based on the training set (with the same transformation also applied to the test set before evaluating prediction performance, see below). We then performed dimensionality reduction, computing principle components of the training set, keeping all components that cumulatively explained 95% of training set variance, and projecting training and test set data into the space defined by these components. We fitted the decoder using 10-fold crossvalidation, and an L2 regularization term of *α*=10 determined by *ad hoc* methods (verifying that alternative *α* values of 0.1, 1 or 100 had no discernible effect on prediction performance for a subset of participants). We evaluated prediction performance by computing the Pearson correlation coefficient between predicted *LLR* and actual *LLR*.

The latency to half-maximum and timescale of sample-averaged *LLR* decoding were estimated for individual ROIs in the same manner as for the alpha-band *LLR* encoding analysis described above (Figure 5a-d), with the exception that timescales were estimated using decoding precision traces starting 0.05 s after the latency of peak decoding precision. This ensured that the peak and transient rapid decay in decoding precision for all ROIs did not contribute to timescale estimates. Parameter differences between ROIs were again assessed via weighted permutation test (see *Model-based analysis of different RO/s during decision-making task* above), with participant-specific weights here determined by the minimum time-averaged decoding precision across the ROIs being compared.

#### Effect of sample position on LLR responses

One concern may be that attention to later samples may have caused the recency effects that we observed in psychophysical kernels. To rule out this possibility, we assessed whether the *LLR* response in a subset of ROIs (V1, V2-V2, V3A/B, IPS0/1, IPS2/3) in the Experiment 1 data varied over sample position via three measures: *LLR* encoding in the alpha-band (8-14 Hz) and gamma frequency-band (40-65 Hz), and *LLR* decoding precision from the evoked response (see below). We extracted each of these measures within time windows following sample onset chosen to include the respective peak encoding/decoding, regressed each of these measures onto sample position, and tested whether the group distribution of regression line slopes was different from zero via permutation test (10,000 permutations). Effects of sample position were mostly absent, weak when present, and not consistent across ROIs or the three measures (data not shown). In sum, there was no systematic difference in the processing of later vs. earlier samples.

#### Impact of residual fluctuations on choice

To interrogate the relevance of residual fluctuations in the ROI-specific neural signals for choice in the data from Experiment 1, we extracted for each ROI the time point-, frequency- and sample-specific standardized residuals *resid_t,f,s_* from the fit of eq. 22 – i.e. a vector that captures across-trial fluctuations in the neural signal that are not explained by decision-relevant computational variables. We then examined whether these residual signal fluctuations predicted variance in behavioral choice over and above the choice-predictive factors identified previously, via the following logistic regression (**Model 6**):

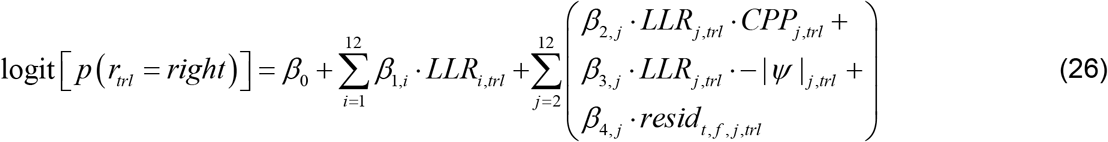

where *β*_4_ captured the effect of interest per sample position *j,* and all other terms were identical to those in eq. 16. We averaged the fitted *β*_4_ across sample positions 2-9 and 10-12 to increase signal-to-noise, under the reasoning that the recency effects in evidence weighting observed in participants’ psychophysical kernels (Figure 2b) would correspond to stronger choice-predictive residual fluctuations toward the end of the trial. For each ROI and sample position subset, we then used cluster-based permutation testing to identify time points at which the fitted weights differed significantly from zero (two-tailed one-sample *t*-test; 10,000 permutations; cluster-forming threshold of *p*<0.05; Figure 6).

Motivated by these results, we performed a more selective version of the above analysis on data from Study 3. We first fit a version of **Model 6** (eq. 26 but excluding the −|*ψ*| modulatory term, which did not contribute to normative belief updating in this generative setting; Supplementary Figure 8a) to source reconstructed estimates of activity in M1, separately for each *H* condition. Averaging the resulting time-frequency effect maps across *H* conditions revealed choice-predictive fluctuations that were more restricted in the frequency domain compared to what we observed for Experiment 1 – primarily to a 12-17 Hz frequency band (Supplementary Figure 8c). We then searched for a corresponding choice signal in visual cortex (two ROIs subsets consisting of V1 and V2-IPS3) by again fitting **Model 6** to signals from these ROIs, but now averaging the fitted coefficients over this frequency band and from 0.2-1.0 s post-sample (informed by the Study 1 results) to yield a single scalar estimate of the choice-predictive signal per subject and *H* condition (Figure 7f). We assessed whether these estimates were significantly different from zero at the group level, and different across *H* conditions, via two-tailed permutation test (10,000 permutations).

#### Power law exponents of intrinsic activity fluctuations

We also assessed regional differences in the contributions of fast vs. slow frequencies to ‘intrinsic’ activity fluctuations in Experiment 1 (Supplementary Figure 7). For each cerebral hemisphere, visual cortical ROI (V1, V2-V4, V3A/B, IPS0/1, IPS2/3), and trial, we computed the power spectrum (from 1-120 Hz) of activity in the 1 s ‘baseline’ interval preceding trial onset using Fast Fourier transform. Power spectra were then averaged over trials and hemispheres. These regionally-specific power spectra of intrinsic activity fluctuations were modeled as a linear superposition of two functional processes^80^: an aperiodic component modeled as a decaying exponential; and a variable number of periodic components (bandlimited peaks, e.g., the ~10 Hz peak). Power *P* at frequency *f* of the aperiodic component was modeled as a power law: log_10_ (*P* (*f*)) = 10^*b*^*f^-χ^*, where *b* was the broadband offset of the spectrum and *χ* was the so-called scaling exponent^81^. The periodic components were modeled as Gaussians with parameters *A, μ* and *σ,* describing the amplitude, peak frequency and bandwidth of the component, respectively. We fit this model to the power spectra from each ROI and participant using the FOOOF toolbox^80^ (default constraints and approach, without so-called ‘knees’ which were absent in the measured spectra, presumably due to the short intervals). The bands 49-51 Hz and 99-101 Hz, which were contaminated by line noise, were excluded from the fits. The model fits are shown in Supplementary Figure 7a.

Our subsequent analyses focused on the power law scaling exponent *χ*. We used this parameter to quantify the relative contributions of fast vs. slow activity fluctuations to the aperiodic component (large exponent equates to greater contribution of slow fluctuations). We assessed whether there was a spatial gradient in the fitted exponents across the visual cortical hierarchy (V1, V2-V4, V3A/B, IPS0/1, IPS2/3) via regular permutation tests (since, unlike for previous analyses of spatial gradients relying on weighted permutation tests, meaningful *χ* estimates were here available for all participants and ROIs). We fitted a line to the hierarchically-ordered, group-average parameters. Repeating this procedure for 10,000 random shuffles of the ROI labels (shuffling at the level of individual participants) yielded a null distribution of slopes, from which we extracted the *p*-value of the slopes across ROIs obtained for unshuffled labels.

#### Limitations of approach

Despite the richness of our results, the current approach also has limitations. In particular, MEG source estimation is limited by signal leakage, due to a combination of the limited spatial resolution of the source reconstruction and volume conduction in the neural tissue. We sought to minimize these effects by restricting our analysis to a selection of ROIs (see above) with relatively low spatial granularity. Furthermore, we focused the interregional comparisons on *differences* (rather than *similarities*) – in other words, the heterogeneity of computational properties along the visuo-motor pathway (Figure 5). For example, signal leakage cannot explain the observed similarity between primary visual cortex and anterior decision-encoding regions, and their respective dissimilarity with the anatomically-intermediate clusters IPS0/1 and IPS2/3. Because of signal leakage, the interregional heterogeneity we report here constitutes a lower bound for the true heterogeneity of neural activity.

A second limitation of MEG is low sensitivity for subcortical sources. Invasive recording techniques will be required to illuminate the contributions of the striatum, superior colliculus, thalamus, and cerebellum to the decision computations that we characterize here.

### Pupillometry data analysis

#### Preprocessing

Eye blinks and other noise transients were removed from pupillometric time series from Experiment 1 using a custom linear interpolation algorithm in which artifactual epochs were identified via both the standard Eyelink blink detection algorithm and thresholding of the first derivative of the per-block z-scored pupil time-series (threshold = ±3.5zs^-1^). The pupil timeseries for each block was then band-pass filtered between 0.06-6Hz (Butterworth), re-sampled to 50Hz and z-scored across time. We then computed the first derivative of the resulting timeseries, which was the focus of all subsequent analyses. As such, our preprocessing of the pupil signal promoted sensitivity to fast, evoked changes in pupil size and removed slower fluctuations within and across task blocks.

#### Sensitivity to change-point occurrence

We first visualized the sensitivity of the pupil derivative signal to change-point occurrence in the following manner. The signal was segmented around −0.5–5.8 s relative to trial onset for all full-length (12-sample) trials, thus encompassing the duration of decision formation on these trials. We then discarded all trials on which >1 change-point occurred, averaged over subsets of trials on which single change-points occurred at sample positions 2-4, 5-6, 7-8, 9-10 or 11-12, and from each of the resulting traces subtracted the average signal from trials on which no change-point occurred. This procedure created 5 time-series of the pupil derivative around change-points that occurred at different points in time over the trial, relative to the case where no change-point occurred (Figure 8b). On the same plot we overlaid relative *CPP* for the same subsets of trials for comparison, as calculated from the best-fitting normative model fits.

#### Sensitivity to sample-wise computational variables and relevance for choice

Next, we assessed the sensitivity of the pupil signal to decision-relevant computational variables derived for each presented sample from the normative model fits. Pupil size is a signal with demonstrated sensitivity to various forms of surprise and uncertainty in other task contexts^45^, but not to ‘signed’ choice-selective variables (such as *LLR* and *ψ* in the present case). Thus, unlike the lateralized MEG signals above for which *CPP* and –*|ψ| modulate* the influence of new information on signal change, for the pupil we interrogated direct relationships with these variables. We did so by segmenting the pupil derivative from 0 to 1s after the onset of each individual sample *s* on full-length trials, and fitting the following linear regression model (**Model 7**; Figures 8c):

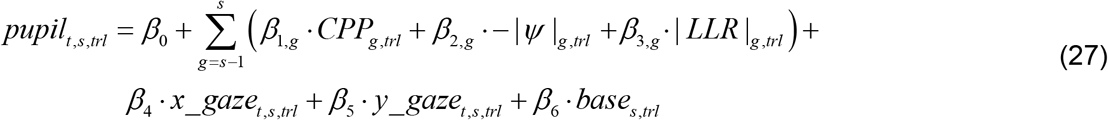

where *t* indicated time point relative to sample onset, *x_gaze* and *y_gaze* were the instantaneous horizontal and vertical gaze positions included to absorb a possible artifactual effect of gaze position on measured pupil size, and *base* was the raw ‘baseline’ pupil diameter from −0.05-0.05 s around sample onset included to control for the effect that absolute pupil size at stimulus onset exerts on the magnitude of the evoked response. |*LLR*| is included to capture a possible relationship between pupil and a low-level form of surprise (since |*LLR*| exhibits a monotonic negative relationship in our task with the unconditional probability of observing a given stimulus over the entire experiment; Supplementary Figure 10). We further include previous sample *CPP*, – *|ψ|* and |*LLR*| in the model because the pupil response to an eliciting impulse is relatively slow, and thus correlations with model-based variables from the previous sample can cause spurious correlations with those from the current sample within the time window considered here (in particular, a positive correlation that was present between *CPP_s-1_* and – *|ψ|_s_*). As above for the MEG signals, significant effects for the terms of interest were assessed via two-tailed cluster-based permutation testing after averaging the associated t-scores over sample positions. We also delineated the sensitivity of the evoked pupil response to other candidate surprise measures, as described in Supplementary Figure 10.

We interrogated the relevance of variability in the sample-wise pupil response for choice via a similar approach to that described for the MEG signals above (**Model 8**; Figure 8d). Specifically, we fit versions of eq. 26 in which the final term (*β*_4_) was *LLR_s,trl_ resid_t,s,trl_*, where *resid* here refers to the standardized residuals from fits of eq. 27. Here this term takes the form of a modulation of *LLR* weighting and not a direct effect on choice as in eq. 26, which accounts for the fact that the pupil response is a modulatory signal and, unlike the lateralized MEG signals, is not sensitive to the relative evidence/belief for different choice alternatives.

#### Relationship with neural signals

Lastly, we investigated the relationship between the pupil response to each sample in Experiment 1, and the previously identified neural signals that encode evidence strength and belief. We extracted the per-sample standardized residuals from eq. 27 at *t* = 0.57s post-sample (time of peak *CPP* encoding in the pupil; Figure 8c), and assessed whether this metric modulated the neural encoding of the associated evidence sample via the following linear model (**Model 9**):

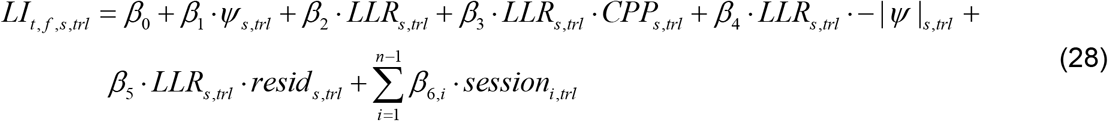

where *LI_t,f,s,trl_* was the power lateralization for a given ROI. This model was an extension of eq. 22 where the term of central interest (*β*_5_) captures the extent to which the pupil response to sample *s* enhances or suppresses the encoding of that sample in the lateralized signal, over and above the variance in the signal captured by variables from the model fits that reflect evidence strength and belief updating.

In addition to the selective modulatory pupil effect captured by **Model 9**, we also assessed whether the pupil response to each sample exerted a more non-selective effect on timefrequency power. Here we averaged the source-reconstructed time-frequency signal across cerebral hemispheres, and for each sample position regressed this signal onto the pupil response again measured at *t* = 0.57s post-sample (Supplementary Figure 9).

## Acknowledgements

We thank Klaus Wimmer for discussion and advice on cortical circuit model and Florent Meyniel for detailed comments on the manuscript.

This work was funded by the Deutsche Forschungsgemeinschaft (DFG, German Research Foundation) – DO 1240/3-1, DO 1240/4-1, and SFB 936 – Projekt-Nr. A7.

## Author contributions

PRM: Conceptualization, Methodology, Investigation, Software, Formal analysis, Visualization, Writing – original draft, Writing – review and editing; NW: Conceptualization, Methodology, Software, Writing – review and editing; DCHB: Investigation, Formal analysis; GPO: Software, Formal analysis, Writing – review and editing; THD: Conceptualization, Methodology, Resources, Writing—original draft, Writing—review and editing, Supervision.

## Data availability

Raw behavioral data and preprocessed MEG/pupil data will be made available on Figshare upon acceptance.

## Code availability

All analysis code to reproduce the reported results and figures from the shared data will be made available on GitHub (https://github.com/DonnerLab) upon acceptance.

## Supplementary Figures

**Supplementary Figure 1.**
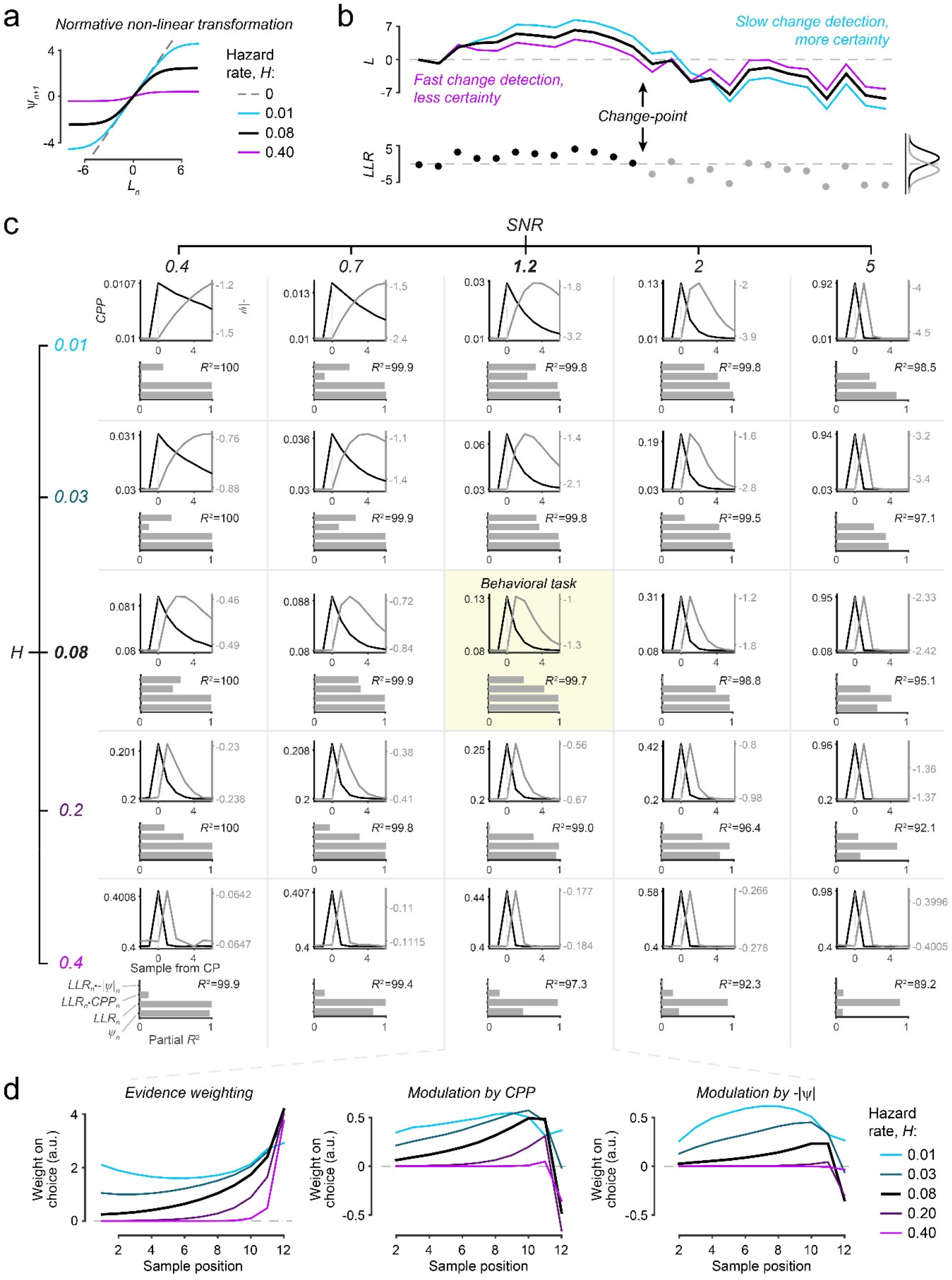
Sensitivity of normative evidence accumulation to change-point probability and uncertainty across a range of generative task statistics. (**a**) Non-linearity in normative model for different hazard rates (*H*): Posterior belief after accumulating the most recent sample (*L_n_*) is converted into prior belief for next sample (*ψ_n+1_*, both variables expressed as log-odds for each alternative) through a non-linear transform, which saturates (slope≈0) for strong *L_n_* and entails more moderate information loss (0<<slope<1) for weak *L_n_.* In a static environment (*H*=0), the model combines new evidence (log-likelihood ratio for sample *n*, *LLRn)* with prior *(ψ_n_*) into updated belief without information loss (i.e. *L_n_=ψ_n_+LLR_n_; ψ_n+1_=L_n_*). In an unpredictable environment (*H*=0.5), the new belief is determined solely by the most recent evidence sample (i.e. *L_n_=LLR_n_; ψ_n+1_*=0) so that no evidence accumulation takes place. (**b**) Example trajectories of the model decision variable (*L*) as function of the same evidence stream, but for different levels of *H*. (**c**) Upper panels of each grid segment: Change point-triggered dynamics of change-point probability (*CPP*) and uncertainty (-|*ψ*|) derived from normative model as function of *H* (grid rows) and evidence signal-to-noise ratio (SNR, difference in generative distribution means over their standard deviation; grid columns). Lower panels of each grid point: Contribution of computational variables (including *CPP* and – |*ψ*|) to normative belief updating, expressed as coefficients of partial determination for terms of a linear model predicting the updated prior belief for a forthcoming sample. Yellow background (center of grid): generative statistics of the task used in Experiment 1; reproduced from main Figure 1e for comparison. Data for each parameter combination were derived from a simulated sequence of 10 million observations. (**d**) Psychophysical kernels of the normative model for the task used in Experiment 1 (12 samples per trial, generative *SNR≈1*.2) but for each level of *H* from panel c. Kernels reflect the time-resolved weight of evidence on final choice (left) and the modulation of this evidence weighting by sample-wise changepoint probability (*CPP*, middle) and uncertainty (-|*ψ*|, right). Kernels were produced by adding a moderate amount of decision noise (*v*=0.7, see *Methods)* to the ideal observer (i.e. the normative model with perfect knowledge of generative statistics); without noise, coefficients at high *H* (where choices are based almost entirely on the final evidence sample) are not identifiable.

**Supplementary Figure 2.**
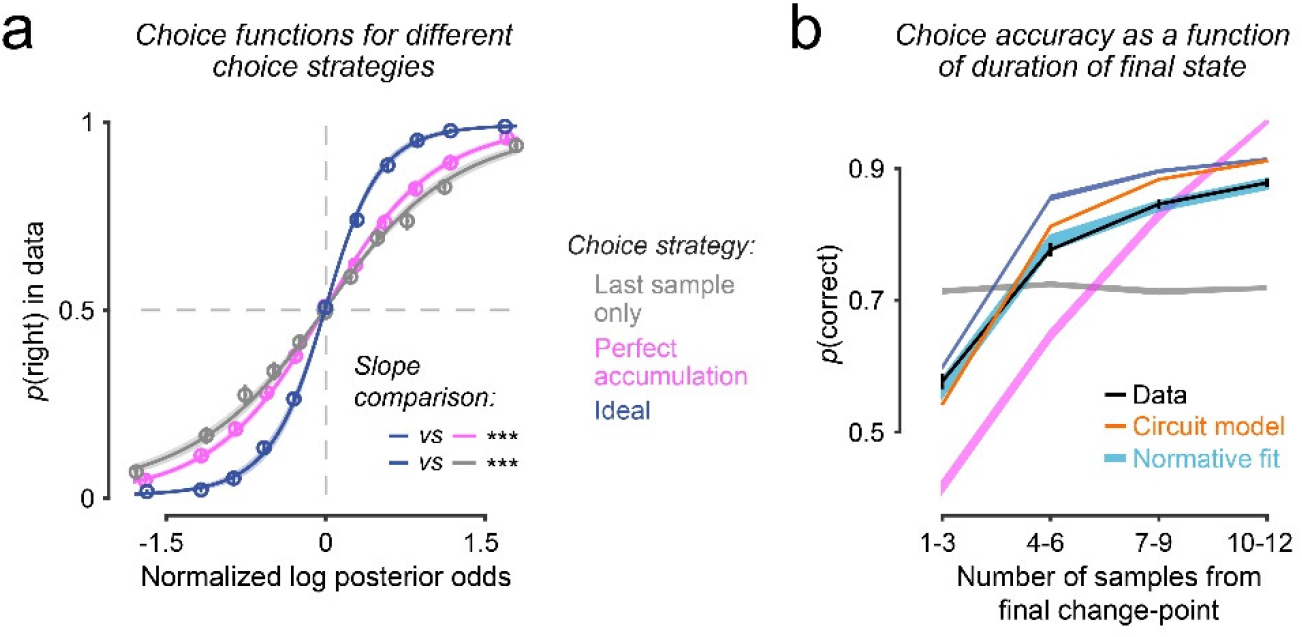
Consistency of human choices with those of idealized choice strategies, and dependence of choice accuracy on final state duration. (**a**) Human choices as function of log-posterior odds (z-scored) derived from each of three alternative strategies: basing choices only on final evidence sample (gray); perfect evidence accumulation (magenta); and an ideal observer with perfect knowledge of the generative task statistics and employing the normative accumulation process for the task (blue). Points and error bars show observed data ± s.e.m.; lines and shaded areas show mean ± s.e.m. of fits of sigmoid functions to the data. Slopes of fitted sigmoids were steepest for ideal observer (ideal vs. perfect accumulation: t_16_=5.9, p<0.0001; ideal vs. last-sample: t_16_=9.6, p<10^-7^; two-tailed paired t-tests), indicating that human choices were most consistent with those of the ideal observer. (**b**) Choice accuracy as a function of duration of the final environmental state for the human participants (black), idealized strategies from panel a, fits of the normative model (cyan), and the circuit model (orange). Participants’ choice accuracy increased with the number of samples presented after the final state change on each trial, consistent with temporal accumulation. Error bars and shaded regions indicate s.e.m.

**Supplementary Figure 3.**
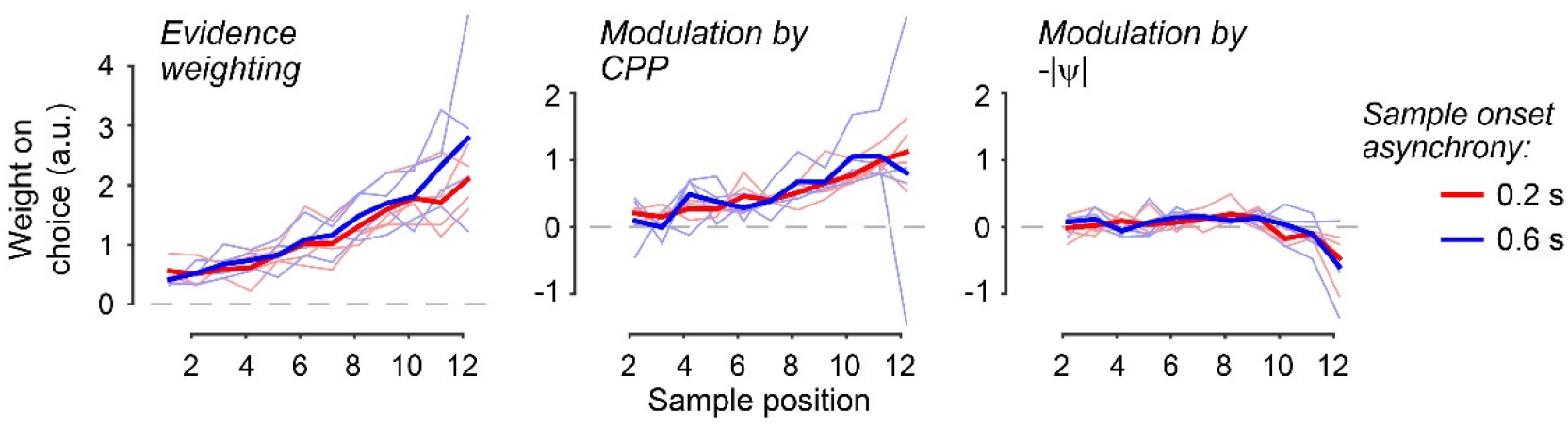
Effect of sample onset asynchrony (SOA) on diagnostic signatures of adaptive evidence accumulation. Psychophysical kernels reflecting the time-resolved weight of evidence on final choice (left), and the modulation of this evidence weighting by sample-wise change-point probability (*CPP*, middle) and uncertainty (-|*ψ*|, right), separately for the two conditions of Experiment 2 (*N*=4) in which participants performed the decision-making task at fast (0.2 s) and slow (0.6 s) sample onset asynchronies. Thin unsaturated lines are individual participants, thick saturated lines are means across participants. Cluster-based permutation tests (10,000 permutations) of SOA effects on each kernel type revealed no significant effects.

**Supplementary Figure 4.**
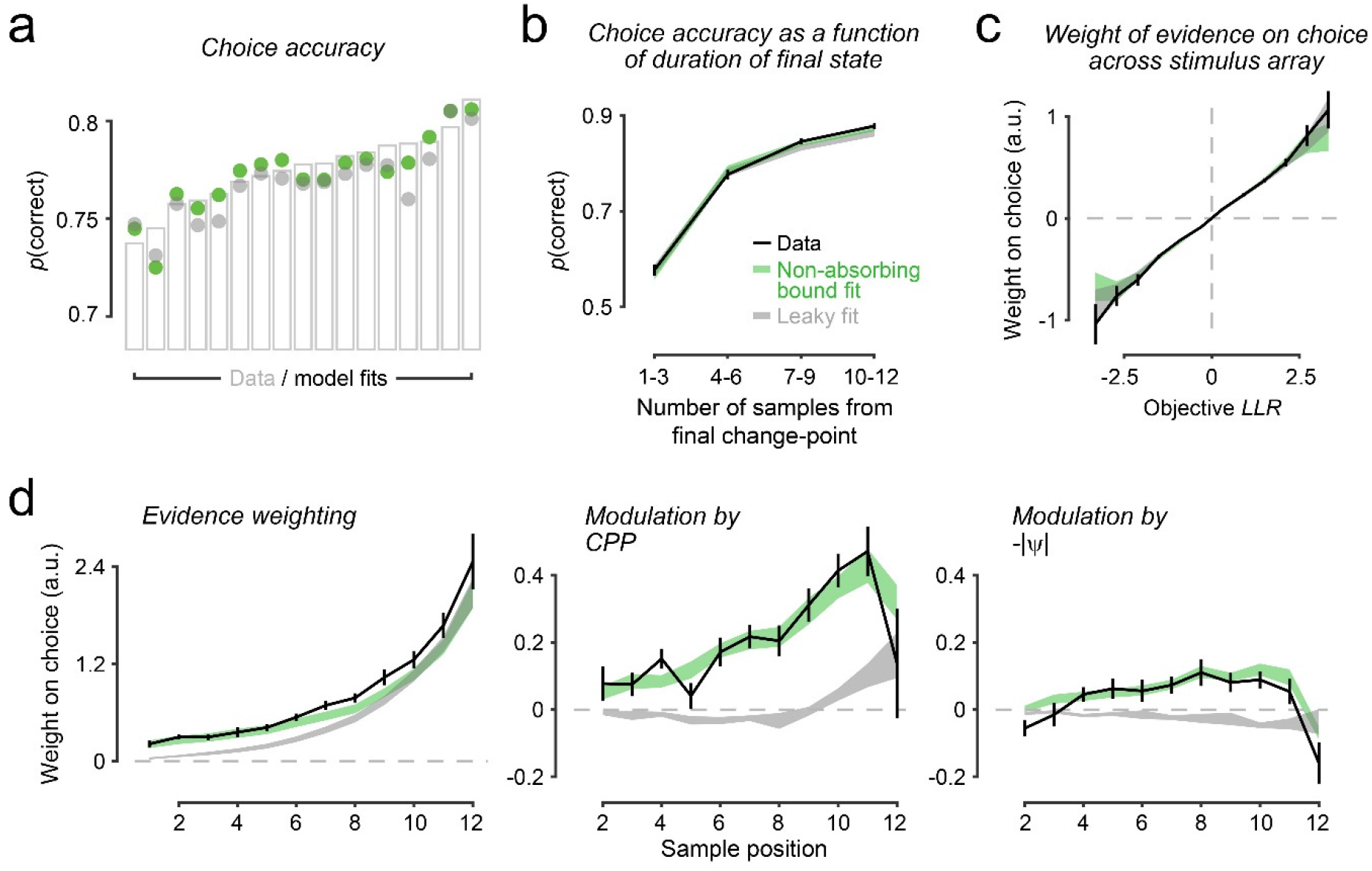
Fit to human behavior of alternative evidence accumulation schemes. Alternative accumulation schemes are leaky accumulation (gray), where the evolving decision variable is linearly discounted by a fixed leak term after every updating step; and perfect accumulation toward non-absorbing bounds (green), which imposes upper and lower limits on the evolving decision variable without terminating the decision process, thus enabling changes of mind even after the bound has been reached. Both schemes combined can approximate the normative accumulation process across generative settings (ref. 16). (**a**) Choice accuracies for human participants (grey bars) and both model fits. (**b**) Choice accuracy as a function of duration of the final environmental state, for the participants and the models. (**c**) Regression coefficients reflecting the subjective weight of evidence associated with binned stimulus locations, for the participants and the models. (**d**) Psychophysical kernels for the data and the model reflecting the time-resolved weight of evidence on final choice (left), and the modulation of this evidence weighting by sample-wise change-point probability (*CPP*, middle) and uncertainty (-|*ψ*|, right). In panels b-d, error bars and shaded regions indicate s.e.m. Fits of the model versions used to generate behavior in all panels included free parameters for both a non-linear stimulus-to-LLR mapping function and a gain factor on inconsistent samples (see Supplementary Figure 5). Even with these additional degrees of freedom, the leaky accumulator model failed to reproduce the *CPP* and – |*ψ*| modulations characteristic of human behavior. By contrast, like the normative model, perfect accumulation to non-absorbing bounds captured all qualitative features identified in the behavioral data. This is in line with the insight that this form of accumulation closely approximates the normative model in settings like ours, where strong belief states are formed often (i.e. low noise and/or low H; ref. 16). This form of accumulation also captures a key feature of the dynamics of the circuit model that we interrogate in the main text; i.e. a saturating decision variable in response to consecutive consistent samples of evidence.

**Supplementary Figure 5.**
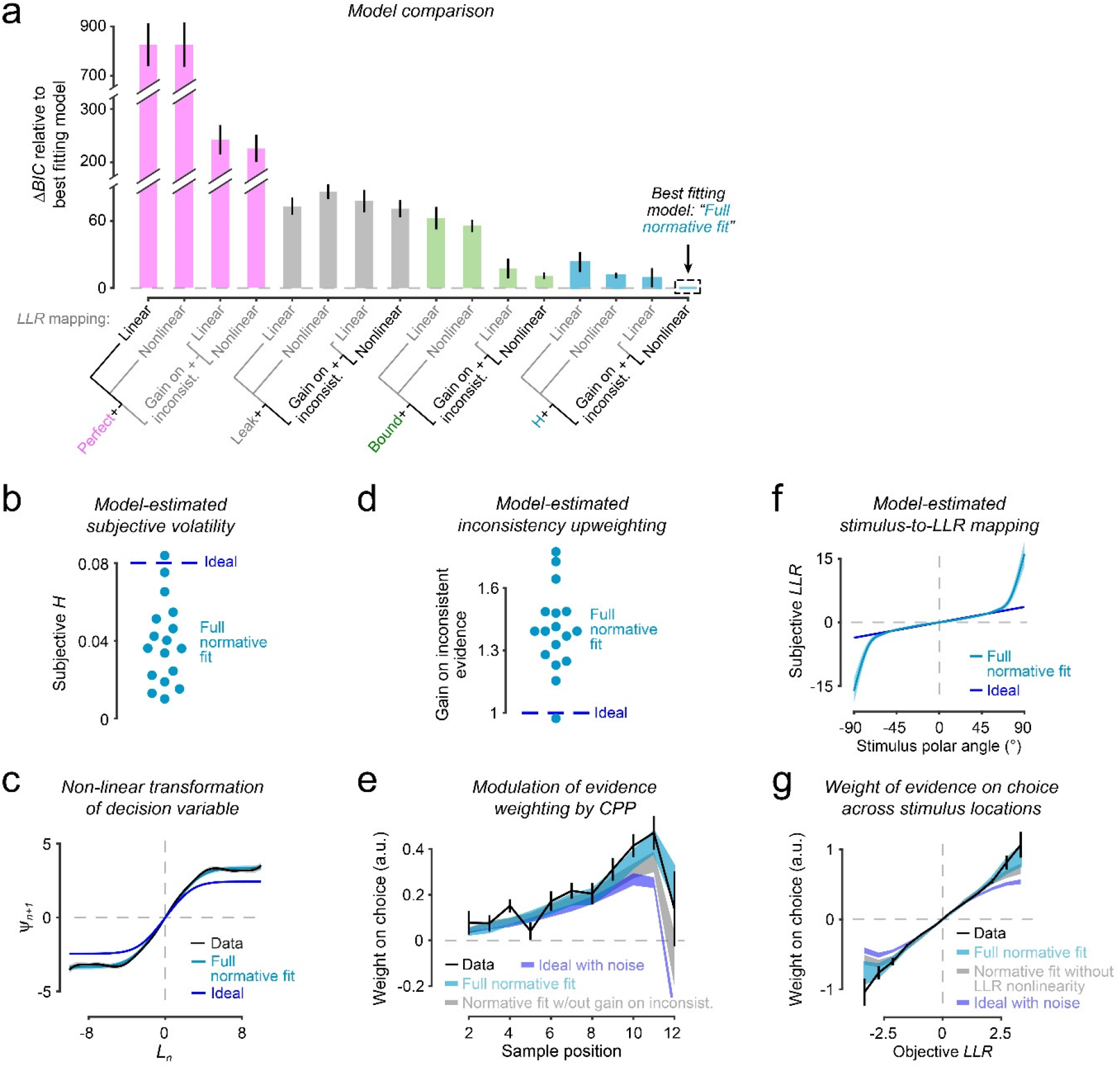
Model comparison and qualitative signatures indicate approximation of normative belief updating by measured human behavior. (**a**) Bayes Information Criteria (BIC) for all models fit to the human choice data, relative to best-fitting model. Bars show mean ± s.e.m. Model constraints are specified by the tick labels. Colors denote unbounded perfect accumulator (magenta), leaky accumulator (gray), perfect accumulator with nonabsorbing bounds (green), and the normative accumulation process with a subjective hazard rate (cyan). Labels refer to the following: ‘Linear’ (‘Nonlinear’)’=linear (non-linear) scaling of stimulus-to-LLR mapping function; ‘Gain on inconsist.’=multiplicative gain term applied to inconsistent samples; ‘Leak’=accumulation leak; ‘Bound’= height of non-absorbing bounds; ‘H’=subjective hazard rate. ‘Perfect’ is a special case of the leaky accumulator where leak=0. All models included a noise term applied to the final log-posterior odds per trial. See *Methods* for additional model details. Labels in black text highlight models plotted in main Figure 2 and Supplementary Figure 4. Best fitting model (‘full normative fit’ in remaining panels) employs normative accumulation with subjectivity in hazard rate, a gain term on inconsistent samples, and non-linear stimulus-to-LLR mapping. (**b**) Subjective hazard rates from full normative fits (cyan), with true hazard rate in blue. Participants underestimated the volatility in the environment (t_16_=-7.7, *p*<10^-6^, two-tailed one-sample t-test of subjective *H* against true *H*). (**c**) Non-linearity in evidence accumulation estimated directly from choice data (black) and from full normative model fits (cyan). Non-linearity of the ideal observer shown in blue. Shaded areas, s.e.m. (**d**) Multiplicative gain factors applied to evidence samples that were inconsistent with the existing belief state, estimated from full normative fits. Gain factor>1 reflects relative up-weighting of inconsistent samples beyond that prescribed by normative model; gain factor<1 reflects relative down-weighting of inconsistent samples. The ideal observer employing the normative accumulation process uses gain factor=1 (dashed blue line). Participants assigned higher weight to inconsistent samples than the ideal observer (t_16_=8.2, *p*<10^-6^, two-tailed one-sample *t*-test of fitted weights against 1). (**e**) Regression coefficients reflecting modulation of evidence weighting by change-point probability (*CPP*). Shown are coefficients for the human participants (black line), full normative fits (cyan), normative model fits without gain factor applied to inconsistent evidence (grey), and the ideal observer with matched noise (blue). Error bars and shaded regions, s.e.m. (**f**) Mapping of stimulus location (polar angle; x-axis) onto evidence strength *(LLR;* y-axis) across the full range of stimulus locations. Blue line reflects the true mapping given the task generative statistics, used by the ideal observer. Cyan line and shaded areas show mean ± s.e.m of subjective mappings used by the human participants, estimated as an interpolated non-parametric function (*Methods*). (**g**) Regression coefficients reflecting subjective weight of evidence associated with binned stimulus locations, for the human participants (black line), full normative model fits including the non-linear *LLR* mapping shown in panel f (cyan), normative model fits allowing only a linear *LLR* mapping (grey), and the ideal observer with matched noise (blue). Error bars and shaded regions, s.e.m.

**Supplementary Figure 6.**
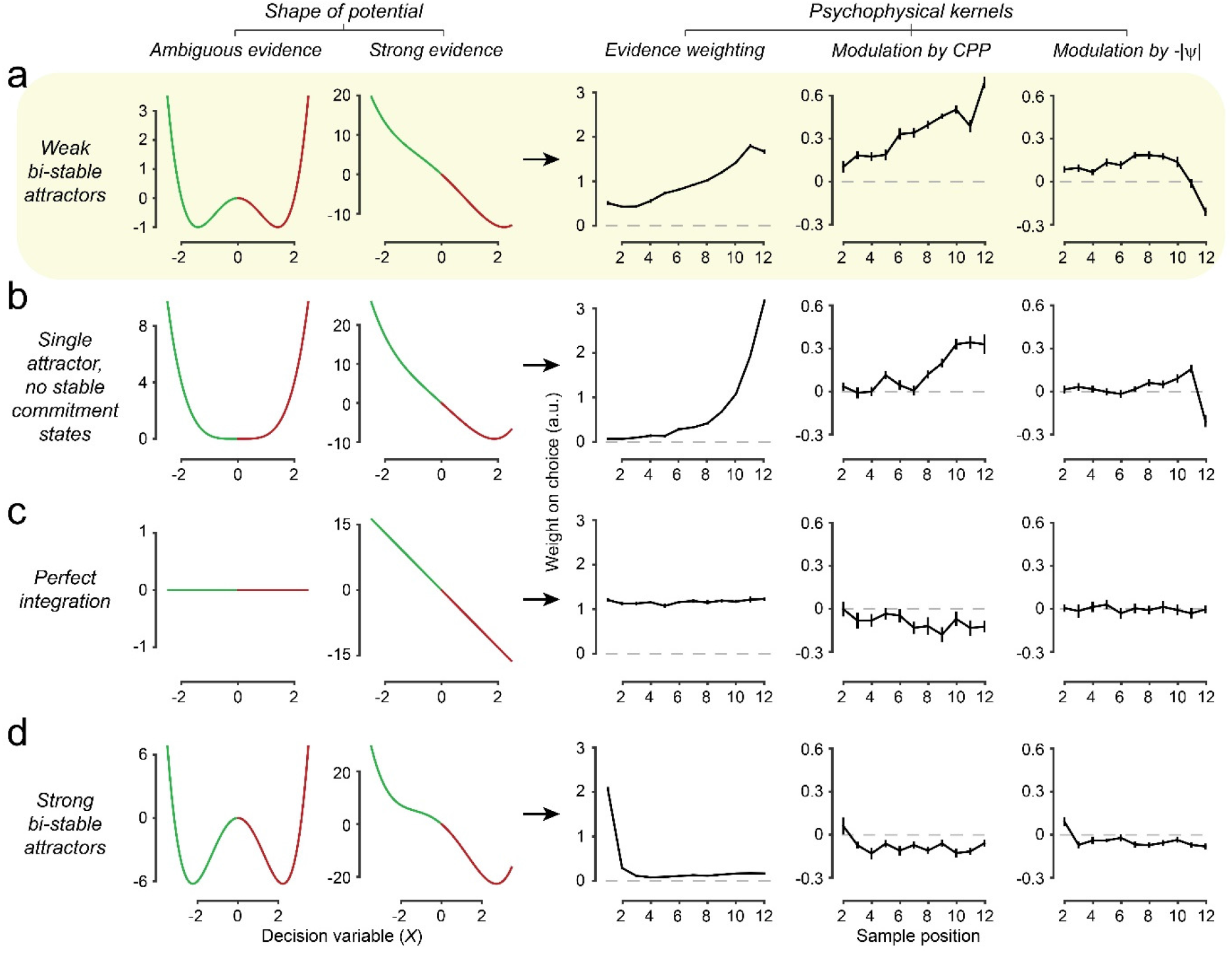
Assessment of boundary conditions at which circuit model approximates normative evidence accumulation. A reduction of the spiking-neuron circuit model (*Methods*) was used to explore the impact of different dynamical regimes on behavioral signatures of evidence weighting. Left two columns: shape of model’s energy landscape (‘potentials’) in response to ambiguous evidence (*LLR*=0, left), or in response to strong evidence (*LLR*=mean+1 s.d. of *LLRs* across experiment=1.35, second from left). Middle-to-right columns: psychophysical kernels and modulations of evidence weighting by *CPP* and – |*ψ*|. (**a**) Double-well potential with small barrier between wells, corresponding to weak bi-stable attractor dynamics. This model variant featured a saturating decision variable in the face of multiple consistent evidence samples (akin to non-absorbing bounds), wells that maintained commitment states but were sufficiently shallow for changes-of-mind to occur in response to strongly inconsistent evidence, and sensitivity to new input when at the ‘saddle point’ between wells (i.e. during periods of uncertainty). As a consequence, it produced all key qualitative features of normative evidence accumulation (recency, strong modulation by CPP; weak modulation by uncertainty). **b**) Single-well potential with no barrier. This model variant also featured a saturating decision variable, but no stable states of commitment. Thus, it exhibited stronger recency in evidence weighting than the one in panel a. (**c**) Flat potential with no wells, corresponding to perfect evidence accumulation without bounds. This model lacked both a saturating decision variable and stable states. As a consequence, it weighed all evidence samples equally (flat psychophysical kernel) and did not produce clear modulations by CPP or uncertainty. (**d**) Double-well potential with high barrier between wells, corresponding to strong bi-stable attractor dynamics. In this model variant, the wells were too deep for changes-of-mind to occur for all but the most extreme inconsistent evidence. Thus, it produced extreme primacy in evidence weighting. The evidence weighting signatures of the regimes in panels c, d were qualitatively inconsistent with the signatures of participants’ behavior (compare Figure 2b). All three signatures were qualitatively reproduced by the regimes in both a and b, with the closest approximation to human behavior provided by the weak bi-stable attractor regime in panel a (highlighted in yellow).

**Supplementary Figure 7.**
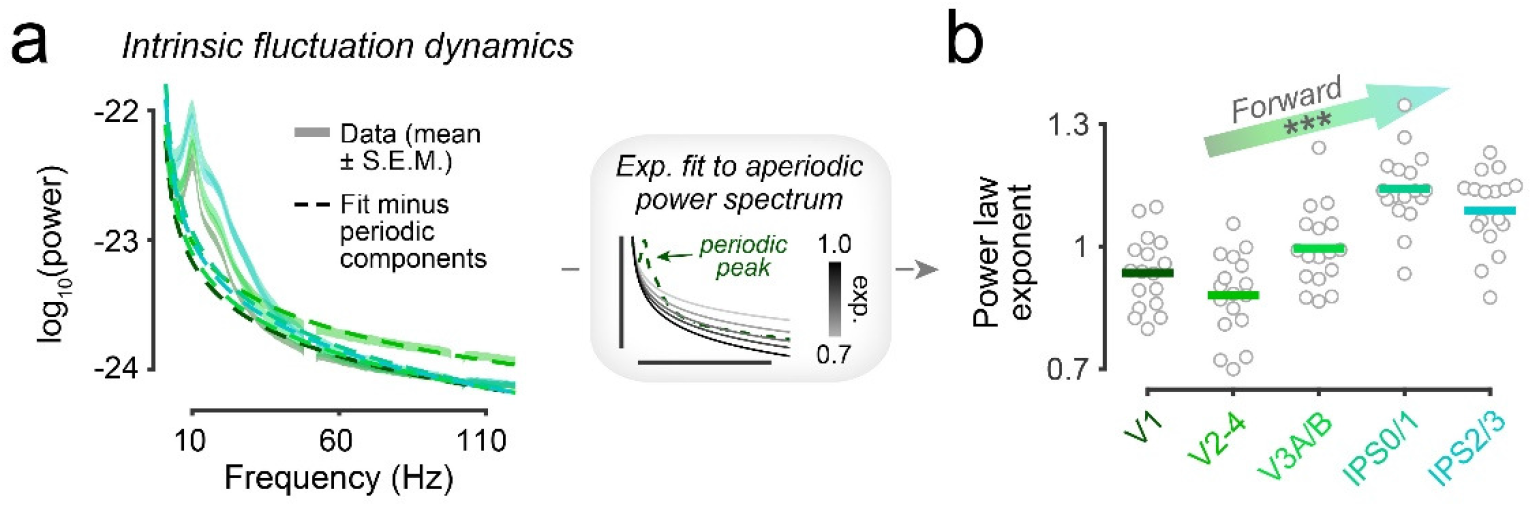
Intrinsic fluctuation dynamics across visual cortical hierarchy. (**a**) Power spectra and associated model fits of intrinsic fluctuations across cortex. For each hemisphere, ROI, and trial, we computed the power spectrum from a 1-second ‘baseline’ period preceding trial onset and averaged these spectra across trials and hemispheres. Shaded areas show mean +/− s.e.m of observed power spectra. We modeled the power spectra as a linear superposition of an aperiodic component (power law scaling) and a variable number of periodic components (band-limited peaks, Methods). The fitted aperiodic components only are overlaid as dashed lines. (**b**) Power law scaling exponents of the aperiodic components estimated from fits in g, providing a measure of the relative contributions of fast vs. slow fluctuations to the measured power spectrum. Lines, group mean; dots, individual participants. ‘Forward’ indicates direction of hierarchy inferred from fitted exponents across the dorsal set of visual field map ROIs (V1, V2-V4, V3A/B, IPS0/1, IPS2/3); \*\*\**p*<0.001, two-tailed permutation test.

**Supplementary Figure 8.**
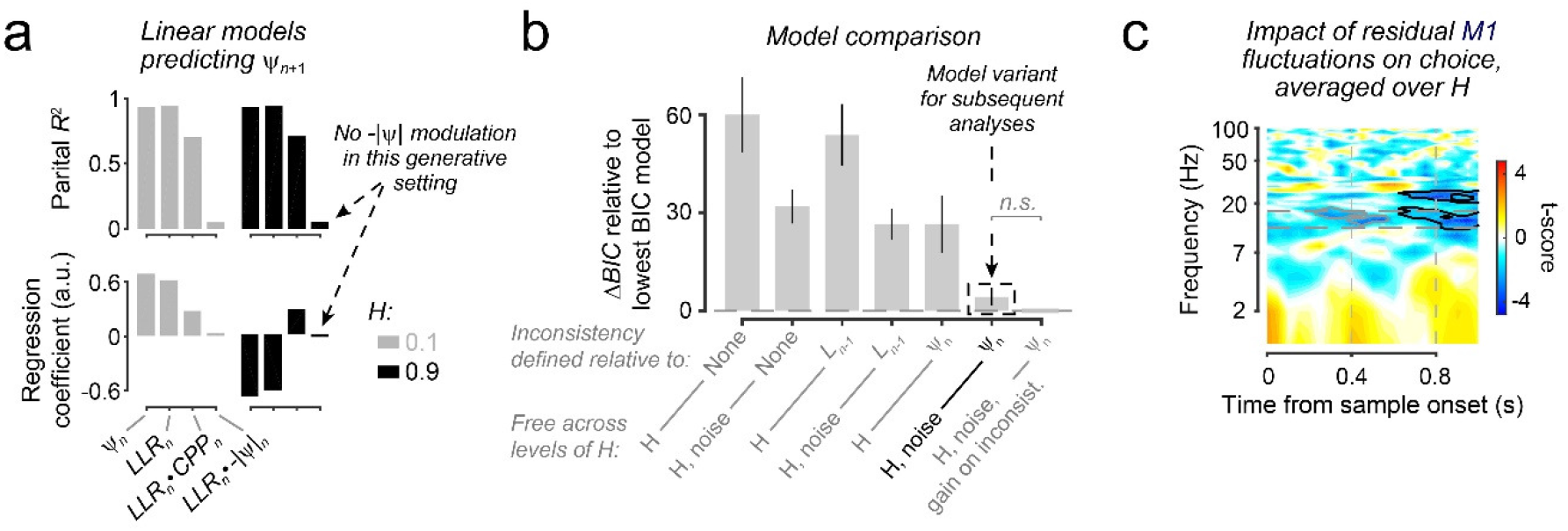
Normative accumulation, model fits and motor choice signal for Experiment 3. (**a**) Contribution of computational variables (including *CPP* and – |*ψ*|) to normative belief updating in generative contexts from Experiment 3, expressed as coefficients of partial determination (upper) and regression coefficients (lower) for terms of a linear model predicting the updated prior belief for a forthcoming sample (*ψ_n+1_*). Note that the strength of the contributions of each term is the same across the two levels of *H*, which is expected when the chosen levels of *H* are symmetric around 0.5 as here. However, the sign of the contributions of prior (*ψ_n_*) and new evidence (*LLR_n_*) are reversed under high *H*, accounting for the flipping of belief sign imposed by the normative non-linearity (Figure 7b, main text). Note also the minimal contribution of –|*ψ*| in this generative setting (*H*={0.1, 0.9}; *SNR*=2), which prompted us to not consider –|*ψ*| modulatory effects in analyses of the data from human participants. (**b**) Bayes Information Criteria (BIC) for all models fit to the human choice data from Experiment 3, relative to model with the lowest BIC score averaged over participants. Bars show mean ± s.e.m. Model constraints are specified by the tick labels. All models included at least one subjective hazard rate, linear scaling parameter applied to the stimulus-to-*LLR* function, and a decision noise term applied to the final log-posterior odds per trial. Models could also include a gain parameter applied to evidence that is inconsistent with either the posterior after the last sample (*L_n-1_*) or the prior for the next sample (*ψ_n_*). We fit models with a variety of different parameter constraints over *H* conditions, as specified via the tick labels. See *Methods* for additional model details. Label in black text highlights model used for remaining analyses. This model included a gain on inconsistent evidence relative to *ψ_n_*, and allowed *H* and noise to vary across *H* conditions. This model produced BIC scores that were marginally, but not significantly, higher than a less parsimonious model in which the gain parameter was also free to vary across *H* conditions. (**c**) Weight of residual fluctuations in the MEG lateralization signal from M1 on final choice, over and above the weight on choice exerted by key model variables. Effects shown are averaged over final 3 evidence samples (positions 8-10) in the trial, and averaged over *H* conditions. Colors reflect group-level t-scores. Black contours, *p*<0.05, two-tailed cluster-based permutation test; gray contours, largest sub-threshold clusters (*p*<0.05, two-tailed t-test). The highlighted frequency band encompassing the majority of the choice-predictive effect in M1 (12-17 Hz) was used to identify choice signals in early visual cortex (main text Figure 7f).

**Supplementary Figure 9.**
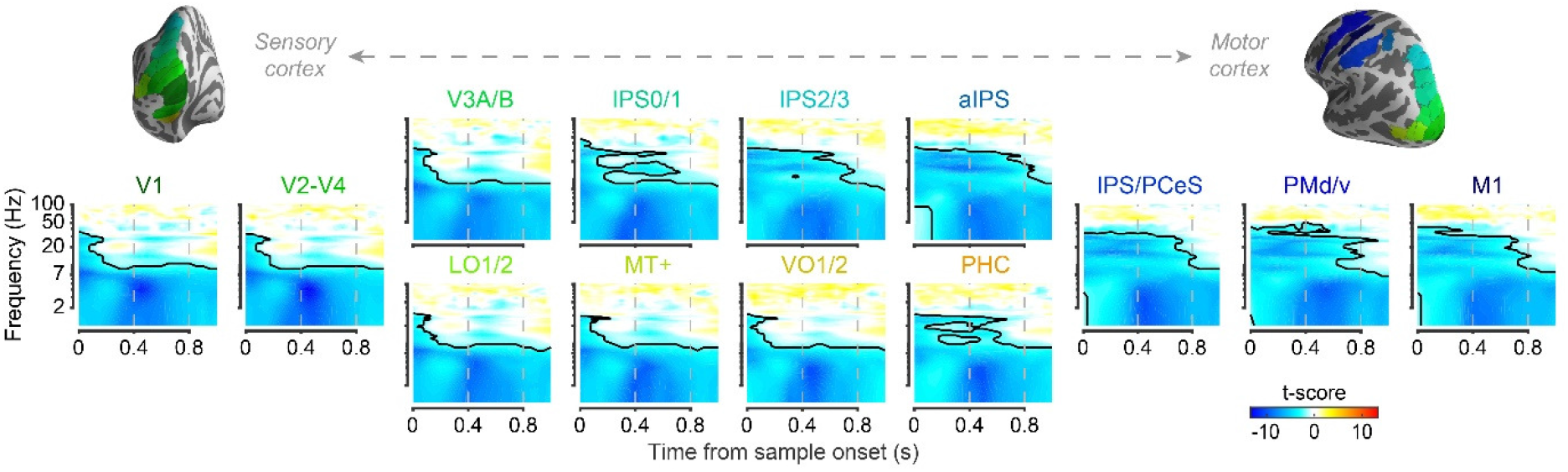
Effect of sample-wise pupil response (derivative) on source-localized, hemisphere-averaged MEG signal. Colors reflect group-level t-scores derived from subject-specific regression coefficients. Contours, *p*<0.05 (two-tailed cluster-based permutation test). Large pupil responses were associated with a strong decrease in low-frequency (<8 Hz) power across all ROIs, a decrease in alpha- and beta-band (8-35 Hz) power predominantly in intraparietal and (pre)motor regions, and a weak increase in high-frequency (60-100 Hz) power in a restricted set of ROIs (*p*<0.05, uncorrected two-tailed permutation tests on regression coefficients averaged over time-points in VO1/2, PHC’ IPS2/3 and aIPS).

**Supplementary Figure 10.**
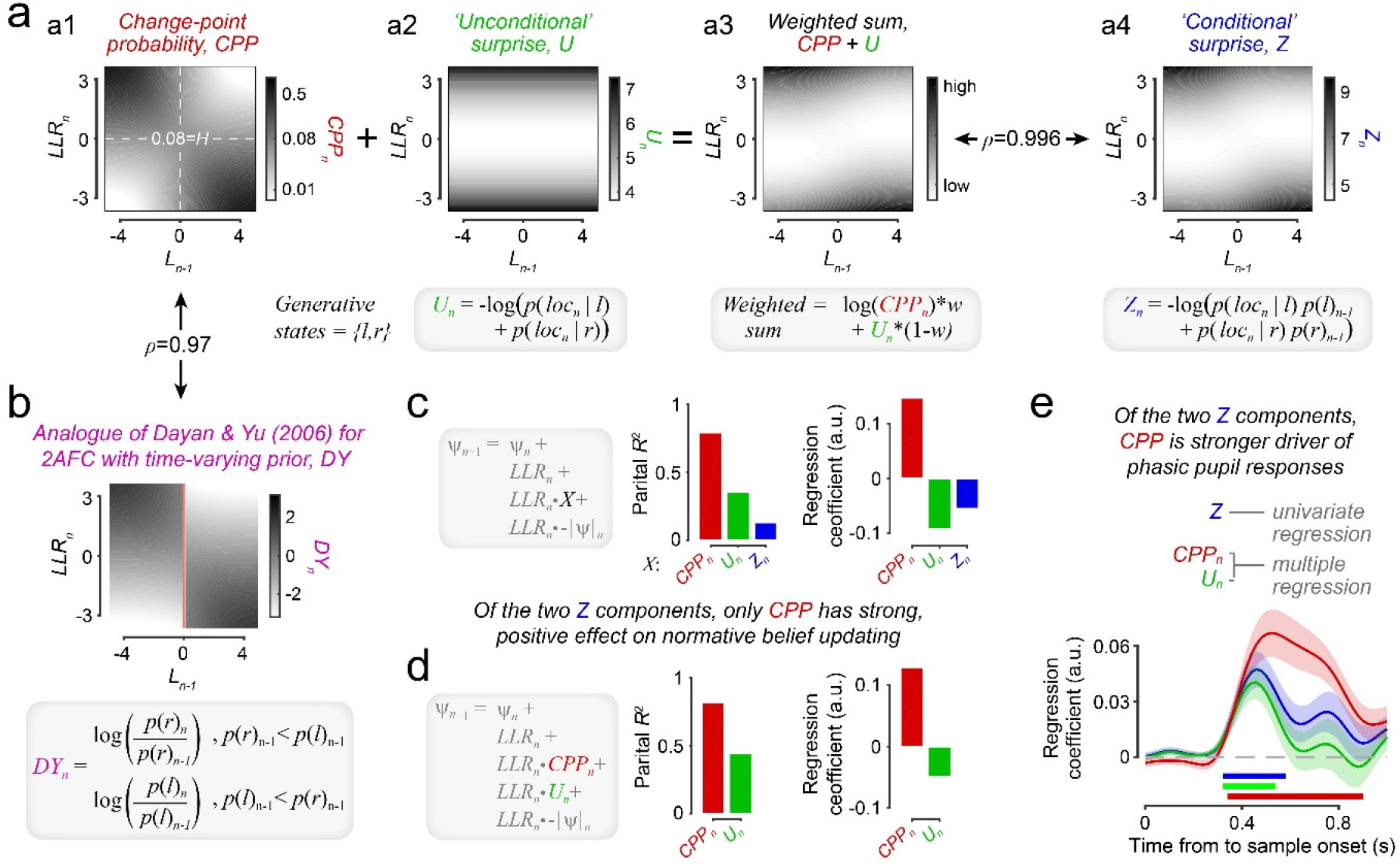
Change-point probability modulates normative belief updating and phasic arousal responses more strongly than alternative metrics of surprise. (**a**) Alternative surprise metrics as a function of posterior belief after previous sample (*L_n-1_*) and evidence provided by the current sample (*LLR_n_*). **(a1)** *CPP* (see main text). **(a2)** ‘Unconditional’ surprise, *U*, defined as Shannon surprise (negative log probability) associated with new sample location (‘*Ioc_n_*’), given only knowledge of the generative distributions associated with each environmental state *S*={*l,r*}. In our task, *U* varied monotonically with absolute evidence strength *(|LLR|)*. **(a4)** ‘Conditional’ surprise *Z:* Shannon surprise associated with new sample, conditioned on *both* knowledge of the generative distributions *and* one’s current belief about the environmental state. Unlike *CPP*, which we use solely to decompose normative evidence accumulation and relate it to neurophysiological signals, in some models this form of surprise serves as the objective function to be minimized by the inference algorithm (e.g. ref. 55) **(a3)** In our task setting, *Z* was closely approximated (Spearman’s *ρ*=0.996) by a linear combination of *log*(*CPP*) and *U* (weights: *w* and 1-*w*, respectively; determined by Nelder-Mead simplex optimization as *w*=0.274). (**b**) Characterization of another form of surprise derived from a model of phasic locus coeruleus activity by ref. 43, hence denoted as ‘Dayan & Yu surprise’ and abbreviated to *DY.* For the oddball target detection task modelled in ref. 43, *DY* was defined as the ratio between the fixed prior probability of an unlikely environmental state and its posterior probability given a new stimulus. In our 2AFC task, where the prior varies during decision formation, we defined *DY* as indicated below the heatmap. We found that this *DY* measure was highly similar to *CPP* (*ρ*=0.97; compare panels b and a1). (**c**) Modulations of normative belief updating by *CPP, U* and *Z*. For each surprise metric, we fit a linear model to updated prior belief (same form as **Model 1;** inset; *X,* surprise metric). Left: coefficients of partial determination; right: contributions expressed as regression coefficients. *CPP* exerted the strongest and only *positive* modulatory effect on evidence weighting, consistent with the observed effects on behavior, cortical dynamics, and pupil (see main text). (**d**) As panel c, but for expanded linear model that included modulations of evidence weighting by *CPP* and *U*’ which combine linearly into *Z*. Only *CPP* yielded a robust, positive effect on evidence weighting. (**e**) Encoding of surprise metrics in pupil response (temporal derivative) to evidence samples. Encoding time course for *Z* was computed by a regression model that included X- and Y-gaze positions as nuisance variables. Encoding time courses for *CPP* and *U* were derived from an expanded regression model that included both terms. *CPP* had the strongest effect of the three considered surprise metrics, whereas the univariate effect of *Z* was almost completely accounted for by a weaker effect of *U.* Thus, while the magnitude of the pupil response was sensitive to both *Z* components, *CPP* was clearly the stronger contributor. Shaded area, s.e.m., significance bars, p<0.05 (two-tailed cluster-based permutation test).

